# Codon decoding by split-tRNA: revisiting the tRNA selection mechanism

**DOI:** 10.1101/2024.11.24.624523

**Authors:** Sergey Mureev, Yue Wu, Zhenling Cui, Kirill Alexandrov

## Abstract

The translation machinery is required to process all codon triplets without exception while maintaining high speed and accuracy, despite orders-of-magnitude differences in cognate pairing stability. For stability-based selection to be efficient, the range of pairing stabilities must be narrowed by raising the lower bound and lowering the upper bound. The constrained structure and intramolecular cooperativity of tRNA complicate understanding of how it modulates codon–anticodon stability and whether it affects selection kinetics beyond codon recognition. To address these questions, we engineered functional split-tRNAs bearing a dangling anticodon in place of the anticodon loop. Our results demonstrate that split-tRNA supports in vitro translation nearly as efficiently as intact synthetic tRNA, challenging the notion that tRNA strain is essential for triggering GTP hydrolysis in response to codon recognition. Using split-tRNA architecture, we found that codon–anticodon stability is likely modulated by the dipole moments of adjacent nucleobases. Our kinetic modeling aligns with a conformational selection mechanism, where the decoding site fluctuates between open and closed states, and the correct codon–anticodon minihelix acts as an allosteric effector that permits its spontaneous closure and stabilizes the closed state. Overall, our data challenge the notion that tRNA is an active player in the selection process.

## Main Text

A trade-off between speed and accuracy is inherent to most biological processes (*1*, *2*). This is particularly true for translation, which must operate sufficiently fast to self-replicate and maintain cellular proteostasis (*3*). As a consequence, the ribosome must have evolved to maximize discrimination between cognate and near-cognate codon-anticodon interactions within the limited time window of tRNA selection. This process consists of an initial assessment of codon-anticodon interaction, known as initial selection, followed by a proofreading step.

Initial selection involves the codon-independent binding of an aminoacyl-tRNA (aa-tRNA), in complex with a GTP-loaded elongation factor (EF-Tu), to the small ribosomal subunit, followed by codon-anticodon recognition. In the next step, the closure of the ribosomal decoding site around the codon-anticodon minihelix triggers the GTPase activity of EF-Tu.

In the closed complex, the decoding site scans the first and second codon positions of the minihelix by interacting with the backbone of its minor groove (*4*). Notably, this interaction creates a new molecular interface that imposes energetic costs distinct from those of codon-anticodon interaction. The combined energetic effects of both interactions can, in principle, confer up to 10,000-fold selectivity for cognate aa-tRNAs over near-cognate variants mismatched at the first or second codon position (*5*, *6*). However, it is widely recognized that, for an optimal speed–accuracy trade-off (*7*–*9*), the codon recognition step is under kinetic control, limiting its capacity to fully exploit stability differences between cognate and near-cognate codon–anticodon pairs. There is, however, one caveat: sterically neutral mismatches at the third codon position may evade surveillance by the decoding site. Consequently, for many tRNAs recognizing codons from the split codon families, such mismatches must be discriminated either via reduced codon-anticodon complex stability or result in elevated error rates during initial selection (*10*–*15*) (table S1).

Despite decades of structural, biochemical, and biophysical analysis, there is no consensus on the mechanism by which the translation machinery accommodates the above constraints. In particular, the extent to which differences in thermodynamic stability between cognate and near-cognate codon-anticodon complexes contribute to selectivity, and whether tRNA affects the selection kinetics beyond codon recognition, remain incompletely resolved (*5*, *16*).

These uncertainties stem, in part, from the scarcity of experimental data on the intrinsic differences in dissociation rates between cognate and near-cognate tRNAs bound to codon in the context of an open decoding site (*17*, *18*). The widely suggested ∼1000-fold disparity is likely attributable to the selective stabilization of cognate tRNA by decoding site closure (*2*, *4*, *8*, *17*, *19*–*22*). Therefore, how tRNA dissociation from its codon and GTPase activation rates compare remains an open question. This is further compounded by uncertainty over the mechanism that couples the codon-anticodon identity to GTPase activation, particularly for split codon family readers. Conceptually, the driver for the coupling can be either the cognate pairing that promotes activation via the forward rate or the near-cognate pairing that inhibits it via the reverse rate. The first model aligns with the classical induced fit framework of tRNA selection, in which codon-anticodon binding energy is partially invested in promoting the assembly of a catalytically competent state for GTP hydrolysis (*7*, *8*, *23*). Given the obligatory distortion of the tRNA during the codon recognition (*4*, *24*), it has been proposed that this may generate strain within the molecule, which is crucial for triggering GTP hydrolysis (*25*–*27*). In support of this view, a mismatch-induced deviation in anticodon loop orientation may be amplified along the tRNA, resulting in misalignment of the acceptor end at the GTPase activation center (*28*, *29*). Alternatively, a stable codon-anticodon complex may accelerate decoding site closure by reducing the entropic component of the corresponding activation barrier (*7*, *17*).

In contrast, according to the conformational selection model, discrimination of sterically neutral third-position mismatches may be facilitated by their accelerated dissociation prior to spontaneous decoding site closure or by destabilization of the closed state.

Determining whether the stability of the codon-anticodon pairing affects the balance between tRNA dissociation and GTPase activation rates is essential for identifying the relevant mechanism.

The ribosome must process all 61 triplets, which exhibit a ∼10,000-fold difference in complementary duplex stabilities (*30*) (table S1). Specifically, codons with G or C in the first position, which form stable to moderately stable codon-anticodon duplexes, are primarily found in unsplit codon families. In contrast, codons with A or U in the first position, which form duplexes of moderate to weak stability, are predominantly associated with split codon families (*30*) (table S1). This partitioning may reflect evolutionary pressure to facilitate rejection of third-position mismatches in split codon families (*31*). A ∼10,000-fold range in stability among cognate duplexes would be expected to exceed the affinity difference between cognate and near-cognate substrates. If uncompensated, this disparity could undermine the efficiency of stability-based discrimination between them. The Extended Anticodon Hypothesis proposes that the anticodon loop adjusts the strength of codon-anticodon interactions by varying nucleotide identities in its flanks (*32*). Understanding the molecular basis of such adjustment would clarify how the anticodon loop determinants narrow the stability range of codon-anticodon duplexes, and whether codon-anticodon complexes formed by readers of split and unsplit codon families are segregated into distinct stability classes. These findings would help clarify the balance between thermodynamic and kinetic control during the codon recognition step.

However, predicting codon-anticodon complex stability is challenging, because multiple factors contribute to the properties of the constrained anticodon loop. Although the impact of a weak stacking context at N38 or the base complementarity between N32 and N38 in reducing stability can be rationalized (*22*, *33*–*36*), the contributions of other determinants remain unclear. For example, it is unclear why an optimal loop context is likely established when A occupies positions 37 and 38, whereas G, despite its favourable stacking properties, is rarely found at position 38 and tends to destabilize the complex when present at position 37 (*37*).

To address the questions raised above, we engineered a functional split-tRNA with a discontinuous anticodon arm, allowing modulation of codon-anticodon complex stability without altering the geometry of the codon-anticodon minihelix. Our findings indicate that the anticodon loop is not essential for translation and challenge the notion that the strain within tRNA is crucial for triggering GTP hydrolysis. This new decoding architecture, combined with a novel codon suppression assay targeting an individual codon within the ORF, allowed us to evaluate the decoding efficiency of all N37N38 split-tRNA variants. We found that codon-anticodon complex stability is primarily influenced by the dipole moment of adjacent nucleobases. These data also allowed us to model the impact of compromised stability in the N36-N39 segment on the kinetic parameters of tRNA selection steps. Based on our analysis and prior research, we propose that the stability of cognate codon-anticodon pairing does not influence the rate of GTP hydrolysis, and that the codon-anticodon minihelix functions primarily as an allosteric effector that stabilizes the closed conformation of the decoding site, consistent with a conformational selection mechanism.

## Results

### Rational for anticodon loop splitting

RNA maximizes base stacking by ordering intraloop nucleotides on the 3’ side of the stem and using coaxial stacking to prevent exposure of bases and base-pair planes to the solvent (*38*, *39*). This results in the formation of a quasi-continuous helix-like structure, propagating from the codon-anticodon minihelix to the anticodon stem via the 3’ anticodon flank (nucleotides 37, 38) (*40*, *41*) (Fig. 1, A and D). Nucleotides 32 and 33 in the 5’ anticodon flank mediate loop closure and do not participate in continuous stacking interactions or contact the ribosomal decoding site (*42*) (Fig. 1D). We hypothesized that the stacking column involving the 3’ anticodon flank makes the dominant contribution to codon-anticodon complex stability. This renders the 5’ anticodon flank functionally dispensable and potentially enables its deletion, resulting in a discontinuous sequence (Fig. 1B). The loss of rigidity in the discontinuous anticodon arm is expected to impose a significant entropic penalty on the stability of the codon-anticodon complex. This, however, may not be detrimental to the functionality of the split system, as the stability of cognate codon-anticodon duplexes naturally varies by nearly 10,000-fold (table S1). Hence, the entropic penalty could potentially be compensated by combining a stable, GC-rich codon-anticodon duplex with an optimal 3’-flanking sequence. The absence of modifications within the anticodon terminus is not expected to impair split-tRNA functionality, as they contribute to stabilization of weak codon–anticodon interactions and are not essential for translation (*43*–*45*).

**Fig. 1.**
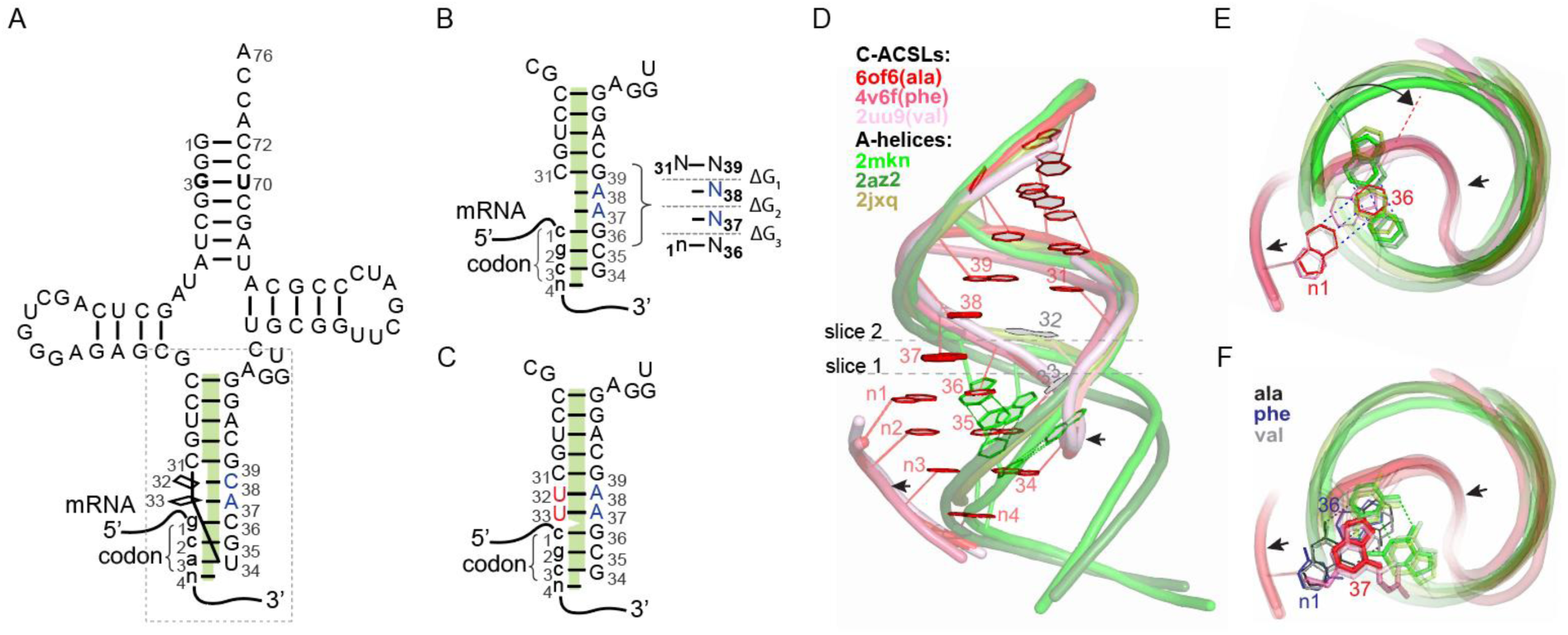
Structural basis of codon-anticodon interactions and implications for constructing functional split-tRNA. (**A**) Schematic of wild-type tRNA^Ala^UGC aligned with its codon, highlighting a quasi-continuous stacking column formed by the codon-anticodon minihelix, the N37N38 flank, and the anticodon stem (green). The dashed box outlines the codon-anticodon stem-loop complex shown in panels (B) and (C). (**B**) Discontinuous anticodon arm of chimeric split-tRNA^Ala^GCG, with a deleted N32N33 flank, in complex with the arginine codon CGC. Stacking interfaces within the N36-N39 segment and corresponding free energy changes (ΔG_1_-ΔG_3_) are shown in the inset on the right. (**C**) As in (B), but with an extended anticodon stem (red) including the A37A38 flank. (**D**) Side view of high-resolution structures showing three codon-anticodon stem-loop complexes (in shades of red) superimposed on canonical A-form RNA helices (in shades of green). Alignment pattern and sequence information are provided in table S2. Molecular backbones are shown as worms, with nucleobases as wires (PDB entries: 6of6 [for Ala C-ACSL] and 2mkn). The inset displays the colour coding for the PDB entries. An arrow marks the protrusion of codon-anticodon base pairs beyond A-helical boundaries. (**E**) Top-down view corresponding to optical slice 1 from (D), illustrating the positions of n1:N36 and the overlapping base pair in A-form helices, as well as the overtwist angle (arc arrow). (**F**) As in (E), but for optical slice 2. The n1:N36 base pairs are color-coded according to their assignment to specific C-ACSL (left inset). The figure shows that in the intact, overwound anticodon loop, N37 serves as a stacking platform for both n1 and N36 of the first C-AC base pair, whereas the A-form helix counterpart of N37 overlaps only with the proximal part of N36.

Splitting the anticodon loop should not interfere with the ability of the elongation factor (EF-Tu) to bind and deliver such tRNA to the ribosome, given the remote position of the tRNA:EF-Tu binding interface.

With this in mind, we designed a split-tRNA based on tRNA^Ala^, as its anticodon arm is not involved in the aminoacylation process. We introduced the arginine anticodon GCG, flanked by the commonly occuring A37A38 sequence (*37*, *40*, *41*), at the dangling end (Fig. 1B).

### Establishing a fluorescence rescue assay for split-tRNA functionality analysis

To assess the functionality of split-tRNA we developed a fluorescence rescue assay that monitors decoding of a specific codon by a specific tRNA. The assay is based on the translation of a mutant, non-fluorescent GFP in which alanine 226 is replaced with arginine. The effect of the mutation can be reversed by supplementing an *in vitro* translation reaction with a chimeric tRNA featuring an arginine anticodon but charged with a function-restoring wild type amino acid, such as alanine (Fig. 2A). The six-fold degeneracy of arginine codons enables construction of a codon-biased GFP ORF, in which an arginine codon from one codon family is used at position 226, while codons from a distinct family encode the remaining native arginine residues (Fig. 2A). The functionality of the assay was confirmed using a coupled *E. coli* S30 *in vitro* transcription/translation system, programmed with DNA templates containing the A226R mutant GFP-coding ORF (*46*).

**Fig. 2.**
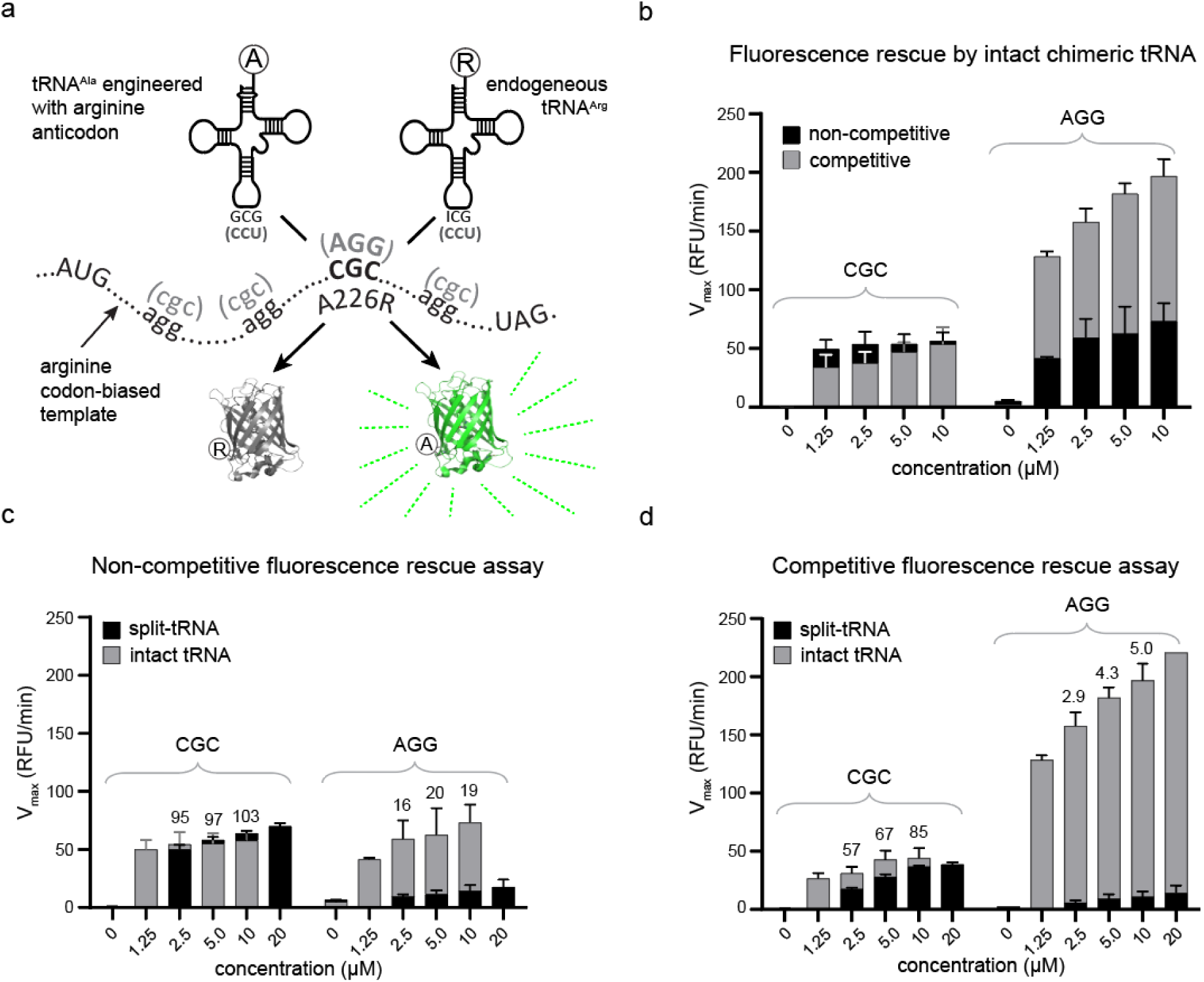
Assessment of split-tRNA functionality using the fluorescence rescue assay. (**A**) A schematic of the assay. The arginine codon at position 226 of a codon-biased template encodes the non-fluorescent GFP(A226R) mutant, which can be decoded either by a chimeric tRNA^Ala^ that incorporates alanine and rescues fluorescence or by endogenous tRNA^Arg^, resulting in non-fluorescent GFP. The ‘cloverleaf’ structures of chimeric tRNA^Ala^ and native tRNA^Arg^ICG are shown above the template. Alternative codon/anticodon combination is indicated in parentheses. (**B**) A bar chart showing maximal fluorescence accumulation rates (V_max_) in *in vitro* translation reactions programmed with GFP-A226R templates containing either a CGC or AGG codon at the mutation site (indicated above the bracket), and supplemented with the corresponding chimeric tRNA^Ala^. Reactions were performed under competitive conditions (grey bars) using S30 *E. coli* extract or under non-competitive conditions (black bars) using fractionated extracts depleted of endogenous tRNA^Arg^ (Fig. S1). Error bars represent standard deviations from 5 pseudo-replicates for CGC and 3 technical replicates for AGG. (**C**) Translation efficiency of split-tRNAs with GCGaa or CCUaa dangling ends (Fig. 1B) compared to intact chimeric tRNAs under non-competitive translation conditions. Values above the bars indicate the percentage of V_max_ for split-tRNAs relative to intact tRNA^Ala^. Error bars represent standard deviations from 3 independent translation experiments. (**D**) As in (C), but under competitive conditions. The difference in apparent decoding efficiencies of the intact chimeric tRNAs for the AGG and CGC templates is explained by the disproportionate abundances of their endogenous competitors.

*In vitro* translation reactions supplemented with purified chimeric tRNA^Ala^ carrying GCG or CCU anticodons showed a dose-dependent increase in fluorescence yield (Fig. 2B). Under these conditions, hereafter referred to as ‘competitive’, the chimeric tRNA competes with endogenous arginine isoacceptor(s) for the codon of interest. To estimate the decoding efficiency of chimeric tRNA, we compared fluorescence yields under competitive and non-competitive conditions. The non-competitive condition was established by supplementing a translation reaction, based on fractionated S30 lysate depleted of total tRNA, with a tRNA complement specifically depleted of endogenous competitors (*45*) (Fig. S1). As shown in Fig. 2B, the CGC template yielded higher fluorescence under non-competitive conditions, whereas AGG generated a stronger signal under competitive conditions, contrary to expectations. This is explained by the combined effect of higher translation efficiency in non-fractionated lysate (competitive conditions) and the much greater abundance of endogenous tRNA^Arg^ICG compared to tRNA^Arg^CCU (*47*).

To assess the reading efficiency of codons other than arginine, we also developed a generalizable ‘two-codon’ assay that monitors fluorescence accumulation in response to complementation of two consecutive codons of interest by chimeric or wild-type tRNAs (Fig. S2, A and B). This assay can also reveal whether the translation system can tolerate a split-tRNA occupying both the peptidyl and aminoacyl sites of the ribosome.

### Construction and functional characterization of tRNA with a split anticodon loop

The split-tRNA based on the tRNA^Ala^UCG isoacceptor was constructed by annealing synthetic RNA fragments spanning nucleotides 1-31 and 34-76 (Fig. 1B and Fig. S2C). We evaluated the codon suppression efficiency of split- and intact tRNAs featuring GCG and CCU anticodons in a fluorescence rescue assay under competitive and non-competitive reaction conditions. Surprisingly, fluorescence yields of reactions containing split-tRNA^Ala^GCG were comparable to those of reactions primed by intact chimeric tRNA^Ala^GCG (Fig. 2, C and D).

We confirmed that the observed translation was indeed mediated by the split-tRNA rather than by its enzymatically repaired derivative, by assessing the structural integrity of the split-tRNA during the translation reaction (Fig. S3, A to D). To ascertain the generalizability of the developed approach, we tested split-tRNA functionality in a eukaryotic translation system using *Leishmania tarentolae* extract (LTE) (*48*, *49*). We found that the split-tRNA exhibited ∼30% of the decoding efficiency of the intact chimeric tRNA under competitive conditions in the LTE system (Fig. S4).

We next assessed split-tRNA selection efficiency for codon-anticodon interactions involving nonisosteric G34:U3 or U34:G3 wobble pairs, as well as third-position mismatches. Decoding efficiency was reduced several-fold for U34:G3 relative to G34:U3, consistent with a previously estimated ∼0.5 kcal stability difference between them (*30*). Third-position mismatches led to a 10- to 100-fold decrease compared to the corresponding intact tRNAs (Fig. S5, A and B). Incorporation of a deoxynucleotide at position 37 also impaired decoding, likely due to increased flexibility of the deoxyribose (Fig. S5D). These findings suggest that the relative contribution of N3:N34 base-pair stacking to codon–anticodon complex stability is substantially greater in split-tRNAs than in intact tRNAs. This can be attributed to the higher entropic cost of accommodating a free dangling anticodon within the A-site.

We also tested whether extending the double-stranded region of the anticodon stem in the split-tRNA over the A37A38 flank would enhance codon-anticodon complex stability by introducing a U33:A37 base pair instead of relying on A37 alone for stacking with N1:N36 (Fig. 1C). Compared to the non-extended variant, the resulting 1-33/34-76 split-tRNA showed either unchanged or reduced decoding efficiency, depending on codon-anticodon interaction strength and the presence of 5’-ribose modifications at N34 (Fig. S3B and Fig. S5). This can be explained by an overtwisted loop configuration (Fig. 1E), which reduces stacking overlap between the N1:N36 base pair and N37 when N37 conforms to A-form geometry (Fig. 1F). This finding aligns with the previous observations that intraloop complementarity in intact tRNAs destabilizes the codon-anticodon complex (*33-36*). Accordingly, we observed that insertion of an extra nucleotide into the 3′ anticodon flank, relieving the overtwist, had minimal impact on decoding (Fig. S6).

In subsequent experiments, we found that the relative decoding efficiency of split-tRNA^Ala^CCU for the alternative arginine codon AGG was significantly lower than that of split-tRNA^Ala^GCG even under non-competitive conditions (Fig. 2C and D). This discrepancy can be attributed to the diminished stacking strength of uridine (U36) in the ‘cardinal’ position of split-tRNA^Ala^CCU compared to guanine (G36) in split-tRNA^Ala^GCG (*32*). Notably, the “two-codon” assay showed that split-tRNAs can decode adjacent codons, indicating that peptidyl- and aminoacyl-split-tRNAs can simultaneously occupy their respective sites on the ribosome (Fig. S2F). This assay also recapitulated the context dependence at the cardinal position, with U36-containing tRNAs showing reduced translation efficiency (Fig. S2F). In this weak context, the A3:U34 pair at the third codon position, compared to the more stable C3:G34 pair, resulted in negligible decoding activity (Fig. S5C). A similar trend was observed for both chimeric split-tRNAAlaGCU and split-tRNASerGCU, which also feature U36 in the cardinal position (Fig. S2, D to G).

Based on these observations, we concluded that in split-tRNA, the high entropy loss associated with the accommodating of the free dangling anticodon in the A-site makes the enthalpic contribution of stacking interactions critical for codon-anticodon complex stability. In contrast, intact tRNAs, with a conformationally constrained anticodon loop, experience a lower entropic penalty, resulting in a reduced reliance on stacking interactions.

### Comprehensive analysis of N37N38 combinations in split-tRNA

Removal of the intraloop constraint creates a novel system where the stability of the codon-anticodon complex is governed by stacking interactions within the N36-N39 segment (Fig. 1B). This enables analysis of the relationship between the nucleotide identity of this segment and the decoding efficiency of split-tRNA. To this end, we constructed 16 variants of split-tRNA^Ala^GCG corresponding to all N37N38 combinations (Fig. 1B), and tested them in a fluorescence rescue assay under competitive and non-competitive conditions. Control reactions supplemented with intact chimeric tRNA^Ala^GCG were used to normalize the fluorescence yields across conditions.

The observed reduction in fluorescence yield among the tested split-tRNAs ranged from negligible to pronounced upon transitioning from non-competitive to competitive reaction conditions (Fig. 3A). This variation likely reflects differences in dissociation rates among the N37N38 split-tRNA variants, which become more apparent in the presence of competing native tRNA^Arg^ICG. To further analyze these results, we plotted the ratios of relative fluorescence accumulation velocities in competitive versus non-competitive environments against the estimated stabilities of the N36-N39 segment (Fig. 3A inset). The stabilities were calculated as the exponential of the sum of free energy changes for the formation of N1-N36/N37, N38/N31-N39, and N37/N38 stacks (Fig. 1B inset, table S3).

**Fig. 3.**
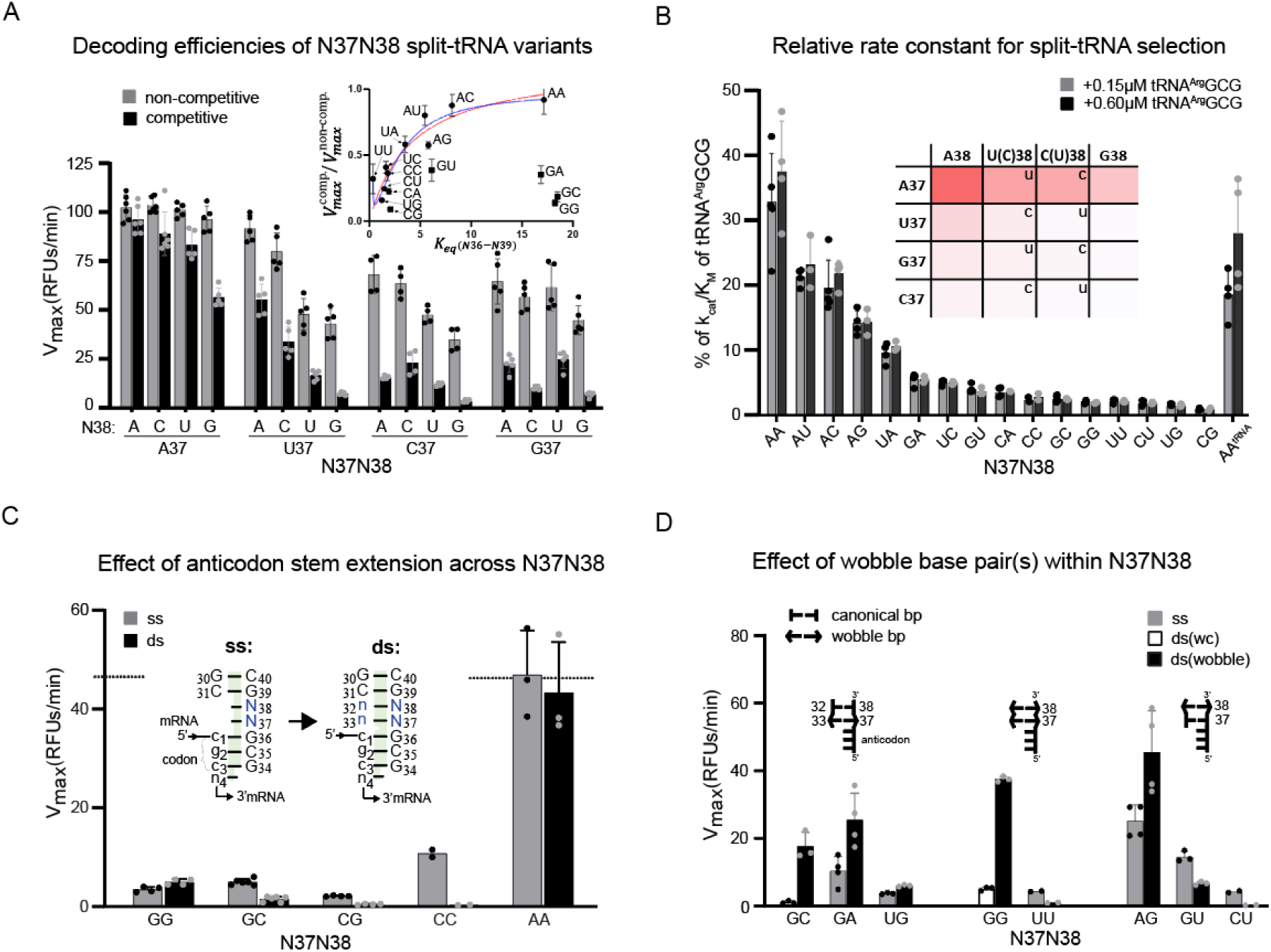
Impact of N37N38 identity on split-tRNA^Ala^GCG decoding efficiency. (A) A bar chart showing the percentage of maximal fluorescence accumulation rate (V_max_) for various N37N38 split-tRNA variants relative to intact chimeric tRNA^Ala^GCG, under competitive and non-competitive translation conditions. Data are shown as mean ± standard error from five independent experiments. The inset shows decoding efficiency plotted against the estimated stability of the N36-N39 segment. Data points (black circles) are fitted to quadratic (blue) or hyperbolic (red) equations (Eq. S11), with outliers (black squares) excluded. The fit is shown as a solid line. (B) A plot of normalized k_cat_/K_m_ for various N37N38 split-tRNA variants relative to that for intact tRNA^Arg^GCG. Bars represent average values from experiments using S30 lysate depleted of endogenous tRNA^Arg^ICG and reconstituted with competing synthetic tRNA^Arg^GCG at 0.15 µM or 0.6 µM. Error bars show standard deviations from three and four independent experiments, respectively. The inset presents a heatmap summarizing nucleobase preferences at positions 37 and 38 for optimal decoding efficiency. (**C**) Comparison of decoding efficiencies of split-tRNA variants with single- or double-stranded N37N38 segments, as shown in the inset. The dashed line indicates the V_max_ value for intact tRNA. Data are based on two to six independent measurements. (**D**) The efficiencies of split-tRNAs with wobble base pairs are compared to those with single-stranded or canonical double-stranded N37N38 configurations. Schematic representations of the terminal N34-N38 section of the split anticodon arm are shown above the respective bars for clarity. Data are based on two to four independent measurements.

Stacking energies for the first two segments were estimated from Dangling End stabilities, while those for the N37/N38 interface were derived from reported experimental measurements or molecular dynamics simulations of dinucleotide monophosphates (tables S4 to S6). The plot revealed only a partial correlation between estimated stabilities and decoding efficiency (Fig. 3A inset). Notably, despite the high stacking stabilization energy associated with G-containing interfaces, split-tRNA variants with G37 exhibited reduced decoding efficiency.

The ratio of fluorescence accumulation velocities (*V*_*max*_) under competitive versus non-competitive reaction conditions reflects the likelihood of productive selection of split-tRNA^Ala^GCG on the CGC codon in the presence of competing tRNA^Arg^ (Supplementary text 1). This ratio can be expressed using the selection rates for split-tRNA (*Rate*^*Ala*^) and competing tRNA^Arg^ (*Rate*^*Arg*^), as shown in Eq. 1:

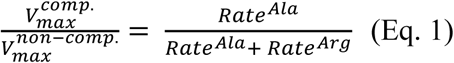

The tRNA selection rate can be defined as the product of the GTP hydrolysis rate by EF-Tu and the probability of successful passage through the proofreading step. Since the decoding site likely remains closed around the canonical codon-anticodon minihelix during accommodation step (*7*, *50*), we assumed these probabilities to be comparable for different substrates, regardless of their anticodon context, provided that cognate codon-anticodon pairing occurs (Supplementary text 1). Therefore, the rates in Eq. 1 can be expressed as the products of the apparent second-order rate constants 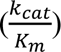 of GTP hydrolysis by EF-Tu and the concentrations of the respective ternary complexes (TC), as detailed in Supplementary Text 1 (Eqs. S1 to S5), resulting in Eq. 2:

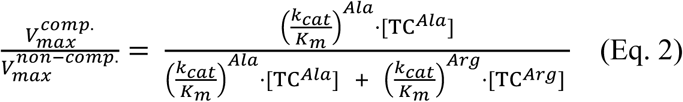

Using Eq. 2 and the concentrations of ternary complexes, we derived second-order rate constants for the selection of N37N38 split-tRNA variants, normalized to the intact tRNA competitor 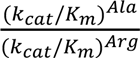 (Supplementary text 1, eq. S6). We found the concentrations of aminoacylated tRNAs to be ∼ 4 µM during the log phase of product accumulation (Fig. S7 and Fig. S8), and used this value as a proxy for estimating ternary complex concentrations. Instead of measuring the concentration of competing native tRNA^Arg^ICG (TC^*Arg*^) in the S30 lysate, we introduced T7-transcribed tRNA^Arg^GCG into a tRNA-depleted lysate to recreate competitive reaction conditions in the same translation environment. Since the introduced tRNA^Arg^GCG effectively competed with split-tRNA at submicromolar concentrations, we presumed that the former was fully aminoacylated. Performing both competitive and non-competitive translation reactions in the same depleted lysate eliminated the need for fluorescence normalization and enabled direct calculation of k_cat_/K_m_ values for all N37N38 variants, normalized to the k_cat_/K_m_ of the competing tRNA (supplementary text 1, eq. S6). The similar k_cat_/K_m_ values obtained at two different concentrations of competing tRNA^Arg^GCG support the robustness of the assay and its suitability for quantitative analysis of decoding efficiency (Fig. 3B). The corresponding plot shows that the presence of an adenine moiety at either position 37 or 38 resulted in the highest decoding efficiency within each set, whereas the A37A38 combination exhibited a synergistic enhancement (Fig. 3B inset).

Compared to variants featuring GCG anticodon, those with CCU anticodon showed similar N37N38 preferences for optimal decoding of the AGG codon, although their normalized k_cat_/K_m_ values were more than an order of magnitude lower (Fig. S9A). In addition to the weak stacking context of U36 in the cardinal anticodon position, this difference may reflect the unusually high decoding efficiency of native tRNA^Arg^CCU, which may represent an adaptation to mitigate high rates of miscoding at AGG codons (Fig. S5C inset). Similar to the AGG codon, much lower decoding efficiency for AGC was observed, particularly for G37-containing split-tRNAs based on tRNA^Ser^GCU (Fig. S9B).

To determine the cause of the low decoding efficiencies observed for G37/G38-containing split-tRNA variants, we first ruled out extrinsic factors, such as N37N38-specific degradation of the anticodon termini or differences in aminoacylation levels (supplementary text 2, Fig. S7 and Fig. S8).

Intrinsic factors that may contribute to the low decoding efficiency of G37/G38-containing split-tRNA variants include the distinct energetic effects of specific N37 and N38 combinations, formation of inter- or intramolecular secondary structures by the protruding N34-N38 strand, and N37N38-dependent coordination of Mg^2+^ (*51*). Experimental testing ruled out secondary structure formation as a major factor contributing to the reduced decoding efficiency of G/C-rich N37N38 split-tRNA variants (supplementary text 2, Fig. S10). Additionally, N37N38 split-tRNA variants generally exhibited similar relative decoding efficiencies for the CGC codon across different Mg^2+^ concentrations, supporting the notion that Mg^2+^ primarily facilitates the initial, codon-independent stage of ternary complex binding to the ribosome (*5*, *6*) (supplementary text 2, Fig. S11).

Next, we examined how thermodynamic factors may influence the observed decoding efficiency patterns. Specifically, we examined whether N37N38-dependent variation in the activation energy barrier for transitioning the N34–N38 region from its free to codon-bound state could account for differences in the forward rate of codon–anticodon complex formation. To address this, we standardized the free energies of the unbound states by extending the double helix of anticodon stem through the N37N38 segment (Fig. 1C and Fig. 3C, inset). The resulting variants showed either unchanged or substantially reduced decoding efficiency (Fig. 3C), suggesting that variation in the forward rate is unlikely to account for the observed differences in decoding efficiency.

Remarkably, involvement of G37 in wobble pairing with U33 enhanced split-tRNA performance by more than 10-fold compared to non-extended or canonically extended variants (Fig. 3D). This wobble geometry likely stabilizes G37/38 in a conformation that introduces new stacking edges, improving alignment within the N36-N39 stem while reducing entropy loss (supplementary text 3, Fig. S12). In contrast, the inverse wobble geometries present in U37C38 or U37U38 variants did not enhance decoding efficiency, likely due to the non-isosteric nature of G:U and U:G base pairs (Fig. 3D).

The orientations of adjacent nucleotides are governed by the balance between repulsion of their individual dipoles and the solvation energy of their net dipole (*52*). Polarizability, a critical factor for effective stacking, follows the order G > A > C > U (*53*, *54*), whereas the dipole moment of individual nucleobases decreases in the order G ∼ C > U > A (*55*–*57*). Consequently, positioning G37 between the C1:G36 base pair and G38 or C38 is expected to result in increased repulsion between the large dipole moments of the three residues in the N36-N38 stack. This effect may be pronounced in the context of a dangling end. Unlike other nucleobases, A lacks strong electronegative ring substituents, resulting in the smallest dipole moment. Simultaneously, its polarizability is comparable to that of G (*53*, *54*, *58*). This unique combination allows A-containing N37N38 variants to adopt a broader range of orientations, providing optimal stacking with neighbouring nucleobases regardless of their identity. The ordering of nucleobases at positions 37 and 38 favours smaller dipole moments over polarizability, with decoding efficiency increasing in the order A >> U(C) > G for the given N38 or N37, respectively (Fig. 3B inset).

### *S*tability of codon-anticodon complexes and its effect on tRNA selection kinetics

As noted in the introduction, the translation system must accommodate several constraints to enable rapid discrimination between cognate and near-cognate substrates (Supplementary Text 4). This is particularly challenging when the substrates differ by sterically neutral mismatches at the third codon position, where selectivity must rely on stability differences between the respective codon-anticodon complexes. The key question is how these differences are leveraged in the framework of the classical induced fit or conformational selection models (Supplementary Text 4). According to the induced fit model, selectivity between correct and incorrect aa-tRNA substrates appears to arise from differences in the fluctuation freedom of their codon-bound states, which modulate the entropic component of the activation barrier for the subsequent conformational step (k₃; see Scheme 1 and Fig. 4A). In conformational selection, stability differences manifest either through enhanced dissociation of less stably bound near-cognate ternary complex (k_-2_; see Scheme 1 and Fig. 4A) or, if codon recognition is kinetically controlled, through more rapid reversal of the locked decoding site configuration (k_-3_; see Scheme 1 and Fig. 4A).

**Fig. 4.**
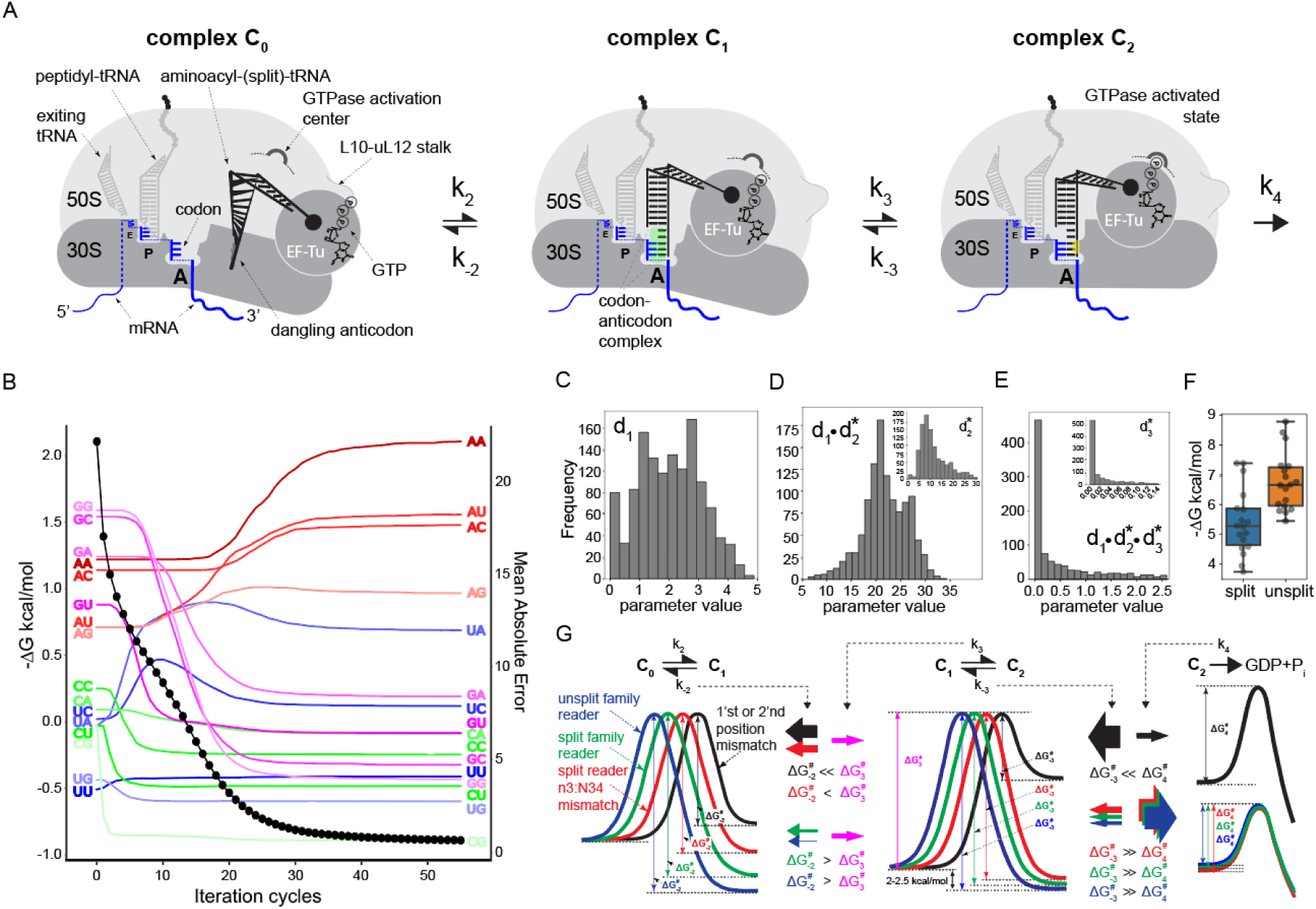
Influence of codon-anticodon complex stability on split-tRNA selection kinetics. (**A**) Schematic representation of tRNA selection on the ribosome, illustrating changes in interaction interfaces with the ternary complex at different selection steps (see Scheme 1). In complex C_0_, mRNA (‘cofactor’) presents the codon triplet as a ‘recognition surface’ for the anticodon (‘ligand’s binding site’). The stability of the interaction interface (green) is assessed in complex C_1_. At the C_1_-to-C_2_ transition, the codon-anticodon minihelix acts as an allosteric effector, providing a new recognition surface for the decoding site’s monitoring bases. The new interaction interface (yellow) in complex C_2_ influences the assembly of the transition state for GTP hydrolysis. (**B**) Evolution of stability terms from fitting Eq. 5 to the regression model (Fig. S13). Traces are averaged from 1000 simulations. The X axis shows fitting iterations; the left Y axis denotes N36-N39 segment stabilities (kcal/mol), and the right Y axis shows the mean absolute error of the fit at each iteration (black circles). (**C**) Frequency distribution of predicted d_1_ values (1000 simulations). Optimal bin count determined by the Freedman-Diaconis rule (see Git repository). (D) As in (C) but for regression coefficient d_1_d_2_; inset shows the distribution of d_2_. (E) As in (D) but for regression coefficient d_1_d_2_d_3_; inset shows the distribution of d_3_. (F) Distribution of estimated codon-anticodon complex stabilities for readers of split and unsplit codon families (table S1). Boxes show the interquartile range (IQR); horizontal line shows median stabilities. Outliers (tRNA^Trp^CCA and tRNA^Leu^UAG) excluded for clarity. (**G**) Free energy profiles across tRNA selection steps, as indicated above each profile.

The split-tRNA framework provides an experimental platform for manipulating codon-anticodon complex stability via different N37N38 combinations without altering the geometric component of selection.

Scheme 1 illustrates the four-step kinetic model of initial selection (*5*), beginning with codon-independent binding of the ternary complex to the ribosome to form complex C_0_, followed by codon recognition (complex C_1_). This is followed by a conformational step, involving closure of the decoding site around the canonical codon-anticodon minihelix, yielding the pre-catalytic complex C_2_ (Fig. 4A):

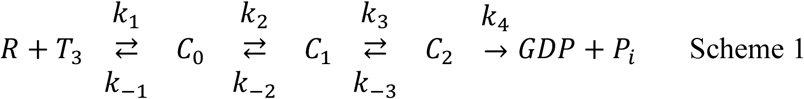

The above scheme is described by Eq. 3, which relates the apparent second-order rate constant 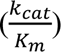 for the GTPase reaction to the kinetic parameters of the selection steps (*5*):

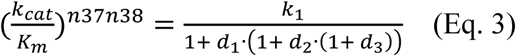

In this equation, k_1_ is the codon-independent rate constant for the association of the ternary complex with the ribosome, which corresponds to the maximum theoretical rate of tRNA selection. This rate is progressively reduced by rejection probabilities, quantified as discard parameters d_1_, d_2_, and d_3_, defined by the ratios 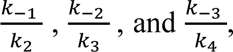 respectively (Scheme 1). The magnitude of these parameters relative to unity indicates whether the given selection step proceeds under kinetic or thermodynamic control.

According to Eq. 3, while d_1_ is independent of codon-anticodon interaction, the stability of the codon-anticodon complex may affect k_cat/_K_m_ through d_2_, d_3_, or both discard parameters (Supplementary text 5). Thus, estimating these discard parameters by fitting experimental data to Eq. 3 may reveal which stages of tRNA selection are sensitive to variations in codon-anticodon complex stability.

To this end, our objective was to transform Eq. 3 into a form that correlates k_cat/_K_m_ for GTP hydrolysis by EF-Tu with the contribution of the N36–N39 segment to codon–anticodon complex stability. This contribution, together with that of the three base pairs and two stacking interfaces of the codon–anticodon minihelix, defines overall complex stability and is inherently reflected in the discard parameters via the forward and reverse rate constants specific to each N37N38 split-tRNA variant. To incorporate individual N36-N39 stabilities as independent variables in Eq. 3, we introduced universal surrogate discard parameters (d_2_* and d_3_*), corresponding to a hypothetical split-tRNA* lacking stacking interactions in the N36-N39 segment. The N36–N39 stability term then enters Eq. 3 as an adjustment factor to these surrogate parameters (Supplementary text 5), yielding Eq. 4:

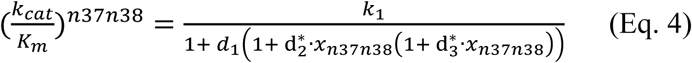

In this equation *x*_*n*37*n*38_ is defined as 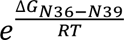 for each N37N38 split-tRNA variant. This approach follows the nearest-neighbour model for nucleic acid stability, where overall stability is determined by the sum of individual stacking contributions.

As noted above, the Δ*G*_*N*36−*N*39_was initially calculated as the sum of free energy changes derived for individual stacking units (Fig. 1B), assuming their independence and optimal nucleotide overlaps (Fig. 1B; Tables S3-S6). However, in the constrained context of an accommodated codon-bound dangling end, Δ*G*_*N*36−*N*39_is influenced by N37N38-specific repulsion and attraction forces. Therefore, in our linear regression model, we approximated the actual Δ*G*_*N*36−*N*39_values by iteratively perturbing the initial estimates with randomly selected weights (Fig. S13).

Making several assumptions and using the experimental decoding efficiencies of each N37N38 split-tRNA variant along with estimated concentrations of competing ternary complexes, we transformed Eq. 4 into a quadratic polynomial form (Eq. 5) suitable for regression analysis (Supplementary text 5):

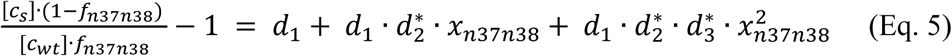

In this equation, [*c*_*s*_] and [*c*_*wt*_] represent the concentrations of aminoacylated N37N38 split-tRNA variants and competing synthetic tRNA^Arg^GCG, respectively. The parameter *f*_*n*37*n*38_ denotes the ratio of maximal fluorescence accumulation velocities 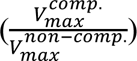 in reactions with or without competing tRNA^Arg^GCG for each N37N38 split-tRNA variant.

The overdetermined system of 16 equations for each N37N38 variant was subjected to 56 fitting iterations, each involving 100 weight perturbations (Fig. S13). We performed 1000 independent simulations, applying random weight perturbations in each case. Figure 4B illustrates the evolution of free energy terms for each variant across 56 iterations, averaged across all simulations. The improvement in the metric primarily results from reducing the stabilities of G37- or C37-containing split-tRNA variants relative to initial estimates. For most variants, excluding A37N38, the stability gain from stacking interactions within N36-N39 segment was moderate and often insufficient to compensate for the entropy loss associated with accommodating a free dangling end in the A-site (Fig. 4B). Simulations indicated that the A37A38 variant contributes more than 2 kcal/mol to codon-anticodon complex stability, whereas combinations involving G, C or U at positions 37 and 38 tend to destabilize the complex by 0.5-1 kcal/mol.

Because the intact anticodon loop likely adopts a conformation similar to that of the dangling end when aligned with a conformationally constrained codon triplet, the energy landscape of the native codon–anticodon complex is expected to be shaped by similar interactions involving dipole moments and polarizabilities within the N36–N39 segment. As a result, the ∼10,000-fold variation in stability observed among isolated codon–anticodon duplexes may be narrowed by approximately 100-fold in the full complex through changes in N37 or N38 identity, the presence of nucleotide modifications, and intraloop complementarity (Table S1). Narrowing this range, especially by increasing the stability of the weakest cognate interactions, may be important for split-codon readers to ensure that the least stable cognate complex exceeds the most stable mismatch by a sufficient margin.

As the regression metric improved, the model converged on coefficients corresponding to 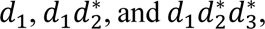 from which the individual discard parameters can be inferred. Figures 4C and 4D show the distributions for *d*_1_and 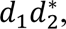 respectively, based on 1000 independent simulations. The d₁ parameter, which reflects the likelihood of aa-tRNA dissociation from the ribosome prior to codon recognition, exhibits a broad distribution with local maxima near 1 and 2.5, consistent with previous estimates (*59*) (Fig. 4C). The d_2_* parameter follows a right-skewed distribution, indicating that the dissociation rate (k_-2_) of the hypothetical split-tRNA from the codon exceeds the rate of the subsequent conformational step (k_3_) by roughly an order of magnitude (Fig. 4D). This value may exceed the fraction of unity reported for intact tRNA by nearly two orders of magnitude (*5*). This difference likely reflects a loss of several kilocalories per mole in stabilization energy due to disrupted stacking interactions within the N36–N39 segment.

The distribution of the 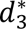 parameter (k_-3_/k_4_) is strongly skewed toward zero (Fig. 4E), suggesting that in a hypothetical split-tRNA, k_-3_ remains negligible relative to k_4_. This implies that the rate of GTP hydrolysis is effectively independent of codon-anticodon complex stability, provided that Watson–Crick geometry is preserved at the first two codon positions (Fig. 4A, Supplementary Text 4). Consequently, while error-inducing mismatches in unsplit-codon readers are rejected due to poor geometric fit regardless of their stability, split-codon readers are likely selected under partial thermodynamic control.

When codon–anticodon complex stabilities are estimated using model-predicted N36–N39 contributions (Table S1), split-codon readers rank lower in stability than readers of unsplit-codon families (Fig. 4G). As a result, their third-position mismatches are expected to fall within the lowest stability tier.

According to the conformational selection model, decoding site closure likely occurs when thermally driven fluctuations within the 30S subunit reach a resonance state that permits spontaneous crossing of the activation energy barrier. If this barrier is crossed on a timescale comparable to the average codon residence time of cognate split-codon readers, their third-position mismatches are expected to dissociate before closure is completed (Fig. 4G).

### Initiation of translation with split-tRNA-mRNA fusion

Our observation that split-tRNA supports translation as efficiently as its wild-type counterpart highlights its potential for engineering the translational machinery. A split-tRNA architecture offers the advantage of functioning in both *trans* and *cis* configurations with mRNA, which could be leveraged to manipulate translation initiation, elongation, and termination (Fig. 5A). To explore this, we tested whether split-tRNA could endow mRNA with a tertiary structure capable of mediating translation initiation. We hypothesized that a continuous A-helix formed by split-tRNA could mimic the codon-anticodon complex and serve as its surrogate, thereby driving translation initiation.

**Fig. 5.**
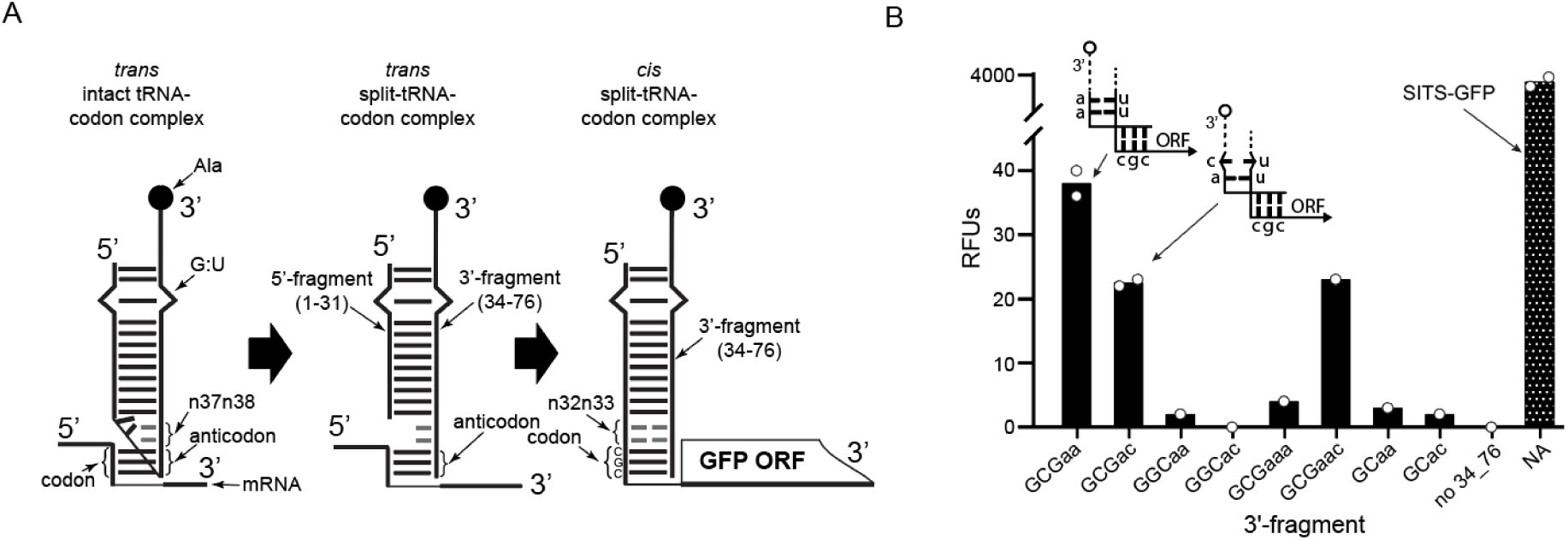
Split-tRNA-mRNA fusion as a mimic of the translation initiation complex. (**A**) Schematic showing the transition from *trans*-architecture codon-anticodon complexes (formed by intact and split-tRNAs) to *cis*-architecture in 5’-split-tRNA-mRNA fusion. The 1-33 5’-fragment is fused to a GFP ORF beginning with CGC codon and annealed with the 3’-fragment (34-76) of split-tRNA, mimicking the initiator tRNA in the ribosomal P-site. (**B**) Endpoint fluorescence counts for split-tRNA/mRNA fusion constructs translated in an *E. coli in vitro* translation system. The 1-33-CGC-eGFP-mRNA was annealed with various 3’-fragments, differing by anticodon triplets and nucleotide residues at position 38 (X-axis). Insets show stylized initiation complexes for the canonical A-U pair and C-U mismatch at positions N32-N38 (the 3’-fragments carrying Ala are marked as empty circles, with dashed lines indicating the remainder of the split-tRNA). SITS-GFP is a positive control containing SITS translation enhancer (*49*). Bars represent average values from two replicates.

To test this hypothesis, we fused a 1-33 split-tRNA fragment with U32U33 to the 5’ end of an eGFP-coding ORF prefaced with a CGC codon and evaluated the resulting mRNA in an *in vitro* translation reaction (Fig. 5, A and B). Although this mRNA by itself was translationally inactive, annealing with the complementary 34-76 fragment (‘GCGaa’) or the fragment yielding a U32•C38 mismatch (‘GCGac’) resulted in low but detectable GFP expression (Fig. 5B). Interestingly, the fragment with the C-extended flank (‘GCGaac’) also produced functional hybrid mRNA (Fig. 5B). Additional controls, including 34–76 RNA fragments that form mismatches in the codon–anticodon duplex and fragments with the anticodon truncated at position 34 (Fig. 5B), confirmed that the observed expression resulted from split-tRNA-mediated translation initiation. These observations strongly suggest that using an exogenous RNA fragment allowed us to reconstruct a structure sufficiently similar to the initiation complex. The low efficiency of this process is expected, as the native codon-anticodon complex is overwound relative to a classical A-helix. This may be addressed by adjusting the identity and number of codon- and/or anticodon-flanking bases, potentially incorporating wobble base pairs. In addition to the demonstrated translation initiation using 5’-mRNA-split-tRNA fusion, 3’-fusions may enable the co-translational formation of mRNA-protein fusions without chemical linkers. Furthermore, removing steric constraints within the anticodon loop should enhance the efficiency of codon reassignment for quadruplet codons and unnatural bases. Recent findings that split-tRNA structures can be assembled and maintained *in vivo* are promising, as they provide avenues for the deployment of split-tRNAs in living organisms (*60*).

## Discussion

Translation machinery must operate at a sufficient speed to sustain self-replication and maintain cellular proteostasis. It must process the full repertoire of codon triplets, whose affinities for their complementary anticodons vary by several orders of magnitude depending on G/C versus A/U content. Maintaining a high rate of tRNA selection therefore requires mechanism to minimize misincorporation without relying solely on codon-anticodon binding strength. Sequence-shape complementarity enables rapid geometric matching of codon-anticodon pairs. However, the geometry at the third codon position is not strictly monitored, likely to avoid delays associated with rejection of isoaccepting tRNA species that rely on wobble pairing. As a result, sterically neutral mismatches at the third position can lead to miscoding in split-codon families and must be rejected due to insufficient thermodynamic stability or discriminated through kinetic modulation involving tRNA and/or ribosomal component. Proofreading is unlikely to serve as the primary mechanism, given its potentially high energetic cost (*61*).

To enable effective stability-based discrimination, the stability of the weakest cognate codon-anticodon complex must exceed that of the most stable mismatch by a sufficient margin. The anticodon context is thought to balance codon-anticodon stabilities (*32*), although the underlying mechanisms remain incompletely understood. More broadly, intramolecular cooperativity within tRNA complicates efforts to resolve how its structural elements and internal dynamics contribute to the thermodynamic and kinetic components of substrate selection.

To create an experimental system in which codon-anticodon complex stability is uncoupled from intra-tRNA strain, we designed a split-tRNA with a dangling anticodon replacing the anticodon loop. Additionally, we developed a high-resolution fluorescence-rescue assay for codon-specific evaluation of tRNA functionality in an *in vitro* translation system.

We demonstrated that split-tRNAs can support translation with efficiency comparable to that of wild-type synthetic tRNA (Fig. 3B), challenging the notion that tRNA strain is essential for EF-Tu-mediated GTP hydrolysis. This finding is consistent with previous cryo-EM-informed molecular dynamics simulations and smFRET studies (*17*, *62*–*64*), which showed that both cognate and near-cognate tRNAs spontaneously sample distorted conformation (Supplementary text 6).

Using the fluorescence rescue assay, we observed a pronounced dependence of split-tRNA decoding efficiency on the identity of the ‘cardinal’ nucleotide at position 36, the G/C content of the codon-anticodon duplex, and the presence of wobble pairing at the third codon position. These findings support the idea that, in the split system, the relative contribution of stacking enthalpy to codon–anticodon complex stability increases to compensate for entropic losses associated with accommodating the dangling anticodon in the A-site.

The increased reliance on stacking suggests that codon–anticodon complex stability is strongly influenced by the identities of nucleobases at positions 37 and 38. To examine this, we tested split-tRNAs carrying all 16 possible N37N38 combinations and observed a decline in decoding activity in the following order: A37N38 > U37A38 > G37A38 > C37A38. Despite the greater stacking propensity of G and C, split-tRNA variants with these residues at position 37 were the least effective competitors (Fig. 3B). Further analysis revealed a preference for smaller dipole moments over greater polarizability at positions 37 and 38 with respect to decoding efficiency (Fig. 3C). We concluded that the large dipole moment of G may induce repulsion with adjacent residues, limiting conformational freedom and increasing the free energy of the codon-anticodon complex. In contrast, the favourable balance between dipole moment and polarizability allows A to achieve optimal stacking overlap with neighboring nucleotides.

Next, we utilized decoding efficiency data and approximate N36-N39 segment stabilities (table S3) for each split-tRNA variant to semi-empirically model how impaired stacking interactions within this segment, in the context of a hypothetical split-tRNA, influence the kinetic balance between successive selection steps (*5*) (Fig. S13). Simultaneously, the stabilities were perturbed using random weights to allow the model to sample and relax toward biologically relevant values during the regression process. The dominant values obtained for the discard parameters are consistent with prior estimates (*5*, *59*) and suggest enhanced dissociation of the hypothetical split-tRNA with disrupted N36-N39 stacking during codon recognition, without affecting the rate of GTP hydrolysis. This scenario is plausible if the activation barrier separating the open and closed states of the decoding site is sufficiently high, allowing GTP hydrolysis to occur before decoding site reopening, regardless of codon–anticodon duplex stability. These considerations point to functionally independent energetic contributions from the interaction interfaces within the complexes C1 and C2 (Fig. 4A).

Multiple lines of evidence support the model of spontaneous decoding site dynamics: the ribosome’s ability to hydrolyze GTP in the context of a binary EF-Tu–GTP complex (*65*); the relatively moderate impact from substitutions of atomic groups interacting with codon–anticodon minihelix (*67*); and the observation that the first step of decoding site rearrangement occurs on a similar timescale for cognate and near-cognate substrates (*6*, *17*, *18*). This may also explain why tRNA^Lys^UUU, with uridines capable of facing any codon base without significant steric clashes, exhibits one of the highest background miscoding frequency (*11*, *13*, *43*).

We used model-derived N36-N39 segment stabilities to estimate the overall stability of codon-anticodon complexes (Table S1). The analysis indicates that N37N38 combinations containing A37 increase the stability of weak codon–anticodon duplexes by more than 2 kcal/mol, whereas those with G37 slightly destabilize otherwise stable duplexes (Table S1). In the cellular context, the former are further stabilized by nucleotide modifications (*43*), while the latter can also be attenuated through intraloop base pairing between N32 and N38 (table S1).

Our estimations are consistent with prior studies (*30*) in placing split-codon readers at the lower end of codon–anticodon complex stabilities compared to those of unsplit-codon families (Fig. 4F). Importantly, mismatches involving split-codon readers would then fall into the lowest stability class. Thus, if the activation barrier for decoding site closure is comparable to the effective barrier for dissociation of split-codon readers from their codons, third-position mismatches formed by these tRNAs can be discriminated under partial thermodynamic control (Fig. 4G). While studies of conformational dynamics suggest that the rate-limiting step of decoding site closure occurs on the same timescale as tRNA^Phe^ dissociation from its codon, several aspects remain insufficiently resolved to allow unambiguous validation of the conformational selection model (17, 89). In particular, comprehensive data on the dissociation rates of cognate and near-cognate substrates from the codon prior to decoding site closure are lacking. Their indirect estimates from independent experimental observations suggest a difference exceeding one order of magnitude, which still appears too modest to account for the full extent of initial discrimination (*8*, *17*, *18*) (Supplementary text 4). Although modelling of discard parameters yields values consistent with previous estimates, experimental determination of split-tRNA affinities for A-site codon remains necessary to validate the model.

From a coarse-grained perspective, the conformational transition of the decoding site may appear as an induced-fit process when treated as a single-step event. In reality, the transition likely proceeds through a rapid initial substep followed by a slower, rate-limiting substep (*17*) (Supplementary text 4). In this scenario, the unused portion of the binding energy difference between cognate and near-cognate substrates during codon recognition may manifest in the accelerated reversal of the initial substep for near-cognates, thereby allowing the full thermodynamic difference in the stabilities of the respective codon–anticodon complexes to be exploited.

Based on our results, we conclude that the fidelity of the translational machinery primarily relies on conserved mechanisms governing RNA-RNA interactions and the intrinsic dynamics of tertiary RNA structures, possibly inherited from a pre-protein world. Our findings challenge the view that tRNA is an active player in the selection process. Instead, we propose that its information capacity is fully utilized in maintaining the universal core structure, modulating codon-anticodon interaction strength, and ensuring unambiguous aminoacylation determinants (*68*).

## Materials and Methods

### Plasmid construction

The constructs for T7-transcription of the eGFP-coding ORF prefaced with 5’-split-tRNA fragments and CGC codon were based on the pLTE-derived codon-biased plasmid 6064, in which all arginine codons of the GFP ORF were converted to AGG. To generate these constructs, the 5’-UTR coding sequence of 6064 was replaced by the respective synthetic fragments 1_33ttGCG and 1_33ctGCG, supplied as gBlocks by IDT (Integrated DNA Technologies) yielding plasmids 14 and 15, respectively. These fragments were subcloned via Gibson assembly method into the 6064 plasmid, opened via AatII and BglII.

### Preparation of ethanolamine-conjugated Sepharose for ion-exchange chromatography

The ethanolamine-conjugated sepharose resin for anion exchange chromatography was prepared as described previously (*69*) with minor modifications. Briefly, 8 g of water-washed epoxy-activated Sepharose 6B (GE Healthcare) was incubated with 40 ml of 1M ethanolamine at RT overnight with agitation. Following extensive washing to remove the traces of ethanolamine, the resulting matrix was stored either as a 20% (volume of sedimented resin after 1 min spinning at 1,000 g / total volume %) slurry in a storage buffer containing 100 mM NaOAc (pH 5.2), 0.25 mM EDTA and 2mM NaN_3_ or as a 50% slurry (v/v %) in a storage buffer containing 10 mM Hepes-KOH (pH7.5), 25 mM KCl, 10 mM NaCl, 0.1 mM EDTA, 1.1 mM Mg(OAc)_2_, 100 mM KOAc, and 2mM NaN_3_.

### Preparation of oligonucleotide-conjugated Sepharose for specific tRNA pulldown

The oligonucleotide-conjugated Sepharose was prepared as described previously (*69*). Briefly, 250 ul of N-Hydroxysuccinimidyl-Sepharose (NHS-Sepharose) suspension in isopropanol (Sigma-Aldrich #H8280), corresponding to 200 µl of settled gel with a conjugation capacity of 16-25 µmol/ml, was typically used for conjugation with 60 nmol of 3’-aminated oligonucleotide. This ratio corresponds to 50-100-fold molar excess of NHS groups over the oligonucleotide. The required amount of NHS-Sepharose slurry was transferred to a filter-bottom tube and washed with a total of 10-15 volumes of ice-cold 0.2 mM HCl in 3 steps, each followed by instant spinning at 500 g, with the final centrifugation step at 1,000 g for 1 min at 4°C. Following the last centrifugation step, the semi-dried resin was transferred to a 400 µl conjugation mixture containing 150 µM of 3’-aminated oligonucleotide, 50% DMSO, and 50 mM NaHCO_3_. The mixture was incubated overnight at RT with constant agitation at 1200 rpm. Following incubation, the resin was transferred to a filter-bottom tube and drained of the solution. To block the unreacted NHS groups, the semi-dried resin was incubated with 400 µl of a blocking buffer containing 0.5 M ethanolamine and 0.5M NaCl for 1 hour at RT with agitation, followed by alternating washes with 50 mM Tris-HCl (pH8.5) and a buffer containing 0.1 M NaOAc (pH 5.2) and 0.5 M NaCl. The oligonucleotide-conjugated Sepharose was finally washed with a storage buffer containing 10mM TrisHCl (pH 7.5), 100 mM NaCl, and 2 mM NaN3 and stored at +4°C as a 50% resin suspension (v/v). The tRNA binding capacity of the obtained resin was assumed to be 100 pmol/µl of 50% suspension.

### Synthesis and purification of mRNA templates

Runoff transcription was used to obtain the mRNAs corresponding to split-tRNA-fragment/ORF fusions. For this purpose, the respective plasmids 14 and 15 (supplementary data 2) were linearized with MluI digestion, 150 bp downstream of the stop codon triplet, directly followed by ethanol precipitation. The T7-transcription reaction contained 50 nM of the linearized plasmid template and was conducted in a 100 µl reaction volume as described in “Synthesis and purification of tRNAs and tRNA fragments”. The RNA transcripts were purified using the Monarch RNA Clean up Kit (#T2030L).

To assemble the 5’-split-tRNA-mRNA fusions, the purified mRNAs with the GFP ORF beginning with a CGC codon and prefaced with 5’-(1-33)-split-tRNA fragments were annealed with 3’-(34-76)-split-tRNA fragments in a 10 µl reaction. This reaction contained 5.5 µM of the mRNA fusion, 6 µM of the synthetic 34-76 fragment, 5 mM NaOAc (pH 5.0), and 0.5 mM EDTA. The annealing reaction and purification were conducted following the same procedure outlined for the assembly of split-tRNA.

### Synthesis and purification of tRNAs and tRNA fragments

The sequences of synthetic RNAs and oligonucleotides are summarized in supplementary data 2, respectively. tDNA templates containing a T7 promoter and a full tRNA-coding sequence for runoff transcription were assembled using a 3-step PCR reaction as previously described(*70*). Briefly, each PCR reaction comprised 1/10^th^ of 10x PCR buffer (containing 160 mM (NH_4_)_2_SO_4_ and 670 mM Tris-HCl pH8.8), 0.2 mM of each dNTP, 2 mM MgCl_2_, 2.5% DMSO and 1 u/µl of X7 DNA polymerase(*71*), along with varying concentrations of forward and reverse oligonucleotides. Specifically in Step 1, the forward oligonucleotide (F) containing the T7 promoter sequence was combined with the reverse oligonucleotide (R1) at concentrations of 1µM each. Overlap extension involved 5 cycles of 1 min denaturation at 98°C, 1 min annealing at 42°C, and 30 s elongation at 72°C. In Step 2, the PCR reaction was primed using 1/10^th^ of the crude Step 1 reaction relative to the volume of the Step 2 reaction. Amplification, using the same concentrations of the 5′-GCGG-extended 3′-G-ending T7-promoter oligonucleotide and R2-oligonucleotide spanning the 3′-part of the tRNA sequence, was conducted for 5 cycles with 1 min denaturation at 95°C, 1 min annealing at 42°C, and 30 s elongation at 72°C. In Step 3, the reaction mixture containing the same T7-promoter as the forward oligonucleotide and R3 as the reverse oligonucleotide, both at 7.5 μM concentrations, was primed with the crude reaction mixture from Step 2 diluted 1:50 into the Step 3 reaction. Amplification was carried out for 28 cycles, each consisting of 30 s denaturing, 30 s annealing, and 20 s elongation.

To assemble the PCR templates coding for the 3’-(34-76)-fragment of tRNA, Step 2 was omitted. Instead, the Step 3 reaction was primed using the crude Step 1 reaction mixture at a 1:50 dilution.

After completion, Step 3 PCR reactions were diluted 2.5-fold with water, supplemented with 1/10^th^ volume of 5 M Ammonium Acetate (pH5.0), and the resulting PCR products were precipitated by the addition of 2.5 volumes of absolute ethanol at room temperature, washed with 80% ethanol following centrifugation, and resuspended to a concentration of 0.1-0.2 µg/µl. The conversion of µg/µl to µM for DNA was performed using the equation: µM = (µg/µl) * (1620/PCR product length). Run-off transcription using T7 RNA polymerase was performed for 2.5 h at 35 °C in a buffer containing 40 mM Hepes-KOH (pH 7.9), 22 mM Mg(OAc)_2_, 2 mM spermidine, 40 mM DTT, 5 mM of each rNTP, 0.25-0.75 μM DNA template, 50 μg/mL T7 polymerase, and 25 μg/mL yeast inorganic pyrophosphatase. The transcripts were purified by ion-exchange chromatography on an ethanolamine–Sepharose matrix as described previously(*69*, *70*). Briefly, the transcription reactions were terminated by adding 0.5 volumes of a buffer comprising 1M NaOAc (pH 5.2) and 2.5 mM EDTA, 1 volume of the 20% matrix suspension, and 2.5 volumes of water. After a 15-min incubation with agitation at RT, the matrix was extensively washing with a buffer containing 120mM NaOAc (pH 5.2) and 0.25 mM EDTA. Subsequently, the transcripts were eluted using an elution buffer consisting of 2 M NaOAc (pH 5.2), 10 mM Mg(OAc)_2_, and 0.25 mM EDTA, then precipitated by adding 2.5 volumes of absolute ethanol. The RNA pellet was washed twice with 75% ethanol containing 2.5mM NaOAc (pH5.2), followed by a wash with 80% ethanol, and finally resuspended in water to a concentration of 100 µM. The conversion of µg/µl to µM for RNA was performed using the equation: µM = (µg/µl) * (2900/transcript length).

### Split-tRNA assembly

Typically, split-tRNAs were assembled by combining chemically synthesized 5’-fragments with either chemically synthesized or T7-transcribed 3’-fragments. The latter and the former fragment types were dissolved in 0.5 mM NaOAc (pH5.2) to concentrations of 100 µM and 250 µM, respectively. Annealing reactions were carried out in a 56 µl reaction volume containing the 3’- and 5’-fragments of split-tRNA at concentrations of 50 µM and 55 µM, respectively, with NaOAc (pH 5.2) adjusted to 5mM. The annealing mixtures were heated to 80°C for 1.5 min in a PCR machine. Upon completion, the PCR machine’s power was turned off, and 2 µl of 290 mM MgCl_2_ were immediately added to a final concentration of 10 mM, holding the samples in the PCR machine. The mixtures were immediately stirred by mechanically moving the pipet tips while holding the pipet’s plunger pressed. After mixing, the samples were allowed to gradually cool to RT. The RNA was precipitated by adding 2.5 volumes of an ethanol/Ammonium Acetate precipitation solution (ethanol/AmAc), consisting of 25 volumes of absolute ethanol and 1 volume of 5M Ammonium Acetate (pH 5.0). The RNA pellet was washed and dissolved as described for the final stage of purification of T7 transcription products.

### RNA fractionation in denaturing and native polyacrylamide gel

Denaturing 10% polyacrylamide gel (19:1, acrylamide:bis-acrylamide) containing 7 M urea in 1x TBE buffer was typically loaded with 0.5-1 pmol of RNA sample and run in 1xTBE at 180 V for 35 min. The gel thickness was 1 mm. RNA bands were visualized by staining with SYBR Green I (Thermo Fisher). The Cy3- or Cy5-labeled products were visualized by fluorescence scanning at 602 nm and 700 nm, respectively, using ChemDocTM Imaging system (BioRad). Under native conditions, RNA samples were loaded on 8% polyacrylamide gel (29:1, acrylamide:bis-acrylamide) prepared with the buffer containing 34 mM Tris, 66 mM Hepes resulting in pH7.5, 0.1 mM EDTA, 3 mM MgCl_2_, and run in the same buffer(*72*). Before loading, RNA samples were adjusted to 5µM with 2x native sample loading buffer containing 100 mM HepesKOH pH 7.5, 0.3 mM EDTA, 5mM MgCl_2_, and 20% glycerol. 1 µl, corresponding to 5 pmols (∼120 ng) of RNA sample, was loaded per slot.

### Split-tRNA Quality Control: evaluation of 5’- and 3’-end integrity and aminoacylation status

The translation reaction was prepared in a 17 µl volume containing 10 µM of split-tRNA. Following the reaction assembly, 5 µl aliquots, corresponding to 40 pmols of split-tRNA, were withdrawn either immediately (while kept on ice) or at time points 35 and 85 minutes during the translation time course. The aliquots were pipetted to a mixture comprising 50 µl of water-saturated phenol and 60 µl of 5 mM NaOAc, pH 5.3, vigorously shaken, and then frozen. The mixtures were subsequently processed in parallel; 40 µl of chloroform was added, followed by shaking at 1600 rpm for 10 minutes at room temperature (RT). The samples were then centrifuged at 12,000 × g for 15 minutes at 4°C. 55 µl of the upper water phase was collected and transferred into 145 µl of a pre-chilled precipitating solution (refer to “Split-tRNA assembly”). Subsequently, the samples were centrifuged at 17,000 × g for 15 minutes at 4°C, and the RNA pellets were directly resuspended by pipetting in 5.5 µl of water, followed by the addition of another 5.5 µl of 5mM NaOAc, pH 5.3.

Half of the sample (equivalent to 20-25pmols of split-tRNA) underwent oxidation with sodium periodate (NaIO4, SigmaAldrich) for 25 minutes at RT in the reaction, assembled wit 1 µl of 0.5M NaIO₄, 2 µl of 0.5M NaOAc, pH 5.0, 2 µl of water, and 5 µl of the RNA sample(*73*, *74*). The other half of the sample was incubated in parallel in the premix, where NaIO4 was replaced with 0.5M NaCl. Subsequently, both reactions were precipitated with cold 26 µl of ethanol/AmAc. The pellets were resuspended in 5 µl of water and supplemented with 4 µl of 0.25M glucose to deplete the residuals of NaIO₄. After a 15-minute incubation at RT, both mixtures were adjusted with 1 µl of 1M Tris-HCl, pH 9.0, and incubated for an additional 30 minutes at 37°C to deacylate tRNAs.

Subsequently, the mixtures were precipitated with ethanol/AmAc, the pellets were washed twice with 75% ethanol, and resuspended in 5 µl of water.

The aliquots of oxidized and non-oxidized samples were used in parallel for primer extension and ligation assays. Before the reverse transcription reaction, 4 pmols of split-tRNA (equivalent to 1 µl of each sample) were combined with 20 pmols of Cy5-labelled DNA-oligonucleotide in a volume of 4.5 µl. The mixture was incubated at 80°C for 7.5 minutes, followed by gradual cooling to room temperature (RT).

The reverse transcription reaction was performed in 10 µl of 1x First-strand buffer (Invitrogen) containing the annealing mixture, 0.5 mM dNTPs, 5 mM DTT, and 10 units/µl of SuperScript™ III RT. The reactions were incubated for 30 minutes at 55°C, followed by inactivation of reverse transcription at 70°C for 5 minutes. The products were precipitated with ethanol/AmAc. Pellets were resuspended in 2 µl of water and added with 8.5 µl of denaturing sample buffer (SB) containing 95% formamide, 25 mM EDTA, and Bromphenol Blue. The mixture was heated for 5 minutes at 95°C, and 2 µl was loaded onto a 10% polyacrylamide gel (19:1, acrylamide:bis-acrylamide) containing 7 M urea in 1xTBE buffer. The loaded volume corresponds to ∼0.8pmol of split-tRNA, equivalent to the maximal obtainable amount of extension product.

Ligations of 3’-oxidized or control RNA samples with the Cy3-labelled hairpin-adopting DNA oligonucleotide were performed in 5 µl of 1x T4 DNA ligase buffer (Thermo Fisher/Fermentas) containing 15% DMSO, 4 pmol of split-tRNA, 20 pmol of Cy3-labelled DNA hairpin, and 1 unit/µl of T4 DNA ligase, following the procedure inspired by the earlier publication(*75*). After 15-hour incubation at 16°C, the reaction mixtures were precipitated, and samples were reconstituted with SB as described in the section on the reverse transcription reaction, but heated for 5 minutes at 80°C.

### Preparation of S30 extract

E. coli S30 extract was prepared as described (*46*) with minor modifications. The overnight BL21-(DE23)-Gold culture was diluted 1:100 into 5 L of filter-sterilized TBGG media (tryptone 12 g/L, yeast extract 24 g/L, glycerol 8 mL/L, glucose 1 g/L, KH_2_PO_4_ 2.31 g/L, K_2_HPO_4_ 2.54 g/L), distributed over 6 baffled conical flasks (850 ml per flask), and cultivated at 37C, 110 rpm until the log phase, corresponding to an OD of 3.5. Cell batches were combined and pre-chilled with 5 x 200 mL of –80°C-frozen packs of LB broth, followed by centrifugation at 4,000 g for 15 min, and washing the pellet twice with 0.5 L of S30A buffer.

After the final wash, the cell pellet (20-30 g) was resuspended in 200% (v/w) S30B buffer, and the resulting cell slurry underwent fluidic disruption (Constant Systems, continuous flow mode) at 20 kpsi, at 4°C. The cell homogenate was repeatedly centrifuged at 30,000 g for 30 min, with 2/3 and 3/4 of the supernatant transferred following each centrifugation, respectively. The supernatant was adjusted with NaCl to a 0.4 M using 5 M NaCl stock and incubated for 45 min at 42°C in a water bath. Following incubation, the extract was subjected to dialysis using a 12-14 kDa cut-off dialysis membrane (SpectraPor®) for 2h against 5 L of S30C buffer, pre-chilled at 4°C, and containing 10 mM Tris-acetate (pH 8.2), 14 mM Mg(OAc)_2_, 0.6 mM KOAc, and 0.5 mM DTT, followed by overnight dialysis against fresh 5 L of the same buffer at 4°C. The dialyzed extract was centrifuged at 30,000 g for 30 min, and 3/4 of the supernatant was transferred, aliquoted, snap-frozen in liquid N2, and stored at -80°C.

### Preparation of S30 extract depleted of total tRNA fraction

The procedure was modified from the previously published method(*76*). Prior to ion-exchange chromatography, the S30 *E. coli* extract was rebuffered from the low potassium/high magnesium conditions of the S30C storage buffer into the high potassium/low magnesium conditions of buffer D containing 10 mM Hepes-KOH (pH7.5), 25 mM KCl, 10 mM NaCl, 0.1 mM EDTA, 1.1 mM Mg(OAc)_2_, and 100 mM KOAc. For this purpose, a PD-10 Superdex 25 column (GE Healthcare) equilibrated with buffer D was loaded with 2.5 ml of S30 extract, followed by idle loading of 0.5 ml of buffer D, and then eluted with an additional 2.5 ml of the same buffer. The amount of ethanolamine-Sepharose resin corresponding to 2 ml of 50% slurry (refer to the ‘Preparation of ethanolamine-Sepharose resin’) was drained of storage buffer by spinning in a filter-bottom tube at 1,000 g for 2 min, washed twice with buffer D, and drained again following another round of centrifugation. The semi-dried resin was mixed with the 2.5 ml of rebuffered lysate, followed by incubation of the resulting slurry in a 15 ml flask for 30 min at 4°C with rotation. After incubation, the slurry was spun down, transferred to a filter-bottom tube, and centrifuged at 1,000 g for 2 min at 4°C. While the flow-through was stored on ice, the semi-dried resin was transferred back to the 15 ml flask and mixed with 1 ml of buffer D containing 180 mM KOAc. Following a 15-minute incubation at 4°C with rotation, the following steps were repeated as described, and the respective flow-throughs were pooled to yield ∼3.5 ml of depleted lysate containing potassium and magnesium ions at total concentrations of 150 mM and 1.1 mM, respectively. The depleted lysate was aliquoted, snap-frozen in liquid N_2_, and stored at -80°C.

### Specific tRNA depletion from total tRNA mixture

Specific tRNA species were depleted from the purified total tRNA fraction by DNA-hybridization chromatography as described previously (*45*, *69*). The amount of total tRNA used for depletion was calculated to ensure a 20-fold molar excess of immobilized oligonucleotide over the target tRNA fraction in the total tRNA mixture. The tRNA concentration in a 20 µg/µl total tRNA fraction can be approximately calculated to be 720 µM using the equation in “Synthesis and purification of tRNAs and tRNA fragments” and assuming the average size of tRNA molecule to be 80 nt. The percentages of specific tRNA fractions in the total tRNA mixture of *E. coli*, ranging from 0.3% to 7.3%, can be found in Dong et al. (*47*). Therefore, if a hypothetical target tRNA constitutes 1% of the total tRNA fraction, 200 µl of 50% oligonucleotide-conjugated resin suspension, equivalent to 20 nmol of bait, can be used to deplete this target tRNA (1 nmol) from 130 µl of 20 µg/µl total tRNA. To assemble the hybridization reaction, a semi-dried oligonucleotide-conjugated resin corresponding to 200 µl of 50% suspension, drained of solution in a filter-bottom tube by centrifugation at 1,000 g for 1 minute, was mixed with 260 µl of 10 µg/µl total tRNA mixture and 260 µl of 2x hybridization buffer (*77*) containing 20 mM Bis-Tris HCl (pH6.5), 1.8 M tetramethylammonium chloride (TMA-Cl), and 0.2 mM EDTA. The hybridization was performed at 78°C for 15 min with constant agitation at 1200 rpm, followed by gradual cooling to RT at a rate of 1°C per minute, with continuous agitation. The hybridization mix was transferred to a filter-bottom tube, and the flow-through was collected following centrifugation at 1000 g for 1 min. Subsequently, the resin underwent two successive washes: first with one reaction volume of 10 mM Bis-Tris HCl (pH6.5), then with a buffer containing 10 mM Bis-Tris HCl (pH6.5), 10 mM MgCl_2_, and 100 mM NaCl. All three flow-throughs were combined and precipitated as described in “Split-tRNA assembly”. The pellet of total tRNA depleted of specific tRNA was resuspended in water to a concentration of 20 µg/µl.

### Assembly of cell-free translation with normal and tRNA-depleted lysates

The translation reactions in a normal (unmodified) lysate were assembled using 24% (v/v) of S30 extract, 12% (v/v) of 1x S30C buffer containing 10 mM Tris-acetate pH 8.2, 14 mM Mg(OAc)_2_, 0.6 mM KOAc, 0.5 mM DTT, 40% (v/v) of feeding solution (236 mM Hepes-KOH pH 7.4, 12.5 mM Mg(OAc)_2_, 375 mM KOAc, 5% PEG 8000, 5 mM DTT, 2.5x protease inhibitor (cOmplete™ EDTA-free, Roche), 0.25 mg/mL folinic acid, 2 mM of each rNTP with an extra 1 mM of ATP, 38 mM of acetyl phosphate, 68 mM of creatine phosphate, 1.25 mM of each amino acid with extra 2.5 mM for Arg, Cys, Trp, Asp, Met, Glu), 0.05 mg/mL T7 RNA polymerase, 45 U/mL creatine phosphokinase, and 20 nM of plasmid template.

Translation reactions with depleted lysate were assembled similarly to those with normal S30 extract, with the following modifications: 36% (v/v) of S30 extract depleted of total tRNA fraction was used in the reaction, the feeding solution contained 24 mM Mg(OAc)_2_ and 243 mM KOAc, and reactions were additionally supplemented with 1 µg/µl of the total tRNA fraction specifically depleted for the given endogenous tRNA (Fig. S1).

Free Mg^2+^ concentration in translation reactions was estimated to fall within 3 to 4 mM, taking into account that ∼4mM of the 10 mM total Mg^2+^ was complexed by rNTPs, and ∼2.8 mM of the remaining 6mM was complexed by creatine phosphate (K_d_ = 25 mM)(*78*).

### Preparation of Leishmania-based cell-free translation system

A Leishmania-based transcription-translation system (LTE) was prepared as described by Kovtun et al.(*48*). *Leishamania tarentolae* cells were pre-cultured in 500 ml of TBGG media (see “S30 extract preparation”) supplemented with 1 ml of 500x Hemin solution (0.25% of Hemin in 50% triethanolamine, 0.22 µm filter-sterilized) to OD ∼3-3.5. This pre-culture was then expanded at a 1:10 dilution to 5 L TBGG media with Hemin, distributed as 1 L per 5 L conical baffled flask. The cells were grown at 26.5 °C with agitation at 74 rpm for approximately 24 hours.

Cells were harvested at a density of 1.0-1.2 x 10^8^ cells/ml (corresponding to OD 3-3.5), pelleted at 2,800 g for 15 min with low deceleration, and resuspended to a concentration of 10^10^ cells/ml in a buffer containing 45 mM Hepes-KOH pH 7.6, 250 mM sucrose, 100 mM KOAc, and 3 mM Mg(OAc)_2_. Cells were then disrupted using a nitrogen cavitation vessel by equilibrating the cell suspension under 70 bar nitrogen pressure for 45 min at 4°C. The lysate was clarified by centrifugation at 10,000 g, followed by supernatant transfer and centrifugation at 30,000 g.

The top two-thirds of the final supernatant was collected and subjected to gel filtration on PD-10 Superdex 25 column (GE Healthcare) equilibrated with a buffer used for cell resuspension but lacking sucrose. 2.5 ml of cleared lysate was loaded onto the column, followed by idle loading of 0.25 ml of the same buffer, and then elution with 2.5 ml of the same buffer.

The 2.5 volumes of the buffer-exchanged lysate was then supplemented with 1 volume of 5x feeding solution containing 6 mM ATP, 0.68 mM GTP, 22.5 mM Mg(OAc)_2_, 1.25 mM spermidine, 10 mM DTT, 200 mM creatine phosphate, 100 mM Hepes-KOH pH 7.6, 5% (v/v) PEG 3000, 5.25x protease inhibitor cocktail (Complete™ EDTA-free, Roche), 0.68 mM of each amino acid, 2.5 mM rNTP mix (ATP, GTP, UTP and CTP), 0.05 mM anti-splice leader DNA oligonucleotide (αSL oligo, supplementary data 2), 0.5 mg/ml T7 RNA polymerase, 200 U/ml creatine phosphokinase. The supplemented lysate was snap-frozen and stored at -80°. Transcription-translation reactions were assembled by adjusting 7 μl of supplemented lysate to a final reaction volume of 10 μl with the plasmid template at a final concentration of 20-40 nM and chimeric (split-)tRNA at 5 or 10 µM.

### Translation in PURE

Translation reaction in the reconstituted system was performed using PURExpress In Vitro Protein Synthesis Kit (NEB, #E6800S) accordingly to the manufacturer’s protocol. PURE translation plasmid is used at 25nM final concentration PURE 6800 is used in 5 ul containing 2 ul of Sol A, 1.5ul of sol B, 0.05ul of RNAsin and 25nM of plasmid template.

### Reverse Transcription coupled quantitative PCR analysis of tRNA depletion

For the quantitative comparison of the individual tRNA fractions in the S30 lysate, the 2.5 ul of its dilutions, ranging from 0.004mg/ml to 0.4mg/ml of total material exhibiting absorption at 260 nm, were subjected to treatment with DNAse I. In addition to S30 lysate the 10 ul of DNAse I treatment reaction also contained 1 ul of DNAse I buffer (NEB #B0303) and 1 ul of DNAse I (2u/ul, #M0303). The reaction mixture was incubated at 37°C for 1h. Subsequently, 0.5 ul of 50mM EDTA was added to chelate Mg2+ ions, and DNAse I was inactivated by heating the reaction at 75°C for 10 min. Following DNAse I treatment, 2.5 ul of a 1:5 dilution of the reaction in water was mixed with 2.5 ul of a 1uM reverse specific oligonucleotide. The mixture was heated to 78°C for 8 min and then gradually cooled at a rate of 0.1°C per sec to RT.

The DNAse I treatment step was deemed unnecessary and omitted when assessing individual tRNA fractions in the total tRNA mixture. Instead, 2.5 μL of total tRNA dilutions, with concentrations ranging from 2.5 to 40 ng/μL, were directly used for the annealing reaction with the corresponding reverse oligonucleotide as described above.

The reverse transcription (RT) reaction was assembled in a 10 ul volume, comprising 4 ul of annealing reaction, 1 ul of 10mM dNTPs mixture, 0.2 ul of Avian Myeloblastosis Virus (AMV) Reverse Transcriptase, and 1 ul of its 10x Buffer (NEB, #M0277). A control reaction, lacking reverse transcriptase, was included to assess potential contamination with genomic DNA containing the amplification target. The reactions were incubated at 50°C for 45 min, followed by the inactivation of reverse transcriptase at 85°C for 5 min. For qPCR, from 0.5 to 2.5 µl of RT reaction was used. Quantitative PCR reactions were performed in the format of 384-well plate (PerkinElmer, #6007290), in 12.5 ul volume, containing 6.25 ul of SsoAdvanced Universal SYBR® Green Supermix (BioRad, #172-5270), 0.5 ul of each 10uM forward and reverse oligonucleotides and 0.025 μL of ROX dye (ThermoFisher). The amplification was performed using the CFX384 Touch Real-Time PCR Detection System (BioRad). Alternatively qPCR reactions were assembled in a similar way using Platinum™ SYBR™ Green qPCR SuperMix-UDG (Thermofisher). The standard cycling program, including preheating at 95°C for 3 min, followed by 40 cycles of denaturation at 95°C for 15 s and extension at 60°C for 30 s was employed. Subsequently, melting curve analysis was conducted over the temperature range between 65°C and 95°C using a default program of the respective Real-Time PCR system.

### RNA structures alignment

See Supplementary Table 2 for the detailed alignment pattern. Briefly, codon-anticodon stem loop complexes (C-ACSL) were superimposed via the conformationally restrained codon triplets. The A-helices were superimposed on the codon-anticodon complexes through the respective nucleotide patterns in the A-helices and anticodon stems. The pdb codes and sources of A-helices structures: 2mkn(*79*), 2az2(*80*), 2jxq(*81*), 6db9(*82*), 433D(*83*), 472d(*84*), 5e7k(*85*). The pdb codes and sources of C-ACSL structures: 6of6 (ala)(*86*), 4v6f (phe)(*87*), 2uu9 (val)(*88*).

### Linear Regression Workflow

A schematic overview and step-by-step description of the analysis workflow are provided in Fig. S13. Further implementation details are available in the README file of the GitHub repository (https://github.com/mureich81/tRNASelectionModel.git).

## Acknowledgments

The authors would like to acknowledge Dr. Patricia Walden and Micaela Fiorito for their excellent organizational assistance. The authors also thank Dr. Elena Eremeeva for her help with RNA fractionation, Roxane Mutschler for providing reverse transcriptase, and Dr. Zhong Guo for discussions on some aspects of enzyme kinetics.

## Author Contributions

Conceptualization: SM

Methodology: SM, ZC

Investigation: SM, YW

Visualization: SM

Funding acquisition: KA

Project administration: KA, SM

Supervision: SM, KA

Writing – original draft: SM, KA

Writing – review & editing: SM, KA

## Data and materials availability

The code used for analyzing trends in kinetic parameters of split-tRNA selection is available on GitHub at https://github.com/mureich81/tRNASelectionModel.git. For detailed information, refer to the README file in the GitHub repository or to the step-by-step description of the workflow in Fig. S13.

The structures of canonical and wobble-containing A-helices are available under the following PDB entries: 2mkn(*79*), 2az2(*80*), 2jxq(*81*), 6db9(*82*), 433D(*83*), 472d(*84*), 5e7k(*85*). The structures of codon-anticodon stem-loop complexes correspond to the following PDB entries: 6of6 (ala)(*86*), 4v6f (phe)(*87*), 2uu9 (val)(*88*).

