## Supplementary notes, figures and tables for "Codon decoding by split-tRNA: revisiting the tRNA selection mechanism"

### Supplementary Text:

#### 1. Obtaining relative $\boldsymbol{k_{cat}/K_m}$ for GTP hydrolysis using fluorescence rescue assay

tRNA selection occurs in two stages: initial selection and proofreading, separated by an irreversible GTP hydrolysis step.

Since decoding site closure occurs on a timescale of tens of milliseconds (*17*, *18*), which implies a high activation barrier between the open and closed states, and the cognate ternary complex is stabilized within the closed state by a factor of 40–50 (*50*, *89*), the decoding site is likely to remain closed around the canonical codon–anticodon minihelix during the accommodation step. Accordingly, the rejection rate for cognate substrates is expected to be low. Therefore, we assumed comparable probabilities of productive selection during proofreading for different substrates, regardless of their anticodon arm context, as long as cognate codon–anticodon pairing is maintained.

The tRNA selection rate can be defined as the product of the GTP hydrolysis rate and the probability of successful passage through the proofreading step. Given the above assumption, this probability can be neglected, as it remains similarly high for cognate substrates and will be cancelled out. Thus, the selection rate can be approximated by the GTP hydrolysis rate and expressed as the product of the apparent rate constant ($\frac{k_{cat}}{K_m}$), the concentration of the respective ternary complex (TC), and the concentration of the elongation complex (EC):

$$Rate = \left(\frac{k_{cat}}{K_m}\right) \cdot [TC] \cdot [EC] \qquad \text{Eq. S1}$$

The rate of fluorescent eGFP accumulation is determined by the efficiency with which the chimeric alanyl-tRNA ($tRNA^{Ala}$), carrying a grafted arginine anticodon and incorporating alanine, competes with wild-type arginyl-tRNA ($tRNA^{Ala}$) at a key arginine codon, which replaces the wild-type alanine codon at position 226 of mutant eGFP (Fig. 2).
Under non-competitive reaction conditions, where only the fluoresent product is formed, the maximal fluorecence accumulation velocity ($V_{max}^{non-comp.}$) can be expressed as the rate of selection of chimeric $tRNA^{Ala}GCG$ or $tRNA^{Ala}CCU$, incorporating alanine at the arginine codon ($CGC\ or\ AGG$) at position 226 ($Rate^{ala}$), adjusted by a scaling factor ($m$). This factor accounts for additional kinetic steps during protein synthesis, folding, and chromophore maturation, the conversion of protein yield to fluorescence yield, and also includes the invariant concentration of elongation complexes ($[EC]$). Thus from Eq. S1 we obtain Eq. S2:

$$V_{max}^{non-comp.} = m \cdot \left(\frac{k_{cat}}{K_M}\right)^{ala} \cdot [TC^{ala}] \qquad \text{Eq. S2}$$

Under competitive reaction conditions, the maximal fluorescence accumulation velocity ($V_{max}^{comp.}$) is given by the product of $V_{max}^{non-comp.}$ and the probability ($p$) of successful selection of $tRNA^{Ala}GCG$ on $CGC$ codon over the competing $tRNA^{Arg}(\frac{I}{G})CG$, as selection of the later results in a non-fluorescent product (Eq. S3):

$$V_{max}^{comp.} = m \cdot \left(\frac{k_{cat}}{K_M}\right)^{ala} \cdot [TC^{ala}] \cdot p \qquad \text{Eq. S3}$$

Therefore, from Eqs. S2 and S3, $p$ is given by the ratio of the maximal fluorescence accumulation velocities under competitive and non-competitive conditions:

$$p = \frac{V_{max}^{comp.}}{V_{max}^{non-comp.}} \qquad \text{Eq. S4}$$

On the other hand, $p$ can also be expressed as the ratio of the selection rate of chimeric $tRNA^{Ala}GCG$ to the sum of this selection rate and that of the competing $tRNA^{Arg}(\frac{I}{G})CG$. Therefore, from Eq. S1 cancelling out $[EC]$, we obtain Eq. S5:

$$p = \frac{Rate^{ala}}{Rate^{ala} + Rate^{arg}} = \frac{\left(\frac{k_{cat}}{K_M}\right)^{ala} \cdot [TC^{ala}]}{\left(\frac{k_{cat}}{K_M}\right)^{ala} \cdot [TC^{ala}] + \left(\frac{k_{cat}}{K_M}\right)^{arg} \cdot [TC^{arg}]} \qquad \text{Eq. S5}$$

From Eqs. S4 and S5 we obtain Eq. 2 of the main text connecting the apparen rate constants of tRNA selection and :

$$\frac{V_{max}^{comp.}}{V_{max}^{non-comp.}} = \frac{\left(\frac{k_{cat}}{K_M}\right)^{ala} \cdot [TC^{ala}]}{\left(\frac{k_{cat}}{K_M}\right)^{ala} \cdot [TC^{ala}] + \left(\frac{k_{cat}}{K_M}\right)^{arg} \cdot [TC^{arg}]} \qquad \text{Eq. 2}$$

Substituting the estimated concentrations of both ternary complexes into Eq. 2, grouping the rate constants for chimeric $tRNA^{Ala}$, and dividing both sides by the rate constant for $tRNA^{Arg}$ results in the Eq. S6. This equation represents the apparent second-order rate constant for $tRNA^{Ala}$ selection normalized by the respective rate constant for the selection of the competing $tRNA^{Arg}$ :

$$\left(\frac{k_{cat}}{K_M}\right)^{ala} / \left(\frac{k_{cat}}{K_M}\right)^{arg} = \frac{[tRNA^{Arg}]}{[tRNA^{Ala}]} \cdot \frac{V_{max}^{comp.}/V_{max}^{non-comp.}}{1 - V_{max}^{comp.}/V_{max}^{non-comp.}} \qquad \text{Eq. S6}$$

In the equation, $[tRNA^{Ala}]$ and $[tRNA^{Arg}]$ represent the concentrations of the respective aminoacylated fractions in the translation reaction mixture.

### 2. Exploring intrinsic and extrinsic factors that may influence codon reading by G37-containing split-tRNA variants

The lack of correlation between the estimated stacking strength and decoding efficiency observed for certain G/C-rich N37N38 split-tRNA variants may be attributable to intrinsic and/or extrinsic factors. Initially, we aimed to rule out extrinsic factors, such as the accelerated degradation of the anticodon dangling ends or differences in aminoacylation levels occurring in an N37N38-dependent manner. A primer extension assay on RNA isolated from *in vitro* translation reactions at various time points indicated no truncation of the anticodon termini over the 85-minute reaction period (Fig. S8). In parallel, we measured the aminoacylated fraction of the split-tRNAs at different time points during translation reaction. These revealed that by the log phase of fluorescence accumulation, most of split-tRNA variants were aminoacylated to around 40%, with the G37C38, C37G38, and G37G38 variants being aminoacylated to 25-30% (Fig. S8). While partial dimerization via protruding 34-38 strands may potentially explain the variations in the first two variants, the last variant showed no dimerization (Fig. S10). We concluded that the concentrations of aminoacylated split-tRNA fractions, ranging from 2.5 to 4 µM, were unlikely to be rate-limiting factors responsible for the observed discrepancies.

In addition to thermodynamic effects inherent to specific N37N38 combinations, intrinsic factors contributing to the observed discrepancies may include the propensity of G/C-rich protruding strands to form intermolecular dimers or intramolecular quadruplet-like motifs, which may hinder aminoacylation or the accommodation of the anticodon terminus within the decoding site, alongside the previously reported N37N38-dependent coordination of $Mg^{2+}$ ions by the codon-anticodon complex (see below).

To evaluate the formation of intermolecular structures we analysed split-tRNAs using native gel electrophoresis. These experiments revealed different degrees of split-tRNA dimerization, particularly notable in split-tRNAs carrying C37A38, C37G38, and G37C38 flanks (Fig. S10). The modifications including iso- or 8-aza-7-deaza-deoxyguanosine at position 37, which interfere with canonical Watson-Crick or Hoogsteen interactions, suppressed the dimerization of the G*37C38 split-tRNA variants (Fig. S10). We concluded that inter- or intramolecular secondary structure is unlikely to be the primary cause of the observed variations in decoding efficiency.

The involvement of $Mg^{2+}$ ions in the codon-dependent stage of Aa-tRNA selection remains a subject of ongoing research. Several studies have investigated the relationship between the equilibrium binding constant for peptidyl-tRNA interaction with the A-site and $Mg^{2+}$ concentration, concluding that the identity of N37 influences the number of $Mg^{2+}$ ions involved in this interaction (*51*, *90*). Another study also found that $Mg^{2+}$ influences the rates of both codon-independent binding and codon recognition steps of tRNA selection (*8*). In contrast, Johansson *et al.* observed an inverse linear relationship between translational accuracy and $Mg^{2+}$ concentration. This led to the conclusion that $Mg^{2+}$ primarily affects the initial codon-independent step of ternary complex binding by slowing its dissociation (*5*, *91*).

Given the structural differences between native and split-tRNA, we conjectured that they should respond differently to changes in $Mg^{2+}$ concentration should it be an integral part of codon recognition. We did not observe significant differences in the impact of increased $Mg^{2+}$ concentration on the relative decoding efficiencies of split-tRNAs regardless of their N37N38 identity (Fig. S11).

3. Structural analysis of wobble interaction within N37N38 segment of split-tRNA

Superimposition of the available canonical and 5'U:G3'-tandem-containing RNA helices onto the codon-anticodon stem loop structure demonstrated no discernible bias in relative alignment among them (Fig. S12). The deviations in geometry are likely to be influenced more by the choice of anchor nucleobases used in the alignment rather than by the impact of the wobble configuration. The G from the wobble base pair, coinciding with the N37, displays either greater or lesser overlap with the underlying N1:N36 base pair, depending on the assembly (Fig. S12, C to E). The absence of a clear trend among the three assemblies could be attributed to the nested positioning of the wobble pairs in the available templates used for alignment, which may not accurately represent the behaviour of the terminally located wobble pairs due to intrahelical constraints. Previous studies have indeed highlighted the differences in thermodynamic stability and conformational flexibility between terminal and nested wobble base pairs (*92*, *93*).

4. Evaluation of dominant tRNA selection models in light of current findings

Aa-tRNA selection is a complex process in which a supramolecular catalytic machinery discriminates among diverse multisubunit substrates at multiple selection stages. While extensive studies have provided insights into individual aspects of this mechanism, a fully integrated framework that reconciles all observations remains elusive (*5*, *16*).

The selection process begins with the initial codon-independent association of the ternary complex (T3) with the ribosome (R), resulting in complex C0 (step 1). This is followed by the codon-recognition step (step 2), resulting in complex C1, where the codon-anticodon interaction occurs. Step 3 involves a conformational change in which the decoding site locks around the first and second positions of the codon–anticodon minihelix, provided they conform to Watson–Crick geometry. This crucial step, which is also referred to as GTPase activation, facilitates the remote assembly of the transition state for GTP hydrolysis, which occurs at the final step (step 4).

$$R + T_3 \underset{k_{-1}}{\overset{k_1}{\rightleftarrows}} C_0 \underset{k_{-2}}{\overset{k_2}{\rightleftarrows}} C_1 \underset{k_{-3}}{\overset{k_3}{\rightleftarrows}} C_2 \overset{k_4}{\rightarrow} GDP + P_i \qquad \text{Scheme 1}$$

Below, we provide a detailed characterization of the molecular events at each stage, along with their interpretation through the lens of a substrate–enzyme analogy:

**Step 1**: The ternary complex (T3) is initially tethered to the ribosome via the L10-$(uL12)_{2\text{-}3}$ stalk, establishing two contacts with the ribosomal subunits: one with the 30S subunit shoulder via EF-Tu, and another between the tRNA elbow and the uL11 stalk of the 50S subunit.

**Step 2**: Codon sampling by the anticodon loop of the tRNA results in codon–anticodon pairing. Upon successful recognition, the tRNA adopts a kinked conformation accompanied by partial unwinding of the anticodon stem and adjacent core domains (Supplementary Text 6). In the enzyme–substrate analogy, this corresponds to the substrate (T3) engaging with the enzyme

(R) by presenting its anticodon to the codon of the mRNA, which serves as a polymeric cofactor (Fig. 4A). From a thermodynamic perspective, the affinity between substrate and enzyme is determined by both the entropic gain from water expulsion from the interacting surfaces and the enthalpic contribution from inter-polymeric hydrogen bonding between codon-anticodon triplets, as well as stacking interactions within the codon-anticodon complex. The difference in thermodynamic stability between cognate and near-cognate codon-anticodon complexes can serve as an initial source of selectivity. Due to the similarity in forward rates inherent to RNA-RNA interactions (*94*), the maximal discrimination achievable at this step is determined by the ratio of the dissociation rate constants, $\frac{k_{-2}^{nc}}{k_{-2}^{c}}$, for near-cognate (nc) and cognate (c) substrates.

**Step 3**: The transition from $C_1$ to $C_2$ is marked by closure of the small ribosomal subunit around the codon-anticodon minihelix. This conformational change involves a slight inward rotation of the 30S head and shoulder towards the intersubunit space, bringing them closer to the body and platform domains (*4*, *21*, *50*). Concurrently, the monitoring bases A1493 and A1492 from helix 44 of the 'body' domain, and G530 from helix 18 of the 'shoulder' domain relocate to engage the codon-anticodon duplex. These bases ensure that the first two codon-anticodon base pairs adhere to A-helical geometry, irrespective of sequence identity.

In the enzyme-substrate analogy, this stage marks a shift in the roles within the enzyme-substrate-cofactor triad: whereas in the preceeding step the anticodon served as the substrate's binding site and the codon triplet as a cofactor, in the context of the closed decoding site, the codon-anticodon minihelix assumes the role of an allosteric effector. This effector modulates the final assembly of the transition state for GTP hydrolysis by EF-Tu at a remote site (Fig. 4A). The minihelix presents the backbone of its minor groove to the monitoring bases, enabling formation of a uniform network of polar contacts at the first and second codon positions. The third position is not subjected to the same level of structural scrutiny. In near-cognate codon-anticodon complexes, mismatches at the first or second positions can impair solvation of exposed hydrogen-bond donors or acceptors, or result in steric clashes between juxtaposed purines (*21*). Additionally, purine–purine mismatches at the third codon position can impair the A-form helical geometry of the entire codon–anticodon duplex (*11*, *43*).

Ultimately, stable closure of the 30S subunit facilitates formation of the transition state at the distal GAC site for GTP hydrolysis.

**Step 4**: Upon successful assembly of the transition state, GTP hydrolysis by EF-Tu proceeds rapidly at the remote catalytic center of the large subunit. In contrast, impaired closure of the decoding site around near-cognate codon-anticodon duplex destabilizes the transition state and suppresses hydrolysis.

To rapidly discriminate between cognate and near-cognate substrates, translation system must accommodate several major constraints on the tRNA selection process:

**1.** Speed-accuracy trade-off: The tRNA selection must operate under strict temporal constraints, requiring the system to maintain the highest possible accuracy without compromising translation speed.

**2.** Genetic code degeneracy, where the number of amino acids exceeds the number of codon families.

**3.** Broad variation in codon-anticodon duplex stabilities.

Attempts to reconcile these constraints give rise to inherent contradictions in current interpretations of the selection process (*5*, *16*):

- Geometry-based evaluation of the codon–anticodon minihelix at step 3 can streamline selection by placing initial selection under full kinetic control ($k_3 \gg k_{-2}$), irrespective of codon–anticodon complex stability. However, strict monitoring of geometry at the third codon position is not feasible, as it would lead to delays associated with idle rejection of isoaccepting tRNAs. The existing strategy of monitoring the first two positons is compatible with "2-out-of-3" decoding in amino acids encoded by unsplit codon families.
- For tRNAs decoding split codon families, rejecting the third-position mismatch becomes critical. This requires either evaluation of codon–anticodon stability during initial selection to discriminate correct pairings from mismatches, or rejection of mismatches during the proofreading stage. The latter is unlikely, as the associated energetic cost would exceed the average hydrolysis of ~1.2 GTP molecules per peptide bond (*61*).
- The possibility of stability-based selection implies that codon-anticodon recognition occurs under near-equilibrium conditions. However, given the broad variation in codon–anticodon stabilities, allowing the system to approach equilibrium at this step would introduce delays incompatible with efficient translation.
- For effective stability-based selection, the stability of the least stable cognate codon-anticodon complex must exceed that of the most stable mismatch by a sufficient margin.

A viable hypothesis must either assume selective involvement of a thermodynamic component in controlling initial selection through the dissociation rate, or invoke an alternative mechanism in which discrimination operates via modulation of the forward rate.
Put simply, the question lies in identifying the primary driver: whether it is the cognate codon–anticodon interaction that promotes progression toward GTP hydrolysis, or the near-cognate interaction that acts by hindering it.

Conceptually, this distinction reflects the difference between induced fit and conformational selection models of substrate recognition. In the induced fit model, allosteric effector or substrate binding lowers the activation barrier for the conformational change leading to assembly of catalytically competent state, thereby accelerating the forward rate. In contrast the conformational selection model implies that the enzyme undergoes spontaneous conformational fluctuations, and the thermodynamic stability of the effector-enzyme complex determines the activation barrier for the transition state of catalytic step as well as for effector dissociation.

**Induced fit model**:

The classical definition of induced fit implies that substrate binding induces a conformational rearrangement of the enzyme, with part of the binding energy driving the structural change (*23*, *95*). In the context of tRNA selection, the energy derived from codon–anticodon interaction is proposed to contribute to rearrangement of the decoding site, thereby facilitating GTPase activation (*7*, *23*).

Below we examine theoretical constraints, prior experimental observations, and results from current semi-empirical modelling that challenge the applicability of the classical induced fit model to tRNA selection:

a) **Theoretical constraints**: Within the framework of induced fit model, a flexible (or "floppy") substrate is expected to reduce the activation barrier for active site assembly less efficiently due to a higher entropic cost involved. In the context of tRNA selection, the stability of the codon-anticodon complex would influence the forward rate of decoding site closure ($k_3$). The same stability also inversely affects the dissociation rate ($k_{-2}$).

It is entirely plausible to envision a cognate codon–anticodon complex whose stability is reduced to the point that $k_3$ becomes comparable to, or even lower than, $k_{-2}$. In such a case, the same stability difference between two substrates—both assuming canonical Watson–Crick geometry at codon positions 1 and 2—would contribute to selectivity via both $k_{-2}$ and $k_3$, effectively amplifying discrimination. However, reusing the same thermodynamic difference to enhance selectivity at two stages prior to the irreversible step violates basic thermodynamic principles.

b) **Previous observations supporting spontaneous open-closed fluctuations of the decoding site**:

- The ribosome can hydrolyze GTP in the EF-Tu–GTP binary complex even in the absence of tRNA (*65*).
- Modifying atomic groups of the monitoring bases that interact with the codon–anticodon minihelix has only a moderate impact on selection efficiency (*67*).
- Despite forming one of the least stable codon–anticodon complexes, $tRNA^{Lys}UUU$ shows elevated background miscoding frequencies (*11, 13*), likely because uridine can form sterically neutral mismatches with any codon base, permitting decoding site closure.
- Conformational dynamics studies show that the initial barrier associated with decoding site rearrangement is traversed on a similar timescale for cognate and near-cognate tRNAs occuping the A-site (*17, 18*). This is consistent with structure-informed MD simulations revealing similar activation barriers for the GTPase activation step (step 3) for both substrate types (*6*).
- The decoding site was found to sample the open conformation even after initial locking around the cognate codon-anticodon complex, highlighting the dynamic nature of the system (*64*).

c) Implication of the **current study:** Using semi-empirical modeling, we estimated discard parameters for a hypothetical split-tRNA lacking stacking interactions within the N36-N39 segment, which is expected to yield highly unstable codon-anticodon complex. The ~100-fold increase in the discard parameter $d_2$ (= $k_{-2}/k_3$) indicates that such unstable complexes can equilibrate during the codon recognition step. In contrast, the near-zero value of $d_3$ (= $k_{-3}/k_4$) suggests that reduced codon–anticodon stability does not influence the rate of GTP hydrolysis (step 3), provided that the canonical geometry of the codon-anticodon minihelix is preserved.

**Conformational selection model**:

This model proposes that the energy barrier for small subunit closure remains constant and is crossed when local thermally driven fluctuations within the macromolecular ribonucleoprotein complex of the 30S subunit reach a resonance condition. According to this framework, opening and closing of the decoding site occurs spontaneously at a constant average frequency. Discrimination between ternary complexes decoding split-codon families and near-cognate complexes with sterically neutral $3^{rd}$-position mismatches can occur either during the codon recognition step via $k_{-2}$, provided that it exceeds the rate of conformational step ($k_3$), or through more rapid reversal of the locked decoding site conformation via $k_{-3}$. As long as the mismatch at the $3^{rd}$ codon position does not affect the canonical geometry of the proximal part of codon-anticodon minihelix, $k_{-3}$ remains much smaller than $k_4$.

Based on this reasoning, we hypothesize that discrimination between cognate split-codon readers and their near-cognate counterparts occurs via lower and higher probabilities of their dissociation, respectively, during the codon recognition step. Accordingly, the activation barrier for decoding site closure must be positioned near the dissociation energy threshold of cognate substrates to enable discrimination of 3$^{rd}$-position mismatches under partial thermodynamic control.

Kinetic dissection of smFRET-resolved conformational dynamics during decoding revealed that, for tRNA(Phe), both dissociation from the cognate codon prior to 30S subunit closure (step 2) and the rate-limiting step of conformational transition preceding GTP hydrolysis (step 3) occur at comparable rates of 10-20 $s^{-1}$ (*17*, *89*). This rate range corresponds to an activation barrier of approximately 16 kcal/mol at 25°C, consistent with estimates from structure-informed, MD-based free energy simulations (*6*).

Given that codon–anticodon complexes formed by split and unsplit codon family readers segregate into lower- and higher-stability classes, respectively, an intermediate activation barrier enables preferential selection of the more stable complexes under kinetic control without compromising the trade-off between decoding speed and accuracy.

Below, we examine conflicting observations and limitations in the available data that preclude an unambiguous validation of the conformational selection model.

Comparative data on intrinsic dissociation rates of cognate versus near-cognate tRNAs remain scarce, as the widely reported ~1000-fold difference likely reflects, in part, selective stabilization of cognate tRNA within the closed decoding site (*2*, *4*, *8*, *17*, *19–22*).

The 40-50-fold excess stabilization granted by the decoding site can be inferred from smFRET-based conformational dynamics studies (*89*) and cryo-EM quantification of structural states (*50*).

Modelling of codon-anticodon energetics in constrained environment suggests 2.1 kcal/mol loss for G-U third-position wobble compared to G-C, projecting to 30-fold difference in dissociation rates (*96*). However, available data from conformational dynamics studies suggest only a modest, several-fold difference in intrinsic dissociation rates of tRNA(Phe) from cognate (UUU) versus near-cognate (CUU) codons (*17*, *18*). Yet, the authors do not exclude the possibility that near-cognate dissociation rates are underestimated due to missed short-lived events and preferential detection of long-lived outliers. Therefore, it remains possible that under physiological $Mg^{2+}$ concentrations, the actual difference in dissociation rates between near-cognate and cognate tRNAs is greater than observed.

The apparent dissociation rate constants for tRNA(Phe) from cognate and near-cognate codons at physiological Mg2+ concentrations, as reported in classical kinetic studies, vary in a codon-dependent manner within the ranges 0.12-0.23 and 100-240 $s^{-1}$, respectively (*8*).

After correcting for the contribution of 40-50-fold auxiliary stabilization from the decoding site, the intrinsic dissociation rates for cognate tRNA(Phe) fall within the range of 5–10 $s^{-1}$, consistent with values extracted from FRET trajectories (*17*, *18*). From this, the intrinsic difference in dissociation rate constants between cognate and near-cognate tRNAs is likely to exceed one order of magnitude, yet remaining modest for being the sole source of initial discrimination. Therefore, an additional contribution from the structural effect of a non-canonical base pair at the third codon position on minihelix geometry cannot be excluded. For instance, the elegant work by Zhang and colleagues demonstrated that the third-position misreading frequency increases with codon–anticodon stability, yet nuanced by anticodon composition (*14*).

### 5. Transforming Eq. 3 to relate decoding efficiency to the free energy of the N36-N39 segment

*Background*

The free energy of the N36-N39 segment is defined as the difference between its free and accommodated codon-bound states or, equivalently, as its contribution to codon-anticodon complex stability, alongside the contribution of the codon-anticodon minihelix itself.

Eq. 3 describes Scheme 1 (Supplementary text 4) for the initial stage of tRNA selection, and relates the apparent second-order rate constant ($k_{cat}/K_m$) for the GTPase reaction to the kinetic parameters $d_1, d_2,$ and $d_3$ that characterize different selection stages, where:

$$\frac{k_{cat}}{K_m} = \frac{k_1}{1 + d_1(1 + d_2(1 + d_3))} \qquad \text{Eq. 3}$$

and

$$d_1 = \frac{k_{-1}}{k_2}, d_2 = \frac{k_{-2}}{k_3}, d_3 = \frac{k_{-3}}{k_4}$$

We used the experimentally determined decoding efficiencies for each N37N38 split-tRNA variant, along with the estimated concentrations of competing ternary complexes, to express the $k_{cat}/K_m$-related term as a dependent variable.
The objective is to incorporate the free energies of the corresponding N37-N39 segments into Eq. 3 as independent variables. These energies, together with the invariant contribution from the three base pairs and two stacking interfaces of the codon-anticodon minihelix, are inherently embedded in the activation barriers of the forward and reverse rate constants that define the discard parameters for each N37N38 split-tRNA variant.
To incorporate these energies as independent variables in Eq. 3, we introduced universal surrogate discard parameters corresponding to a hypothetical split-tRNA lacking stacking interactions within the N36-N39 segment. Below, we outline how the discard parameters for each split-tRNA variant can be expressed as functions of the universal surrogate discard parameters scaled and the stability of the corresponding N36-N39 segment.

*Quantitative relationship between $d_1$ and N36-N39 stability term*

$$d_1 = \frac{k_{-1}}{k_2}$$

The initial binding of the ternary complex to the ribosome is independent of codon-anticodon interaction. In line with the general properties of RNA-RNA interactions, the subsequent codon-anticodon recognition proceeds with similar forward rates ($k_2$) for both cognate and near-cognate pairs (*94*). Therefore, given the similarity in both $k_{-1}$ and $k_2$ rate constants between cognate and near-cognate ternary complexes, the parameter $d_1$ remains unaffected by the strength of the codon-anticodon interaction.

*Quantitative relationship between $d_2$ and N36-N39 stability term*

$$d_2 = \frac{k_{-2}}{k_3}$$

The reverse rate, $k_{-2}$, is influenced by both the activation free energy required for accommodation of the anticodon terminus in the decoding site and the standard free energy change associated with codon–anticodon complex formation. The latter can be estimated by summing the stabilities of adjacent stacking units, using the nearest-neighbour model of nucleic acid stability. Accordingly, we assumed that $k_{-2}$ can be expressed as described in Eq. S7:

$$k_{-2} = X \cdot e^{\frac{-(dG^{\#}+(-dG_{C:AC})+(-dG_{N36-N39}))}{RT}} \quad \text{Eq. S7}$$

In this equation, X is a frequency factor (*95*); $dG^{\#}$ denotes the activation free energy required for accommodation of the anticodon arm in the decoding site; $dG_{C:AC}$ represents the stability gain associated with formation of the codon:anticodon minihelix, comprising three base pairs and two stacking interfaces; and $dG_{N36-N39}$ corresponds to the free energy difference between the accommodated and unaccommodated states of the N36-N39 segment. The latter reflects additional stabilization of the codon–anticodon complex due to its integration into a continuous stacking column that includes both the minihelix and the core helix formed by the anticodon arm and the D-stem.

$dG_{C:AC}$ and $dG_{N36-N39}$ enter the reverse rate equation with negative signs, reflecting the reversed order of free energy subtraction for bound versus unbound states. Substitution of Eq. S7 into the expression for $d_2$ yields Eq. S8:

$$d_2 = \frac{1}{k_3} \cdot X \cdot e^{\frac{-(dG^{\#}+(-dG_{C:AC})+(-dG_{N36-N39}))}{RT}} \quad \text{Eq. S8}$$

As mentioned above, to express the $d_2$ parameter for each N37N38 split-tRNA variant in terms of the stability of its N36-N39 segment, we introduced a surrogate discard parameter, $d_{2*}$, corresponding to a hypothetical split-tRNA$^{\Delta37\Delta38}$ variant lacking stacking interactions within the N36-N39 region. In this construct, $d_2^*$ is defined solely by the activation free energy and the stability of the codon-anticodon minihelix:

$$d_2^* = \frac{1}{k_3} \cdot X \cdot e^{\frac{-(dG^{\#}+(-dG_{C:AC}))}{RT}} \quad \text{Eq. S9}$$

Since RNA:RNA interactions are primarily governed by the dissociation rate, regardless of the length and stability of the base-paired segment (*94*), we assume that the activation free energy associated with the transition of the anticodon arm from the free to the codon-bound state is similar between the hypothetical split-tRNA$^{\Delta37\Delta38}$ and any of the sixteen N37N38 split-tRNA variants. Accordingly, from Eqs. S8 and S9, we derive Eq. S10, which expresses $d_2^{n37n38}$ for each N37N38 split-tRNA variant as the product of the surrogate discard parameter $d_2^*$ and the corresponding stability term $e^{\frac{dG_{N36-N39}}{RT}}$:

$$d_2^{n37n38} = d_2^* \cdot e^{\frac{-dG_{N36-N39}}{RT}} \quad \text{Eq. S10}$$

*Quantitative relationship between $d_3$ and N36-N39 stability term*

$$d_3 = \frac{k_{-3}}{k_4}$$

Similar to $d_2$, the stability of the codon-anticodon complex may also influence the discard parameter $d_3$ through either of its components (Scheme 1), resulting in an algebraically equivalent outcome for the parameter's value. The reverse rate constant $k_{-3}$ is determined by the activation free energy required for decoding site closure, increased by the stabilization gained within the closed complex. This stabilization can be conservatively estimated as the sum of two contributions: (i) the free energy difference between the closed-state interaction of the monitoring bases with the minor groove and their baseline stabilization in the stacked conformation of the open complex, and (ii) a possible additional contribution from reduced thermal fluctuations upon integration into a contiguous stacking framework involving the 36–39 segment.

Accordingly, we assume that, similar to $d_2$, the $d_3$ parameter for any given N37N38 split-tRNA variant can be derived from the surrogate parameter $d_3^*$ by scaling the reverse rate constant for decoding site closure with the stability of the corresponding N36–N39 segment:

$$d_3^{n37n38} = d_3^* \cdot e^{\frac{-dG_{N36-N39}}{RT}} \qquad \text{Eq. S11}$$

*Development of a regression model for discard parameter prediction*

$d_2^{n37n38}$ and $d_3^{n37n38}$ expressed in Eqs. S10 and S11 via the respective surrogate parameters scaled with the stability term $e^{\frac{dG_{N36-N39}}{RT}}$, denoted as $x$, can be then included into Eq. 3 to yield Eq. 4 correlating second apparent rate constant for the initial tRNA selection and the stability within N36-N39:

$$\frac{k_{cat}}{K_m} = \frac{k_1}{1 + d1\left(1 + d_2^* \cdot x\left(1 + d_3^* \cdot x\right)\right)} \qquad \text{Eq. 4}$$

Explicitly including Eq. 4 into Eq. 2 yields Eq. S12:

$$\frac{V_{max}^{comp.}}{V_{max}^{non-comp.}} = \frac{k_1 \cdot [c_S] \cdot x^2}{\left(k_1 \cdot [c_S] + N \cdot [c_{wt}] \cdot (1 + d_2^*)\right) \cdot x^2 + N \cdot [c_{wt}] \cdot d_2^* d_3^* \cdot x + N \cdot [c_{wt}] \cdot d_1 d_2^* d_3^*} \qquad \text{Eq. S12}$$

In this equation $N$ represents the apparent second order rate constant $\left(\frac{k_{cat}}{K_M}\right)^{arg}$ for competing wild-type intact $tRNA^{Arg}$, $[c_S]$ and $[c_{wt}]$ denote the concentrations of the aminoacylated N37N38 split-tRNA variant and competing synthetic $tRNA^{Arg}$GCG, respectively, which are utilized as proxies for the concentrations of the respective ternary complexes.

Defining $\frac{V_{max}^{comp.}}{V_{max}^{non-comp.}}$ as $f$, Eq. S12 can be transformed into a linear polynomial form, expressed as Eq. S13:

$$\frac{(1-f) \cdot k_1 \cdot [c_S]}{f \cdot N \cdot [c_{wt}]} - 1 = d_1 + d_1 d_2^* \cdot x + d_1 d_2^* d_3^* \cdot x^2 \qquad \text{Eq. S13}$$

Assuming $k_1$ and the second apparent rate constant $N$ for wild-type intact $tRNA^{Arg}$ (*8*) are of similar magnitudes, these terms cancel out, yielding the final Eq. 5:

$$\frac{[c_S] \cdot (1-f)}{[c_{wt}] \cdot f} - 1 = d_1 + d_1 d_2^* \cdot x + d_1 d_2^* d_3^* \cdot x^2 \qquad \text{Eq. 5}$$

When applied to each N37N38 split-tRNA variant, this results in an overdetermined system of 16 equations suitable for regression analysis.

6. tRNA core and misreading: Why is Trp-tRNA unique?

Since codon-anticodon recognition and GTP hydrolysis occur 75Å apart, it is attractive to assign the role of transducer to the tRNA itself owing to its structural integrity and natural ability to connect the decoding site and the GTPase center. Indeed, structural studies showed obligatory distortion of tRNA core upon codon recognition in the context of the ternary complex (*24, 97, 98*). Accordingly, several hypothesis such as 'Alignment/Misalignmnet' (*99*), 'Molecular Spring' (*100*) and 'tRNA Waggle' (*101*) have been proposed to explain a possible discrimination mechanism. These hypothesis implicate the role of 'strain' within the distorted tRNA as a critical factor influencing the alignment and/or stability of the near-cognate ternary complex for productive GTP hydrolysis. However, cryo-EM-informed MDS, confirmed by single molecule spectroscopy, suggested spontaneous sampling of distorted conformations by tRNA at low energy barrier (*17, 63, 64*). These same distortions were confirmed for both cognate and near-cognate tRNAs (*62*), suggesting that the bent and unbent tRNA states are similar in stability and separated by a low energy barrier on the order of *k*T.

Extensive experiments on mutagenesis within the elbow of $tRNA^{Trp}$ yielded various miscoding-prone variants (*102–105*). This raises the question of whether the observed effect indicates the core's involvement in signalling for GTPase activation in response to codon recognition, or if it merely reflects the impact of distal part of the stacking column on codon-anticodon complex stability.

Early work by Hirsh and colleagues revealed that mutation in the core of tRNA, distant from the anticodon loop, can result in miscoding. Specifically, they showed that $tRNA^{Trp}$CCA harboring a G24A replacement in the D-stem gains ability to read-through the UGA stop codon via A(3)-C(36) wobble at the 3rd codon position. Yarus and colleagues identified alternative mutations at positions 9 or 27-43, which also resulted in a miscoding-prone phenotype (*102–106*). Following a global selection of miscoding variants from a randomized pool of $tRNA^{Trp}$, mutations at position 59 were shown to result in miscoding (*107*).

Schultz and Yarus analysed the effect of all N27:N43 combinations on the ability of $tRNA^{Trp}$CUG to miscode "amber" UAG codon via G(36)-U(1) first position wobbling (*104*). They concluded that the presence or absence of a canonical N27-N43 base-pair correlates only weakly with miscoding properties, with most of the effect being independently contributed by individual nucleotides at positions N27 and N43. For example, G27-C43, which represents a mere reversal of the wild-type base pair, also yielded a miscoding-prone variant. They also observed that mismatched nucleotides at positions N27-N43 did not result in a temperature-sensitive phenotypes, indicating no structural defects or stability losses.

These observations are unexpected, given that N27-N43 is the most proximal pair of the anticodon stem mediating its coaxial stacking with the D-arm. Predicting the effect of various alterations within the core is complicated due to the amalgamation of multiple factors. On one hand, mismatches within the vertical stack would be expected to exert a destabilizing effect. On the other hand, the mutated region is far from the anticodon end of the anticodon stem/D-stem stack to strongly impact codon-anticodon interaction. Importantly, mutant sites fall within the conformationally dynamic region that undergoes unwinding when tRNA adopts an A/T state upon codon-anticodon recognition in the context of the ternary complex. This region includes two hinges which span the segment from bp 30:40 to bp 26:45 and the adjacent core-segment assembled of the G10-C25-G45 / U11-G24 / U12-A23-A09 / C13-G22-7MeG46 / A14-s4U8-A21 / A15-U48 / h2U16-G59, respectively.

The phenomenon of coding adjustment through the alterations within these segments is unique to $tRNA^{Trp}$ and has been well characterized biochemically (*102–105*, *107*), structurally (*106*) and kinetically (*26*, *107*). The intermediate position of this hinge between the decoding site and the GTPase centre led to three alternative explanations for how its mutations lead to miscoding. These explanations involve modulation of interaction kinetics with the decoding site(*103*), impact on the rate of GTPase activity (*26*), or influence on the subsequent accommodation rate (*107*). Schultz and Yarus proposed a hypothesis explaining why point mutations at diverse sites of $tRNA^{Trp}$ lead to miscoding-prone variants rather than to the expected increase in codon reading stringency(*105*). As mentioned earlier, evolutionary pressure drives biological systems to balance efficiency and accuracy, with the trade-off generally biased towards efficiency. The promiscuous recognition of synonymous codons is a natural mechanism for enhancing translational efficiency. Given that $tRNA^{Trp}$ decodes a single codon, UGG, it may have been finely tuned by evolutionary pressure to ensure hyperaccuracy, preventing any wobble recognition that could risk stop codon read-through or the incorporation of the bulky Trp in place of cysteine. Additionally, $tRNA^{Trp}$ encodes the least frequent amino acid, so the compromise between accuracy and efficiency may, in the case of $tRNA^{Trp}$, favour accuracy, as it would not significantly reduce protein elongation speed.

At the same time, according to our assessment, $tRNA^{Trp}$CCA forms a codon-anticodon complex of exceptionally high stability due to both, consecutive G-C pairs in the 2nd and 3rd codon positions and the optimal context in the anticodon loop. This stability stands apart from the typical range of stabilities observed in tRNAs decoding split codon families (table S1). It appears that the increase in decoding stringency, in the case of $tRNA^{Trp}$, is not attained by manipulating the anticodon loop context, but rather through saturating adjustments within the core, potentially resulting in altered overall plasticity.

Hence, it is highly probable that most modifications introduced to the core would revert the constraints in $tRNA^{Trp}$, thereby alleviating the decoding restrictions. Essentially, the acquisition of a miscoding phenotype by $tRNA^{Trp}$ can be perceived as a reversion to the standard level of tRNA promiscuity. For instance, the U11-G24 wobble base pair appears to specifically occur in *E. coli* $tRNA^{Trp}$, whereas in other tRNAs, this position is occupied by standard Watson-Crick base pair (*37*).

**Fig. S1**

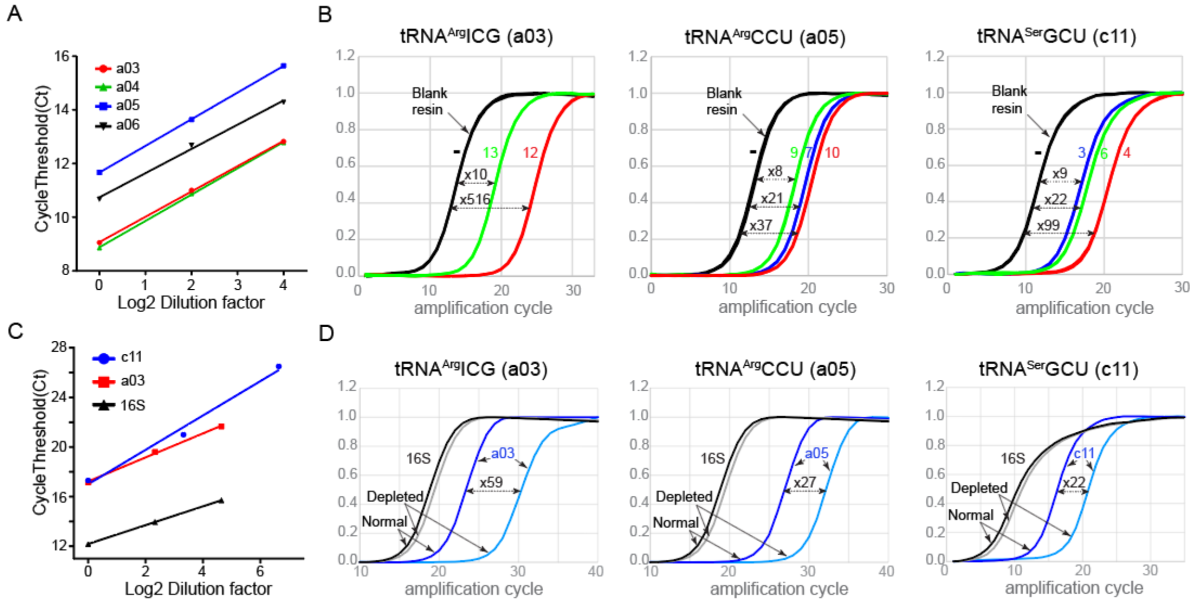


E

| Depleted tRNA: | tRNA$^{Arg}$ICG (a03) | | | tRNA$^{Arg}$CCU (a05) | | | | tRNA$^{Ser}$GCU (c11) | | | |
|---|---|---|---|---|---|---|---|---|---|---|---|
| RT-qPCR oligos: | H03/D05 | | | H05/D09 | | | | G04/F11 | | | |
| Pulldown oligo N: | - | **12** | **13** | - | **7** | **9** | **10** | - | **3** | **4** | **6** |
| Flowthrough (FT)* | 10.6 | 20.2 | 14.7 | 14.8 | 20.1 | 18.6 | 20.8 | 8.1 | 12.6 | 16.1 | 13.6 |
| Int. standard (FT) | 8.4 | 9.0 | 9.2 | 8.4 | 9.3 | 9.3 | 9.2 | 8.4 | 9.7 | 9.8 | 9.3 |
| Fold depletion (FT): | NA | **515.9** | **9.7** | NA | **20.6** | **7.7** | **36.5** | NA | **8.5** | **98.5** | **22.2** |
| Washing (W)* | NA | 12.5 | 11.9 | NA | 12.1 | 12.2 | 13.5 | NA | 13.0 | 16.3 | 14.1 |
| Int. standard (W) | NA | 10.4 | 8.4 | NA | 10.2 | 9.7 | 9.2 | NA | 9.7 | 9.4 | 10.1 |
| Fold enrichment (W): | NA | **82.5** | **12.7** | NA | **139** | **65.3** | **163** | NA | **0.8** | **1.2** | **0.4** |

F

| **RTqPCR oligos:** | **H03/D05** | | **H05/D09** | | **G04/F11** | |
|---|---|---|---|---|---|---|
| **NC** | **aq** | **Lt** | **aq** | **Lt** | **aq** | **Lt** |
| | 28.3 | 27.0 | 27.2 | 27.6 | 26.3 | 25.4 |

G

| Target RNA: | a03[1] | a05[1] | 16S[1] | c11[2] | 16S[2] |
|---|---|---|---|---|---|
| RT-qPCR oligos: | H03 F D05 R | H05 F D09 R | 16S_F 16S_R | G04 F F11 R | 16S_F 16S_R |
| Ct depleted* | 26.9 | 29.4 | 16.2 | 17.3 | 6.6 |
| Ct normal | 20.1 | 23.8 | 15.3 | 12.2 | 5.9 |
| Ct normal (-RT) | 28.1 | 30.9 | 29.6 | 30.0 | 27.9 |
| Fold depletion: | **58.7** | **27.4** | NA | **21.6** | NA |

**Assessment tRNA$^{Arg}$ICG, tRNA$^{Arg}$CCU, and tRNA$^{Ser}$GCU depletion from purified total tRNA or crude S30 lysate by RT-qPCR.** Specific depletion from the total tRNA fraction was achieved by affinity pulldown with immobilized complementary DNA oligonucleotides, while nonspecific depletion from the S30 lysate was accomplished via ion-exchange chromatography on ethanolamine-sepharose (see Methods). (**A**) Standard curve from RT-qPCR analysis of three dilutions (40ng/µl, 10ng/µl and 2.5ng/µl) of the total E. coli tRNA fraction using oligonucleotides specific for tRNA$^{Arg}$ICG (a03), tRNA$^{Arg}$CCG (a04), tRNA$^{Arg}$CCU (a05), and tRNA$^{Arg}$UCU (see panel E). To maintain consistency, the total tRNA concentration in the 10ng/µl and 2.5ng/µl dilutions was adjusted to 40 ng/µl by mixing E. coli tRNA with 3 or 15 volumes of inert total yeast tRNA (both at 40 ng/µl). The x-axis represents the log2-transformed dilution factors, while the y-axis shows the corresponding cycle threshold (Ct). (**B**) Normalized RT-qPCR amplification signals for total tRNA batches specifically depleted of the indicated tRNAs. Numbers next to each amplification curve, color-coded accordingly, represent the oligonucleotide variants complementary to degenerate regions of each tRNA. Depletion folds were calculated as the exponential of the difference between Ct values from the depleted tRNA batch and the 'mock-depleted' batch (processed on blank resin). These fold values, shown above the respective double-headed arrow, were normalized by the exponential of the difference between Ct values for tRNA$^{Val}$GAC, which served as an internal standard to control for variations in total tRNA inputs (see panel E). (**C**) As in (A), but RT-qPCR performed on S30 lysate dilutions, with 16S ribosomal RNA (16S) as the internal standard. (**D**) As in (B), but assessing the depletion extent of target tRNA in S30 lysate depleted of all tRNAs (marked as depleted) by comparing amplification signals from the depleted lysate to those from undepleted (marked as normal) lysate. Depletion folds, shown above the double-headed arrow, were normalized using relative 16S rRNA content to account for differences between the depleted and reference lysates (see panel G). (**E**) The table summarizing the numerical data for specific depletion of $tRNA^{Arg}$ICG, $tRNA^{Arg}$CCU, and $tRNA^{Ser}$GCU from the total tRNA fraction. “Pulldown oligo” indicates the immobilized complementary oligonucleotide number, while a dash indicates a blank resin sample. “Flowthrough” (FT) corresponds to Ct values from the initially depleted fraction collected after incubation with the affinity resin. “Washing” (W) corresponds to Ct values from concentrated fractions pooled after the washing steps (see Methods). An asterisk denotes the mean Ct value from two parallel pulldowns. The internal standard (‘Int. standard’) corresponds to Ct values for flowthrough (FT) or washing (W) fractions using oligonucleotides H10 and H11 specific for $tRNA^{Val}$GAC as described in (B). "Fold depletion" and “Fold enrichment” represent the reduction in target tRNA concentration in the initial flowthrough (FT) and the enrichment of target tRNA in the washing flowthrough, respectively, both normalized by the internal standard. “Fold enrichment” represents an inverse indication of complex stability between the pulled tRNA and the complementary immobilized DNA oligonucleotide. Yellow-highlighted columns represent pulldown fractions used in subsequent experiments. (**F**) Negative control Ct values obtained using either water (aq) or total tRNA from *Leishmania tarentolae* (Lt) to prime RT-qPCR reactions. (**G**) Similar to (E), but assessing the depletion of target tRNAs from depleted lysate compared to normal lysate. Superscripts 1 and 2 correspond to data from two independent experiments. ‘-RT’ indicates control reactions without reverse transcriptase, used to monitor contamination with genomic DNA containing the amplification target.

**Fig. S2**

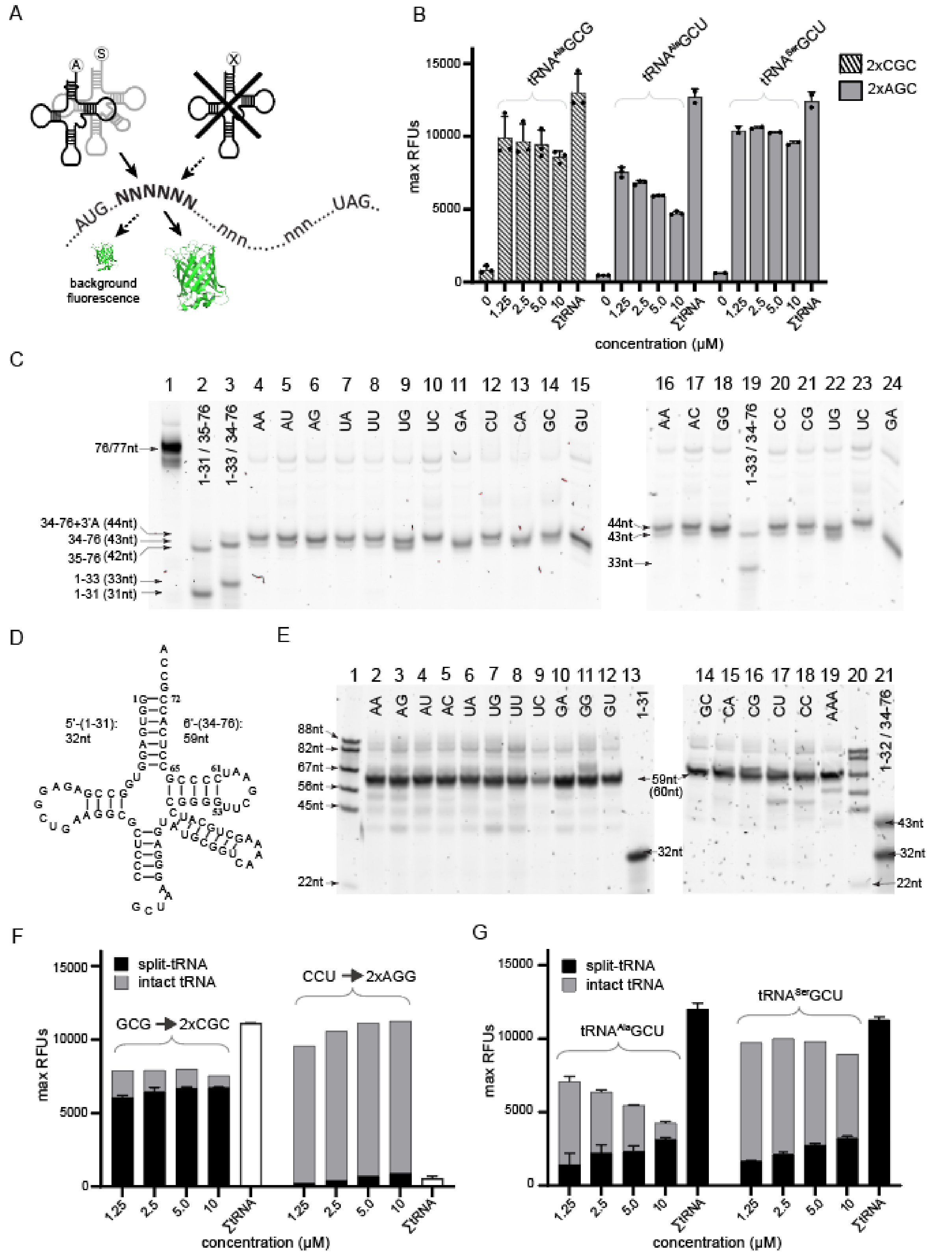


**Evaluation of split-tRNA functionality by a two-codon complementation assay.** The assay was conducted in a noncompetitive translation environment using total tRNA mixtures depleted of tRNA$^{Arg}$ICG, tRNA$^{Arg}$CCU and tRNA$^{Ser}$GCU to evaluate decoding efficiency on consecutive CGC, AGG, and AGC codons, respectively (2xCGC, 2xAGG, 2xAGC in supplementary data 2). (**A**) Schematic representation of a synthetic codon-biased ORF featuring two consecutive codons of interest ('NNN NNN') inserted upstream of the Lys-3 codon in the wild-type GFP coding sequence. The tandem codons make decoding highly sensitive to the concentration of the respective suppressor. Codons denoted as ‘nnn’ belong to a different codon family than ‘NNN’, are decoded by alternative isoaccepting tRNAs that do not recognize ‘NNN’, and encode the native amino acid positions within the ORF. The ‘cloverleaf’ structures on the left represent engineered alanine-accepting tRNAs (A) or wild-type $tRNA^{Ser}$GCU (S), where $tRNA^{Ala}$ is grafted with GCG, CCU, or GCU anticodons to create the respective chimeric variants. On the right, the crossed-out tRNA represents native ‘NNN’ codon suppressors depleted from the lysate using affinity chromatography via pull-down on DNA oligonucleotide-conjugated resin (see Methods). The dashed arrow indicates trace residual tRNA responsible for background fluorescence. (**B**) Concentration-dependent performance comparison of intact synthetic tRNAs, based on engineered alanine tRNA with GCG or GCU anticodons, and wild-type $tRNA^{Ser}$GCU for decoding codon-biased templates containing consecutive CGC and AGC codons (see plasmids 6281 and 6227 in supplementary data 2). The Y-axis shows maximal RFU values, while the X-axis represents concentrations of the corresponding synthetic suppressor tRNAs. Reactions with zero synthetic suppressor tRNAs indicate background translation supported by residual endogenous $tRNA^{Arg}$ICG or $tRNA^{Ser}$GCU remaining in the depleted lysate or total tRNA mixture after pull-down. ‘ΣtRNA’ denotes reactions supplemented with total *E.coli* tRNA at 1 µg/µl, reconstituting the standard *in vitro* translation environment. (**C**) Denaturing 10% urea-PAGE (19:1) loaded with size markers, including intact $tRNA^{Ala}$CCU in lane 1 and split-tRNAs assembled from synthetic 1-31/35-76 and 1-33/34-76 fragments in lanes 2, 3, and 19. The corresponding *in vitro* transcribed 34-76 fragments for all N37N38 combinations were loaded to lanes 4-18 and lanes 20-24. The size difference observed between the synthetic 34-76 fragment and the major bands of most *in vitro* transcribed fragments is due to the nontemplated addition of a 3’-end adenyl moiety by T7 polymerase. This adenyl moiety is efficiently trimmed by endonucleases present in the S30 extract, which normally participate in tRNA maturation following excision from the precursor molecule (Fig. S8). (**D**) ‘Cloverleaf’ representation of the split-tRNA version of wild-type $tRNA^{Ser}$GCU. Its recognition by aminoacyl-tRNA synthetase relies primarily on tertiary structure rather than sequence specificity and does not depend on the anticodon loop motif (*108*). (**E**) Similar to (C), *in vitro* transcribed 3’-(34-76)-fragments derived from wild-type $tRNA^{Ser}$GCU (lanes 2-12 and 14-19) were resolved on a 10% urea-PAGE gel. FAM-labelled DNA fragments served as size standards in lanes 1 and 20. Commercial synthetic 1-31 5’-fragment corresponding to $tRNA^{Ser}$GCU and split-$tRNA^{Ala}$GCG assembled of 1-32/34-76 fragments were used as size markers in lanes 13 and 21, respectively. (**F**) Analysis of translation competence of split-tRNAs with GCG or CCU anticodons 3’-flanked by A37A38, compared to intact chimeric $tRNA^{Ala}$ with the respective anticodons, using a “two-codon” complementation assay. Split-tRNAs were assembled as shown in Fig. 1B. Total tRNA was used at 1 µg/µl. (**G**) Similar to (F), but evaluating split-tRNAs with GCU anticodon 3’-flanked with A37A38, either based on $tRNA^{Ala}$ or derived from wild-type $tRNA^{Ser}$GCU (as shown in (D)). Translation reactions were depleted of endogenous $tRNA^{Ser}$GCU (Fig. S1) and programmed with a template containing consecutive AGC codons (supplementary data 2).

**Fig. S3**

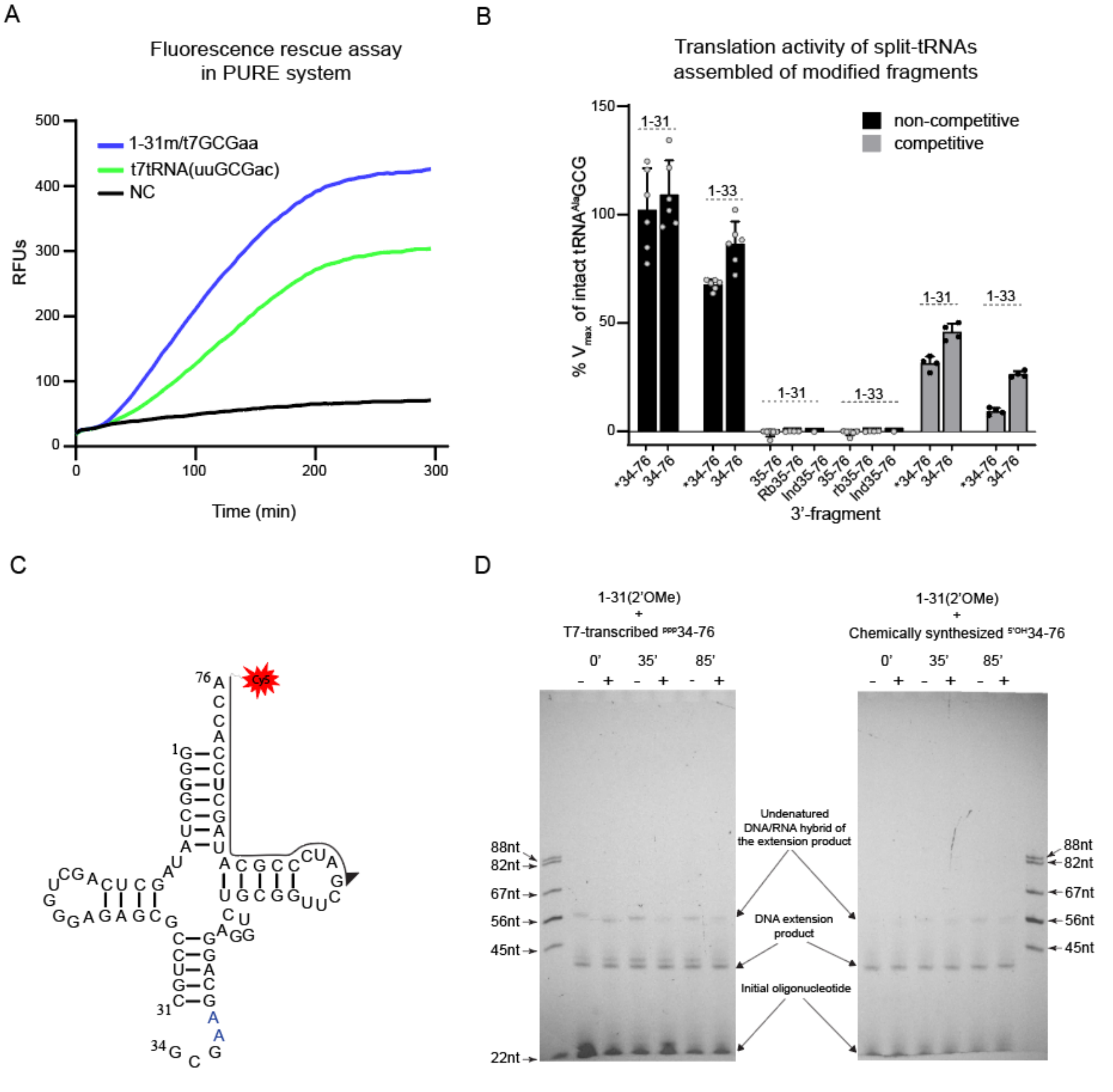


**Analysis of split-tRNA architecture preservation during *in vitro* translation.** (**A**) Fluorescent product accumulation in *in vitro* translation reactions using the PURE system, primed with mRNA encoding the A226R(CGC) GFP sensor, and supplemented with either split-tRNA$^{Ala}$GCG (blue line) or intact chimeric tRNA$^{Ala}$GCG (green line) at 10 µM and 3.5 µM, respectively. 'm' denotes 2'OMe at the 5'-(1-31)-fragment (F5' in supplementary data 2). The negative control reaction, lacking an exogenous CGC-codon reader, is represented by the black line.

(**B**) Relative decoding efficiency of split-tRNAs assembled from modified 5'-fragments (shown above each bar group) and 3'-fragments (X-axis). The 5'-fragments span positions 1-31 or 1-33, resulting in either no extension of the native anticodon stem or its extension by two base pairs (U32U33/A37A38) (F5 and F6 in supplementary data 2) (Fig. 1C). Both fragments feature a 3'-C3 Spacer (-O(CH2)$_3$OH) at C31 or U33. The 3'-(34-76)-fragments carry GCG anticodon with A37A38 3'-flanking sequence, with an asterisk indicating variants containing a 5'-C3 Spacer at G34. Fragments labelled as 35-76 lack G34 to test CGC codon recognition based on the 'two-out-of-three' decoding principle. 'Rb' and 'Ind' denote an abasic site filled with a ribose harboring a 3'-hydroxyl group or a 5-nitroindole residue replacing G34, respectively (F7-F10, F12 in supplementary data 2). The abasic modification accounts for potential stacking interaction between O4' of the N34 ribose and the pyrimidine ring of C1054, while the 5-nitroindole, lacking base-specific hydrogen bonding, tests whether 34/35 stacking is sufficient for productive decoding. The Y-axis represents the percentage of maximal fluorescence accumulation velocity of each split-tRNA relative to intact tRNA$^{Ala}$GCG. Experiments were performed under competitive conditions with S30 *E.coli* extract (grey bars) or noncompetitive conditions using extracts depleted of endogenous tRNA$^{Arg}$ICG (back bars). Error bars represent the standard deviation across independent replicates (dots on bar charts). (**C**) Schematic sequence representation of chimeric split-tRNA$^{Ala}$GCG annealed with a Cy5-labelled DNA primer for reverse transcription. (**D**) Fluorescence scans of 10% urea-PAGE gels showing primer extension products from total RNA extracted at indicated time points during *in vitro* translation. Reactions were supplemented with split-tRNA$^{Ala}$GCG (as shown in (C)), assembled from the same 1-31(2'OMe) 5'-fragment and either T7-transcribed (left gel) or synthetic 34-76 3'-fragmnets (right gel). The '+' and '-' signs indicate sample fractions subjected to or not subjected to periodide oxidation (for details, see Fig. S7 and Fig. S8 and their legends).

**Fig. S4**

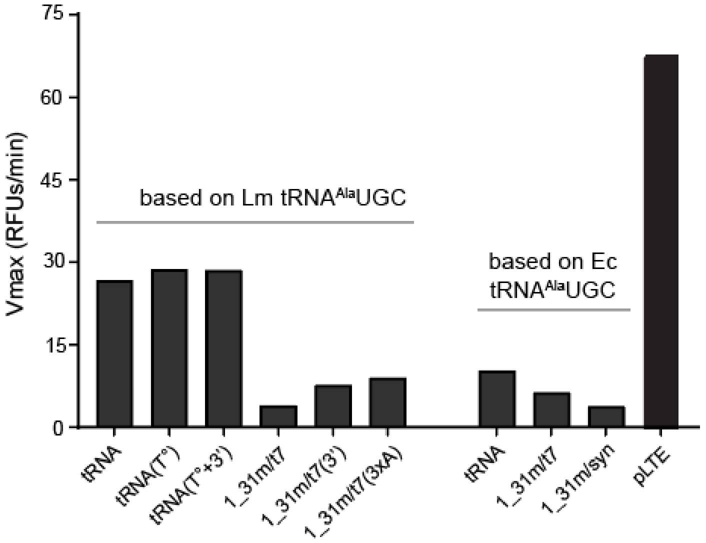


**Analysis of split-tRNA's ability to support translation in a eukaryotic *in vitro* translation system based on *Leishmania tarentolae* extract (LTE)** (*48, 49*) **using fluorescence rescue assay.** The plot shows the maximal fluorescence accumulation velocity ($V_{max}$) for reactions supplemented with either intact or split-tRNA$^{Ala}$GCG constructed from tRNA$^{Ala}$UGC derived either form *L. major* (Lm) or E. coli (Ec), as indicated above the bars. Split-tRNAs were assembled from synthetic 2'OMe-modified 5'-fragments ('1_31m') and either T7-transcribed ('t7') or synthetic ('syn') 3'-fragments (34_76). 'T°' indicates tRNA samples refolded after T7-transcription. " 3' " indicates DNA templates with two terminal 2'OMe-modified nucleotide residues (mGmU-5') at the 5' end (supplementary data 2) to prevent T7 polymerase from adding extra non-templated bases during transcription (*109*). This modification was included to address the potential lack of 3'-proofreading activity in LTE, a mechanism responsible for efficient 3' terminus proofreading in the *E.coli* cell-free system. '3xA' indicates an extra adenine insertion whithin the 'N37N38' flanking sequence. 'pLTE' denotes a positive control reaction programmed with a pLTE plasmid encoding wild-type GFP (supplementary data 2).

**Fig. S5**

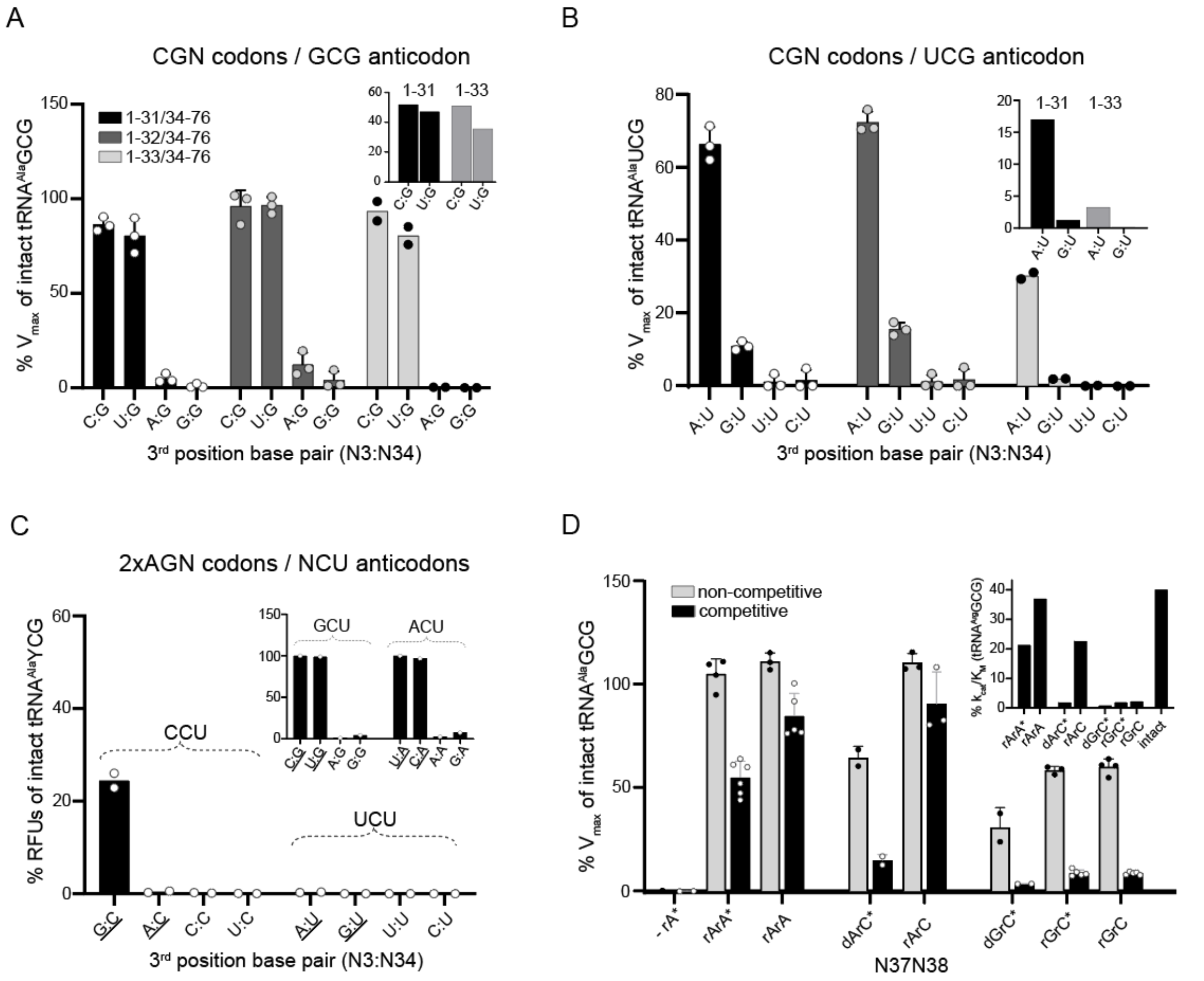


**Effect of 3$^{rd}$ position mismatch and N37 modification on decoding efficiency by split-tRNA compared to intact tRNA.** Translation reactions were conducted with an additional 5 mM $Mg^{2+}$ to enhance decoding efficiency unless specified otherwise. Decoding efficacy of CGN or AGN codons was assessed using either a fluorescence rescue assay or a two-codon complementation assay (Fig. S2). (**A**) Relative decoding efficiencies of CGN codons assessed using three homologous split-tRNAs, each featuring an invariant T7-transcribed 3'-fragment (positions 34-76) carrying a GCG anticodon and A37A38 flanking bases, combined with synthetic 5'-fragments spanning positions 1-31, 1-32, or 1-33 (F5, F16, F6 in supplementary data 2), resulting in no stem extension, a single base pair extension (U32-A38), or a two base pair extension (U32U33/A37A38), respectively. The Y-axis shows the percentage of maximal fluorescence accumulation velocity ($V_{max}$) for split-tRNAs relative to intact tRNA$^{Ala}$ bearing cuGCGaa anticodon loop (F7 in supplementary data 2), measured under non-competitive conditions using fractionated lysate supplemented with a tRNA complement lacking either tRNA$^{Arg}$ICG (for CGY and CGA codons) or tRNA$^{Arg}$CCG (for CGG codon) (Fig. S1). The inset shows CGC and CGU decoding efficiencies under competitive conditions. CGR codons exhibited negligible decoding efficiency and are excluded from the plot. (**B**) Similar to (A), but assessing the decoding efficincy of split-tRNA$^{Ala}$UCG on CGN codons relative to intact tRNA$^{Ala}$UCG. (**C**) Evaluation of split-tRNA$^{Ala}$ variants with CCU or UCU anticodons for decoding AGN codons using a two-codon complementation assay (cognate pairings underlined), performed under non-competitive conditions with fractionated lysate supplemented with a tRNA complement lacking tRNA$^{Arg}$CCG, tRNA$^{Arg}$CCU, tRNA$^{Arg}$UCU, or tRNASerGCU for decoding of AGR or AGY codons, respectively (Fig. S1). The inset shows relative decoding efficiencies of intact chimeric tRNA$^{Ala}$ with GCU or ACU anticodons on AGN codons. Notably, 4–7 % decoding of near-cognate codons via nonstandard 3'rd position G:G and A:G pairing was observed. (**D**) Bar chart showing the percentage of $V_{max}$ for various N37N38 split-tRNA variants relative to intact chimeric tRNA$^{Ala}$GCG under non-competitive and competitive translation conditions. Reactions were conducted without additional $Mg^{2+}$, with split-tRNAs supplemented at 10µM. Split-tRNA variants were assembled using the same 1-31 5'-fragment with a 2'OMe modification at the 3' end, combined with synthetic (marked by asterisk) or T7-transcribed 3'-fragments. '-' denotes an abasic site at position 37; 'd' and 'r' indicate deoxy- or ribonucleotides, respectively. Bars represent the average of two, four or six independent measurements, with error bars showing standard deviations. The inset shows percentages of $k_{cat}/K_m$ for each N37N38 split-tRNA variants relative to the $k_{cat}/K_m$ of the intact tRNA$^{Arg}$GCG competitor. 'intact' refers to the chimeric tRNA$^{Ala}$GCG with an intact anticodon loop.

**Fig. S6**

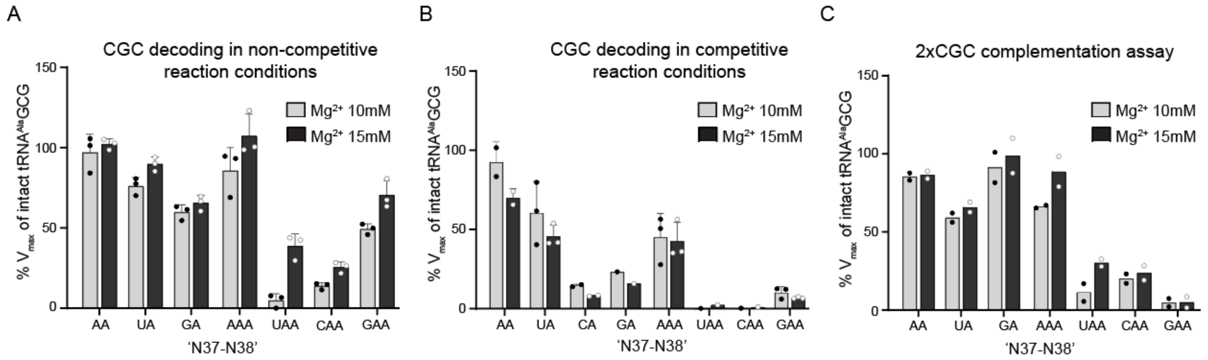


**Decoding of single and consecutive CGC codons by split-tRNAs with an extra nucleotide within the 'N37N38' anticodon flank.** (**A**) Comparison of relative decoding efficiencies between conventional N37A38 variants and those with an extra A. Reactions were conducted in a noncompetitive environment with either no additional $Mg^{2+}$ or 5 mM $Mg^{2+}$. (**B**) As in (A), but reactions conducted in a competitive translation environment. (**C**) As in (A), but using a template with two consecutive CGC codons.

**Fig. S7**

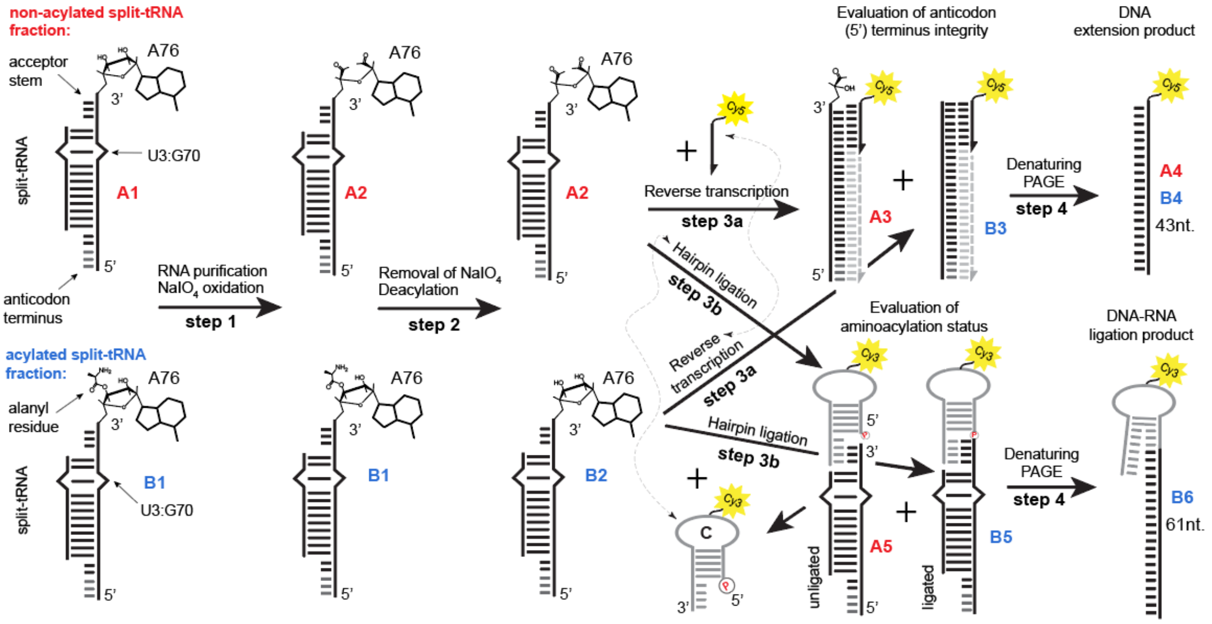


**Schematic overview of split-tRNA 5'-end integrity and aminoacylation status analysis during translation.** Split-tRNA is depicted as a stylized representation with the core shown as a double-stranded segment, featuring a U3:G70 wobble base pair - a critical determinant for alanylation, depicted as a bulge. The 3’-terminal adenosine (A76) is represented as a chemical structure. 'A' and 'B' denote the nonacylated and acylated split-tRNA fractions co-purified from the translation reaction. The acylated fraction (B1) is resistant to oxidation in step 1, while the free 2’- and 3’-hydroxyls of the A76 ribose in nonacylated split-tRNA (A1) are oxidized by $NaIO_4$, converting the ribose to a dialdehyde (A2). After deacylation in an alkaline environment (step 2), A2 and B2 are analyzed for 5'-end integrity (step 3a) and aminoacylation status (step 3b). In step 3a, RNA is annealed with a Cy5-labeled oligonucleotide and extended by reverse transcriptase, yielding A3 or B3 RNA-DNA heteroduplexes. A3 lacks the 3’ adenosine (A76) due to hydrolysis of its dialdehyde derivative during deacylation and/or annealing. Extension products are analyzed by 10% denaturing PAGE (Fig. S8). In step 3b, the B5 split-tRNA fraction, with an intact 5’-ACCA76-3’ acceptor stem, is ligated to an ACCA-specific Cy3-labelled 5’-phosphorylated DNA probe (C). The probe forms a hairpin structure with a cohesive 3’-end to facilitate ligation reaction: 5’p-CGCACtt(Cy3)ttGTGCGTGGT-3’ (lower and uppercase denote loop and stem nucleotides, with the underlined cohesive end). Leveraging the split-tRNA architecture, its ligation product (B6) can be easily separated by size from ligation products involving endogenous tRNAs using denaturing PAGE (Fig. S8). This analysis workflow is based on methods from previous studies (*74*, *75*, *110*).

**Fig. S8**

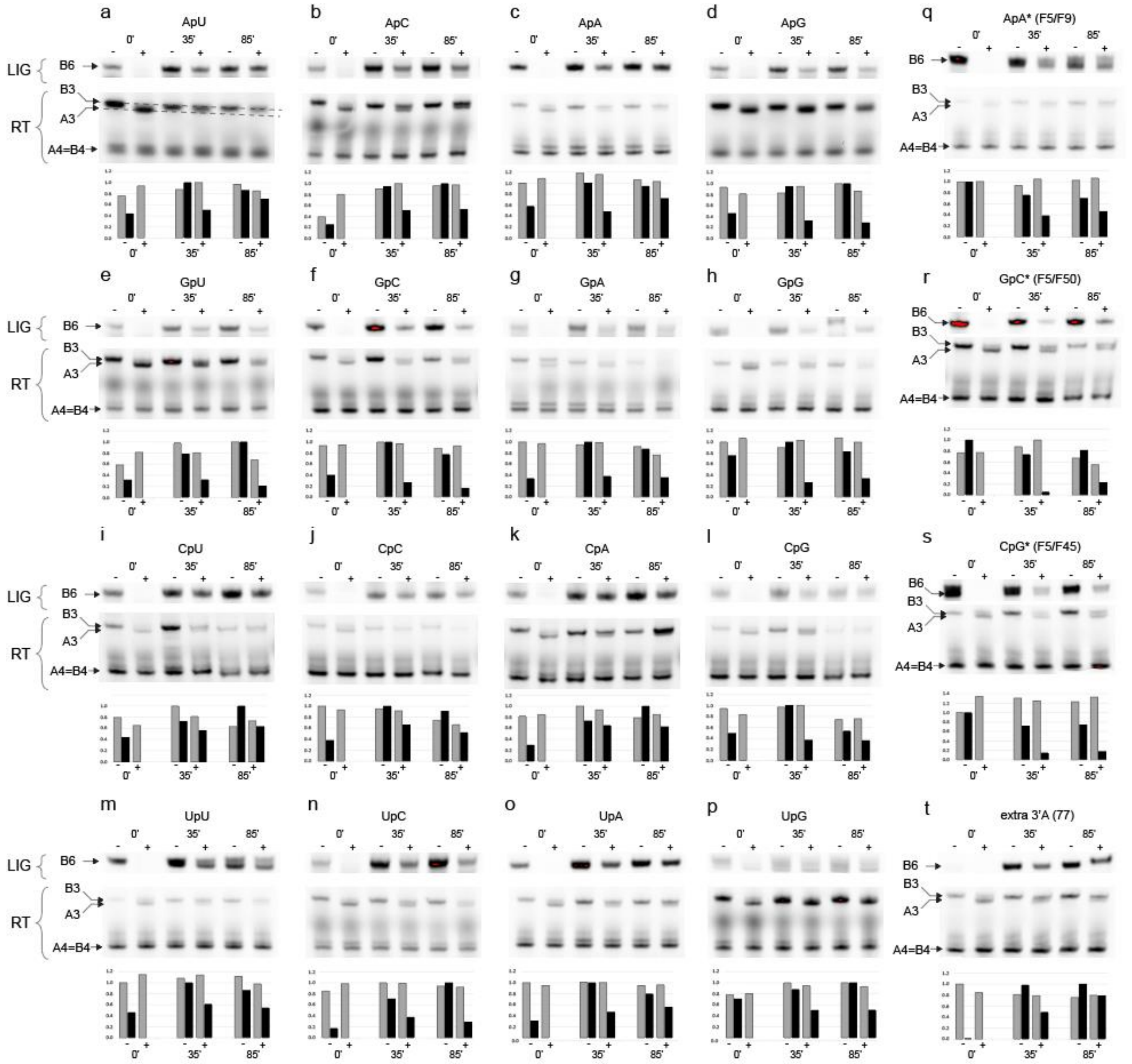


**Time course analysis of 5'-end integrity and aminoacylation status of various N37N38 split-tRNA variants during *in vitro* translation.** Fluorescence scans of 10% urea-PAGE gels loaded with products of DNA/RNA ligation or primer extension reactions of split-tRNA fragments extracted at 0, 35, and 85 minutes from *in vitro* translation reactions. RNA samples underwent the procedures outlined in Fig. S7, and the products were resolved on 10% denaturing PAAG. Each panel consists of three sections: **the top** (‘LIG’) shows fluorescent DNA/RNA ligation products dependent on split-tRNA aminoacylation (B6 in Fig. S7); **the middle** (‘RT’) shows reverse transcription products, with ‘A4=B4’ indicating a mixture of identical cDNAs from acylated (B1 in Fig. S7) and nonacylated (A1 in Fig. S7) split-tRNA, while A3 and B3 denote incompletely denatured DNA-RNA hybrids. The size difference between A3 and B3 hybrids result from the loss of 3’ adenine moiety from the aminoacceptor terminus of the nonacylated split-tRNA fraction due to periodide oxidation (Fig. S7). **The bottom** section displays a bar chart of relative densities, scaled to the maximum within each “LIG” or RT group (grey and black bars represent scaled densities of cDNA (A4 and B4) and ligation products (B6), respectively. ‘+‘ and ‘-’ indicate samples subjected or not to periodide oxidation. Panels (**a-p**) show analysis results for the 16 N37N38 split-tRNA variants indicated above each panel. These variants were assembled from a universal synthetic 1-31 fragment with a 2’-OMe modification at C31 (F5’ in Supplementary Data 2) and the respective T7-

transcribed 34-76 3'-fragment. The addition of an extra nontemplated nucleotide at the 3'-end by T7 RNA polymerase leads to a reduced amount of ligation product in the non-oxidized sample at the 0-minute time point, as the extended product cannot form an optimal complex with the hairpin-adopting DNA probe, thereby suppressing the ligation reaction. The extended terminus is efficiently repaired during translation, as shown by the significant increase in the amount of ligation product at the 35-minute time point. Panels (**q-s**) summarise the analysis results for the indicated N37N38 split-tRNA variants assembled from synthetic 5'- and 3'- F-prefixed fragments (shown in parenthesis; Supplementary data 2). Compared to the variants in panels (a-p), the 3'-terminus homogeneity allows for maximal yield of ligation product at the 0-minute time point. Panel (**t**) shows the analysis for the control split-tRNA variant assembled with a T7-transcribed 3'-fragment (34-77) containing an extra A at the aminoacceptor terminus. The absence of ligation product at time zero indicates a complete lack of substrate compatible for ligation with the DNA probe. Band densitometry was performed using ImageJ Fiji software (NIH), with integrated density values obtained for each band. Bar charts show relative densities, scaled to the maximum value for each dataset. Ligation (LIG) and RT-extension products, labeled with Cy3 and Cy5 dyes respectively, were visualized by gel fluorescence scanning at 602 nm and 700 nm using ChemDoc$^{TM}$ Imaging system (BioRad).

**Fig. S9**

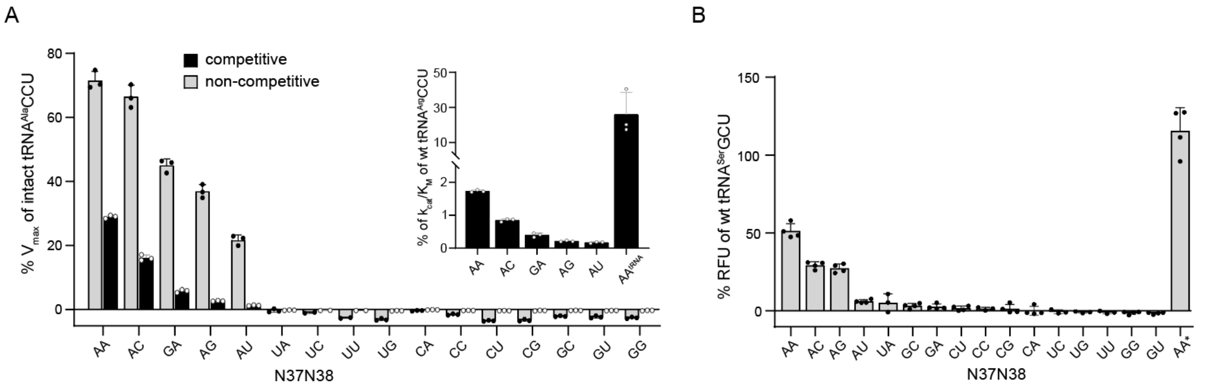


**Effect of N37N38 identity on the decoding efficiency of split-tRNA$^{Ala}$CCU and split-tRNA$^{Ser}$GCU.** (**A**) Bar chart showing the percentage of $V_{max}$ for various N37N38 split-tRNA variants relative to intact chimeric tRNA$^{Ala}$CCU under non-competitive and competitive translation conditions. Competitive conditions were recreated by supplementing lysate depleted of endogenous tRNA$^{Arg}$CCU with 0.025µM of its T7-transcribed counterpart. Split-tRNA variants were added at 17 µM. Each bar represents the mean of three independent measurements. The pronounced N37N38-dependent signal attenuation under non-competitive conditions is likely due to competition from endogenous tRNA$^{Arg}$UCU, which can also decode the AGG codon. The inset shows normalized $k_{cat}/K_m$ values for N37N38 split-tRNA variants relative to intact tRNA$^{Arg}$CCU, calculated using Eq. S6 (Supplementary text 1). The last bar indicates the normalized $k_{cat}/K_m$ value for intact chimeric tRNA$^{Ala}$CCU. Only variants with positive, non-zero values are shown. Error bars represent standard deviations. (**B**) Complementation of two consecutive AGC codons by split-tRNA variants based on wild-type tRNA$^{Ser}$GCU (Fig. S2, D to E) supplemented at 10 µM under noncompetitive conditions. The bar chart shows the percentage of maximal relative fluorescence units (RFU) for split-tRNA relative to intact tRNA$^{Ser}$GCU. The asterisk indicates the result for chimeric intact split-tRNA$^{Ala}$ grafted with a GCU anticodon. Each bar represents the mean of four independent measurements, and error bars indicate standard deviations.

**Fig. S10**

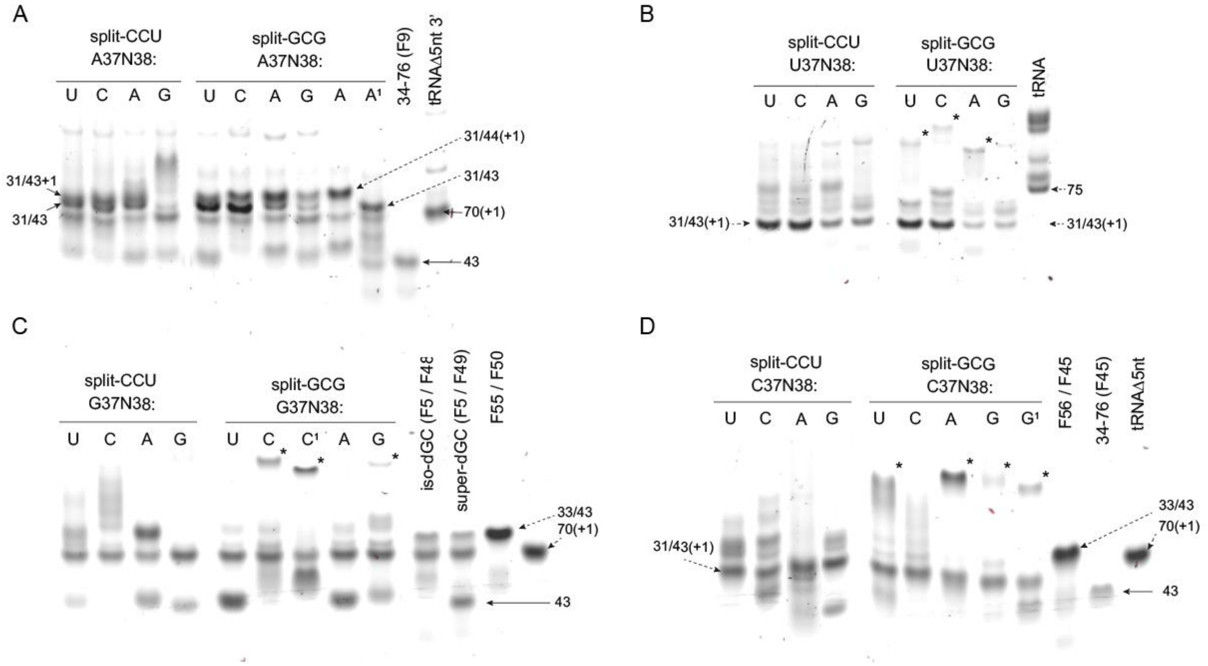


**Analysis of various N37N38 split-tRNA variants for their propensity to form stable dimers via protruding 5-mer anticodon dangling ends.** An 8% native PAGE scan was performed at a 530 nm emission wavelength of Syber Green-stained RNA samples representing the indicated N37N38 split-tRNA variants with either CCU or GCG anticodons. (**A**) Split-tRNA variants with A37N38 flanking sequences. '31/43' indicates split-tRNA products assembled from 1-31 (F5') and 34-76 fragments, with fragment sizes represented by the numbers. '31/44' represents products assembled using a 1-nt extended 3'-fragment transcribed from an extended PCR template. '+1' indicates nontemplated 3'-extensions. '34-76 (F9)' represents a synthetic 3'-fragment (supplementary data 2). 'tRNAΔ5nt' is the size marker corresponding to tRNA$^{Arg}$CCU truncated by 5nt at the 3'-end. Superscript '1' marks fully synthetic split-tRNAs assembled from universal 1-31 (F5) and 34-76 (F9) fragments. (**B**) As in (A) but for split-tRNA variants with U37N38 flanking sequences. Asterisks denote dimerization products of variable stability. (**C**) As in (A) and (B) but for split-tRNA variants with G37N38 flanking sequences. 'iso-dG' and 'super-dG' refer to split-tRNAs assembled from fully synthetic 5'- and N37-modified 3'-fragments (supplementary data 2). 'F55/F50' denotes split-tRNAs with anticodon stems extended into the G37C38 region. (**D**) As in (A-C) but for split-tRNA variants with C37N38 flanking sequences.

**Fig. S11**

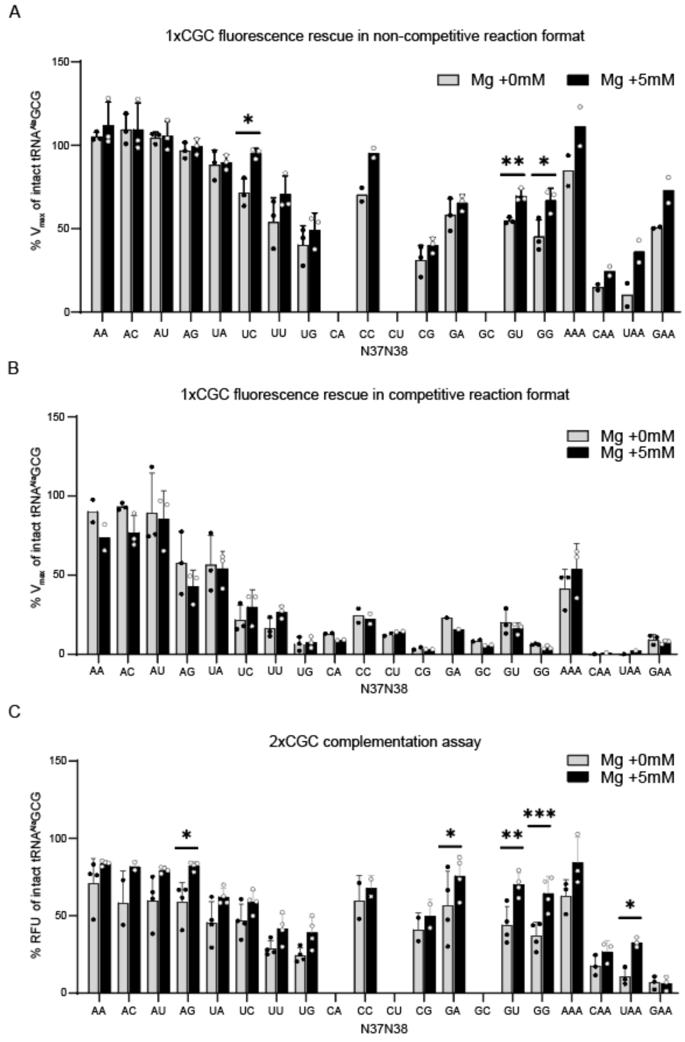


**Effect of $Mg^{2+}$ concentration on CGC codon decoding efficiency by various split-tRNAs in competitive and noncompetitive formats.** Split-tRNAs are assembled using synthetic 1-31 fragments with 2'OMe blocking groups (F5') and T7-transcribed 3'-fragments, unless noted otherwise. (**A**) Fluorescence rescue assay in a noncompetitive environment for various split-$tRNA^{Ala}$GCG N37N38 variants with 0 or 5 mM of additional $Mg^{2+}$. The Y-axis shows the percentage of maximal fluorescence accumulation velocity for split-tRNA relative to intact $tRNA^{Ala}$ with the cuGCGaa anticodon loop. An asterisk indicates the split-tRNA fully assembled from synthetic F5 and F9 fragments (supplementary data 2). (**B**) As in (A), but under competitive conditions. (**C**) As in (A) and (B), monitoring the complementation of two consecutive CGC codons by the respective split-tRNAs. The Y-axis shows the percentage of maximal relative fluorescence units, as short-term fluorescence accumulation in the negative control reaction makes $V_{max}$ less reliable.

**Fig. S12**

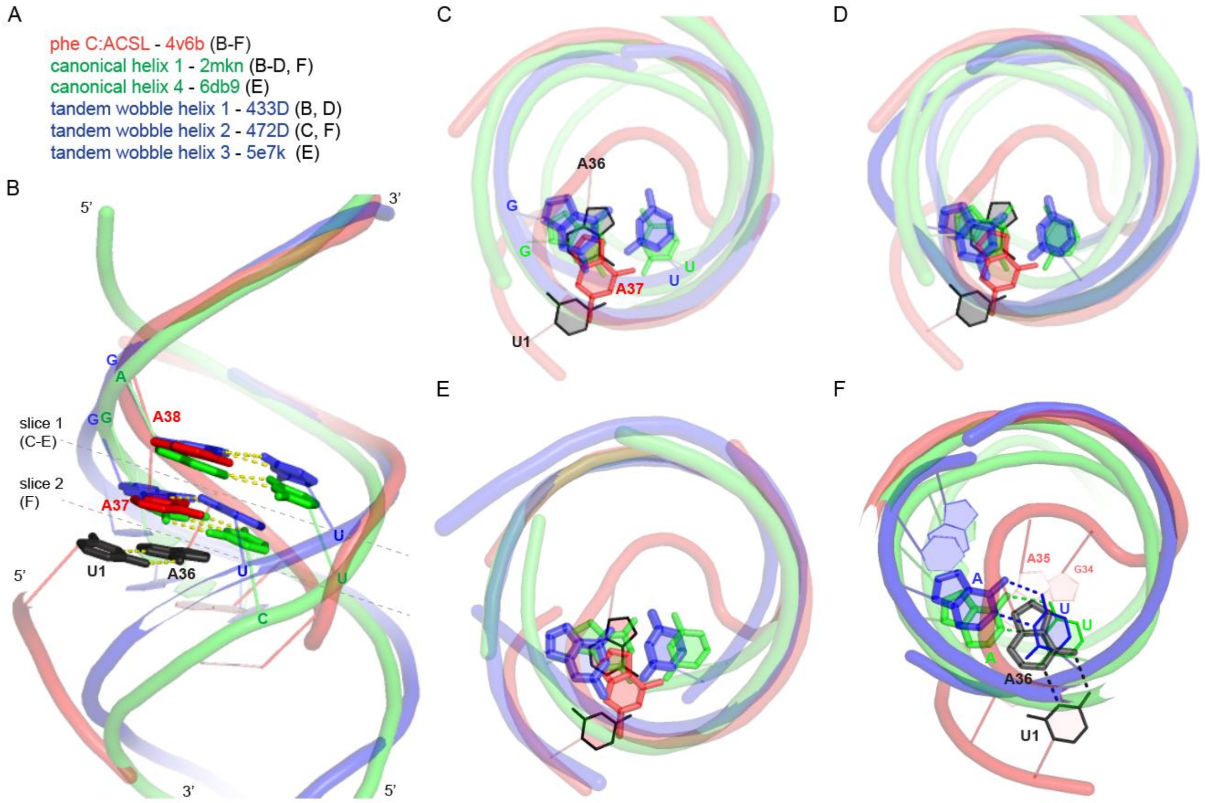


**Superimposition of canonical and wobble-containing helices onto the phenylalanine codon-anticodon stem loop structure (C:ACSL).** (**A**) List of color-coded RNA structures used in the alignments, each linked to its corresponding PDB code, with panel references indicated in parentheses. Alignment details are summarized in table S2. (**B**) Side view of the superimposition showing canonical helix 1 and tandem wobble helix 1 aligned with the C:ACSL complex formed by tRNA$^{Phe}$GAA. Backbones are shown as worms, with bases in wire representation. Canonical and wobble base pairs at positions coinciding with A37 and A38 of the ACSL are depicted as green and blue sticks, respectively, as the color code of their respective backbones. The U1:A36 codon-anticodon base pair is shown in black to distinguish it from A37 and A38 of the same C:ACSL complex. Nucleotide labels are positioned adjacent to nucleotide stems or nucleobases. Dashed lines indicate cross-sections 1 and 2 used for top-down views in the respective panels. (**C**) Top-down view of the alignment showing canonical helix 1, tandem wobble helix 2, and the C:ACSL complex at slice 1. Hydrogen bonds are omitted for clarity. (**D**) As in (C) but using tandem wobble helix 1 instead of helix 2. (**E**) As in (C) but showing canonical helix 4 and tandem wobble helix 3 corresponding to the 15-24/516-525 ds fragment derived from the *Thermus thermophilus* 23S rRNA structure (http://rna.ucsc.edu/rnacenter/images/figs/thermus_23s_2ndry.pdf). (**F**) Top-down view of the alignment shown in (C), corresponding to slice 2. Canonical base pairs from canonical and wobble helices aligned with the N1:N36 codon-anticodon base pair are represented as sticks. Hydrogen bonds are displayed for clarity.

**Fig. S13**

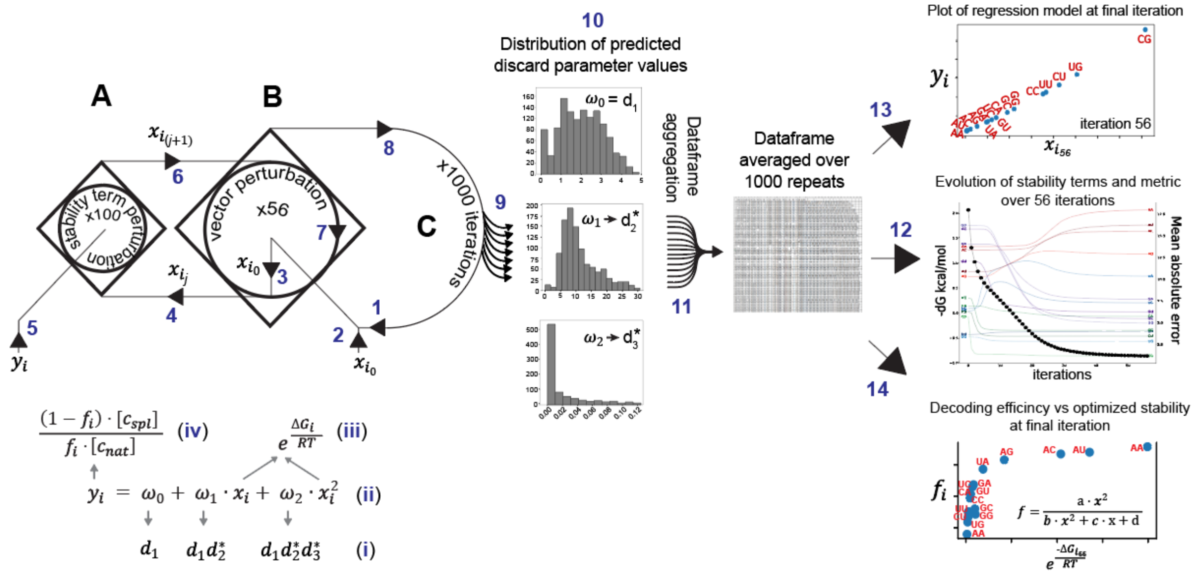


### Workflow for identifying trends in kinetic parameters of tRNA selection

Within the constraints of the kinetic equation of tRNA selection (Eq. 3), the regression model is fitted to decoding efficiencies with simultaneous iterative perturbation of initially estimated N36-N39 stability values for each N37N38 split-tRNA variant (Supplementary text 5). This process is repeated 1000 times to identify trends in the regression coefficients. Since these coefficients are proportional to discard parameters for hypothetical split-tRNA devoid of stacking interactions within N36-N39 segment (i), the observed trends provide insights into how different selection stages are influenced by the metastable codon-anticodon complex. The model is based on a system of 16 second-order polynomial equations (ii) that use a vector of independent variables, represented by stability terms $x_i$ (where $i$ refers to N37N38 variant) expressed as inverse exponentials (iii). A target vector of dependent variables $y_i$ is defined by decoding efficiency and the concentrations of both split-tRNA$^{Ala}$ and competing tRNA$^{Arg}$ (iv). The algorithm comprises three hierarchical modules (A, B, C): function **A** is executed within cycle **B** for 56 iterations, and cycle **B** is nested within the main outer iteration cycle **C**, which repeats 1000 times. In each iteration of cycle **C** (step 1), cycle **B** receives the initial vector $x_{i_0}$ (step 2), composed of estimated stability terms (see table S3), which is then passed to function A during the first iteration of cycle B (steps 3, 4).

Within function A, randomly selected stability terms are perturbed one at a time over 100 trials. After these 100 perturbations, function A selects the vector variant that best fits the constant target vector presented in step 5.

For the remaining 55 iterations, cycle B executes nested function A (steps 4-7) using an updated vector of stability terms ($x_{i_j}$) from the previous iteration. The output of cycle B (step 8) is a dataframe with columns for predicted regression coefficients, scores, and updated stability vectors from each iteration. This dataframe has 56 rows, showing the evolution of stability terms and regression coefficients as the metric improves.

The outer cycle C executes cycle B for 1000 times, resulting in 1000 independent dataframes (cumulative step 9). These dataframes are used to extract the optimized regression coefficients $\omega_0$, $\omega_1$, and $\omega_2$, plot their distributions, and identify trends in the discard parameters for hypothetical split-tRNA devoid of stacking in the N36-N39 (step 10). In step 11, all dataframes are averaged row-wise to produce a single aggregated dataframe. This averaged dataframe is used to visualize the evolution of stability terms (step 12), plot the regression model (step 13), and show the relationship between decoding efficiency and stability (step 14) for each cumulative iteration of cycle B. The plots for steps 12 and 13 are displayed for the final iteration only. The inset represents the general form of the equation describing the plot (Supplementary text 5):

$$f = \frac{k_1.[c_s] \cdot x^2}{\left(k_1 \cdot [c_s] + N \cdot [c_{wt}] \cdot (1 + \mathrm{d}_1)\right) \cdot \boldsymbol{x^2} + N \cdot [c_{wt}] \cdot d_1 d_2 \cdot \mathbf{x} + N \cdot [c_{wt}] \cdot d_1 d_2 d_3}$$

**Table S1**

**Readers of unsplit codon family boxes**

**N36 = C/G:**

| | N34 | N35 | N36 | N37 | N38 | 32:38 | $\Delta G^{dupl}$ | $\Delta G^{mod}$ | $\Delta G^{36\text{-}39}$ | $\Delta G^{total}$ |
|---|---|---|---|---|---|---|---|---|---|---|
| **Ala** | G | G | **C** | A | U | + | -7.7 | 0.00 | -1.72 | -8.42 |
| | U | G | **C** | A | C | - | -5.65 | 0.00 | -1.66 | -7.31 |
| **Arg** | C | C | **G** | G | A | - | -6.35 | 0.00 | -0.29 | -6.64 |
| | A | C | **G** | A* | A | - | -4.6 | -0.25 | -2.09 | -6.94 |
| **Gly** | C/G | C | **C** | A | A | + | -7.7 | 0.00 | -2.09 | -8.79 |
| | U | C | **C** | A | A | - | -6.15 | 0.00 | -2.09 | -8.24 |
| **Leu** | C/G | A | **G** | G | U | - | -4 | 0.00 | 0.03 | -3.97 |
| | U | A | **G** | G | U | - | -3.25 | 0.00 | 0.03 | -3.22 |
| **Pro** | C | G | **G** | G | A | + | -6.35 | 0.00 | -0.29 | -6.14 |
| | G | G | **G** | G | U | + | -7.7 | 0.00 | 0.03 | -6.67 |
| | U | G | **G** | G | A | + | -5.65 | 0.00 | -0.29 | -5.44 |
| **Val** | G | A | **C** | A | U | - | -4.3 | 0.00 | -1.72 | -6.02 |
| | U | A | **C** | A* | A | - | -3.45 | -0.25 | -2.09 | -5.79 |

**N36 = U/A:**

| | N34 | N35 | N36 | N37 | N38 | 32:38 | $\Delta G^{dupl}$ | $\Delta G^{mod}$ | $\Delta G^{36\text{-}39}$ | $\Delta G^{total}$ |
|---|---|---|---|---|---|---|---|---|---|---|
| **Ser** | C | G | **A** | A** | A | - | -4.7 | -0.75 | -2.09 | -6.54 |
| | G | G | **A** | A | A | - | -6.05 | 0.00 | -2.09 | -7.14 |
| | U | G | **A** | A** | A | - | -4 | -0.75 | -2.09 | -5.84 |
| **Thr** | C | G | **U** | A** | A | + | -4.6 | -0.75 | -2.09 | -5.94 |
| | G | G | **U** | A** | A | + | -5.95 | -0.75 | -2.09 | -7.29 |
| | U | G | **U** | A** | A | - | -3.9 | -0.75 | -2.09 | -5.74 |

**Readers of split codon family boxes**

**N36 = C/G:**

| | N34 | N35 | N36 | N37 | N38 | 32:38 | $\Delta G^{dupl}$ | $\Delta G^{mod}$ | $\Delta G^{36\text{-}39}$ | $\Delta G^{total}$ |
|---|---|---|---|---|---|---|---|---|---|---|
| **Asp** | G | U | **C** | A* | C | - | -4.4 | -0.25 | -1.66 | -6.31 |
| **Glu** | U | U | **C** | A* | C | - | -3.35 | -0.25 | -1.66 | -5.26 |
| **Gln** | C | U | **G** | A* | U | - | -3.5 | -0.25 | -1.72 | -5.47 |
| | U | U | **G** | A* | U | - | -2.85 | -0.25 | -1.72 | -4.82 |
| **His** | G | U | **G** | A* | U | - | -3.9 | -0.25 | -1.72 | -5.87 |

**N36 = U/A:**

| | N34 | N35 | N36 | N37 | N38 | 32:38 | $\Delta G^{dupl}$ | $\Delta G^{mod}$ | $\Delta G^{36\text{-}39}$ | $\Delta G^{total}$ |
|---|---|---|---|---|---|---|---|---|---|---|
| **Asn** | G | U | **U** | A** | A | - | -3.15 | -0.75 | -2.09 | -4.99 |
| **Lys** | U | U | **U** | A** | A | - | -2.1 | -0.75 | -2.09 | -3.94 |
| **Tyr** | G | U | **A** | A** | A | - | -3.45 | -0.75 | -2.09 | -5.29 |
| **Arg** | C | C | **U** | A** | A | - | -5.55 | -0.75 | -2.09 | -7.39 |
| | U | C | **U** | A** | A | - | -4 | -0.75 | -2.09 | -5.84 |
| **Ser** | G | C | **U** | A** | A | - | -5.55 | -0.75 | -2.09 | -7.39 |
| **Cys** | G | C | **A** | A** | A | + | -5.8 | -0.75 | -2.09 | -7.14 |
| **Trp** | C | C | **A** | A** | A | - | -5.8 | -0.75 | -2.09 | -7.64 |
| **Leu** | C | A | **A** | A** | A | + | -3 | -0.75 | -2.09 | -4.34 |
| | U | A | **A** | A** | A | + | -2.4 | -0.75 | -2.09 | -3.74 |
| **Phe** | G | A | **A** | A** | A | + | -3.25 | -0.75 | -2.09 | -4.59 |
| **Ile** | G | A | **U** | A | A | - | -3.45 | 0.00 | -2.09 | -4.54 |
| **Met** | C | A | **U** | A** | A | - | -3.2 | -0.75 | -2.09 | -5.04 |

**Summary of anticodon contexts in the *E.coli* tRNA repertoire and inferred stabilities of their codon-anticodon complexes.** The repertoire is divided into two groups decoding unsplit and split codon families, shown in the left and right panels, respectively. Each group is further categorized based on the G/C or A/U identity of cardinal nucleotide N36. The first five columns display the identities of anticodon nucleotides N34, N35, and N36, along with adjacent N37 and N38, which form the 3' loop flank. The "32:38" column indicates complementarity between positions 32 and 38. Gray shading highlights determinants known or inferred to reduce codon-anticodon stability. Asterisks denote modifications at N37: one asterisk for small modifications (e.g., methyl- or thio-substitutions) and two for bulky modifications. The remaining columns provide free-energy estimates in kcal/mol: $\Delta G^{dupl}$ – codon-anticodon duplex stability, averaged from Refs.(*41*, *111*, *112*); $\Delta G^{mod}$ – stabilization gained from modifications from Refs.(*21*, *113*); $\Delta G^{36\text{-}39}$ – stability contribution from the N36-N39 segment, predicted by the regression model (Fig. 4B, Fig. S13); and $\Delta G^{total}$ – total stability of the codon-anticodon complex, calculated as the sum of the other three terms, adjusted for N32:N38 complementarity. Adjustments assume a 1 kcal/mol destabilization from A32:U38 complementarity, and a 0.5 kcal/mol destabilization from U32:A38, considering that strong A37A38 stacking may mitigate distortions from the N32:N38 pairing.

**Table S2**

| Assembly used in Fig. 1 C-E: C:ACSL(phe)/C:ACSL(phe)/C:ACSL(phe)/2mnk/2az2/2jxq: | | | | |
|---|---|---|---|---|
| **Steps:** | **Structure:** | **PDB code:** | **strand:** | **Sequence:** |
| 1 | C-ACSL (ala) | 6of6 | 5'-3' / 5'-3' | CUUGCAU GGC AUGCAAG / GCCa |
| | C-ACSL (val) | 2uu9 | 5'-3' / 5'-3' | UCCCU (CMO) AC (6MZ) AGGA / GUAa |
| 2 | C-ACSL (ala) | 6of6 | 5'-3' / 5'-3' | CUUGCAU GGC AUGCAAG / GCCa |
| | C-ACSL (phe) | 4v6f | 5'-3' / 5'-3' | GGGGAUU GAA (MIA) AUCCCC / UUUu |
| 3 | C-ACSL (phe) | 4v6f | 5'-3' / 5'-3' | GGGGAUU GAA (MIA) AUCCCC / UUUu |
| | canonical A-helix1: | 2mkn | 5'-3' | GCCGUGGUCUGGUGGCCGG |
| | | | 3'-5' | CGGCACCAGACCACCGGCC |
| 4 | canonical A-helix1: | 2mkn | 5'-3' | GCCGUGGUCUGGUGGCCGG |
| | | | 3'-5' | CGGCACCAGACCACCGGCC |
| | canonical A-helix2: | 2az2 | 5'-3' | GCA (5BU) GGACGCG (5BU) CCA (5BU) GC |
| | | | 3'-5' | CG (5BU) ACC (5BU) GCGCAGG (5BU) ACG |
| | canonical A-helix3: | 2jxq | 5'-3' | GCAGAGAGCG |
| | | | 3'-5' | CGUCUCUCGC |

| Assembly used in fig. S13 A-C, E: C:ACSL(phe)/2mnk/4327D/433D | | | | |
|---|---|---|---|---|
| **Steps:** | **Structure:** | **PDB code:** | | **Sequence:** |
| 1 | C-ACSL (phe): | 4v6f | 5'-3' | GGGGAUU GAA (MIA) AUCCCC |
| | canonical A-helix1: | 2mkn | 5'-3' | GCCGUGGUCUGGUGGCCGG |
| | | | 3'-5' | CGGCACCAGACCACCGGCC |
| 2 | C-ACSL (phe): | 4v6f | 5'-3' | GGGGAUU GAA (MIA) AUCCCC |
| | wobble A-helix1: | 433D | 5'-3' | GGUAUUGCGGUACC |
| | | | 3'-5' | CCAUGGCGUUAUGG |
| 3 | C-ACSL (phe): | 4v6f | 5'-3' | GGGGAUU GAA (MIA) AUCCCC |
| | wobble A-helix2: | 472d | 5'-3' | GUGUUUAC |
| | | | 3'-5' | CACGGAUG |

| Assembly used in fig. S13D: C:ACSL(phe)/6db9/5e7k (23S rRNA 15-24:516-525) | | | | |
|---|---|---|---|---|
| **Steps:** | **Structure:** | **PDB code:** | | **Sequence:** |
| 1 | C-ACSL (phe): | 4v6f | 5'-3' | GGGGAUU GAA (MIA) AUCCCC |
| | canonical A-helix4: | 6db9 | 5'-3' | GGAUGCGCC |
| | | | 3'-5' | CCUACGCGG |
| 2 | C-ACSL (phe): | 4v6f | 5'-3' | GGGGAUU GAA (MIA) AUCCCC |
| | wobble A-helix3: | 5e7k | 5'-3'(15-24) | GGGCCCACGG |
| | | | 3'-5'(516-525) | UUCGGGUGCC |

**Detailed procedure for structural alignment.** Sub-tables are organized according to the corresponding figure panels indicated at the top, displaying one or multiple perspectives of the given alignmnet(s). The 'Steps' column specifies the sequence of superposition operations used in alignment. The 'Structures' column lists the various Codon-AntiCodon Stem Loop (C-ACSL) complexes, canonical A-helices, or A-helices containing tandem wobble base pairs (2x 5'U:G3') that overlap with the N32N33/N37N38 segment within the assembly. Nucleotide residues highlighted in green represent the anchor residues used for superposition at each step. The sequence corresponding to the 5e7k PDB code represents the helical motif (15-24/516-525) in the 23S rRNA of *T. thermophilus*, containing tandem wobble base pairs at the end of the double-stranded region.

**Table S3**

| Parameter: | Segment: | UpU | UpC | UpA | UpG | CpU | CpC | CpA | CpG | Source: |
|---|---|---|---|---|---|---|---|---|---|---|
| $\Delta G_1$ | N38/G39:C31 | -0.06 | -0.25 | -0.33 | -0.08 | -0.06 | -0.25 | -0.33 | -0.08 | Table S4 |
| $\Delta G_3$ | N37/N38 | 1.23 | 0.89 | 1.02 | 1.29 | 0.52 | 0.44 | 0.68 | 0.64 | Table S6[1] |
| $\Delta G_3$ | N37/N38 | 1.43 | 0.44 | -0.07 | 0.29 | 0.09 | 0.26 | 0.19 | -0.21 | Table S6[2] |
| $\Delta G_2$ | C1:G36/N37 | -0.73 | -0.73 | -0.73 | -0.73 | -0.49 | -0.49 | -0.49 | -0.49 | Table S5 |
| $\Delta G_2$ | A1:U36/N37 | -0.23 | -0.23 | -0.23 | -0.23 | -0.19 | -0.19 | -0.19 | -0.19 | Table S5 |
| $\Delta G_1+\Delta G_2+\Delta G_3$ | G36-N39 | 0.43 | -0.09 | -0.04 | 0.48 | -0.02 | -0.30 | -0.14 | 0.08 | total[1] |
| | | 0.54 | -0.32 | -0.59 | -0.02 | -0.24 | -0.39 | -0.38 | -0.35 | total[3] |
| $K_{eq1}*K_{eq2}*K_{eq3}$ | G36-N39 | 0.48 | 1.17 | 1.08 | 0.44 | 1.04 | 1.66 | 1.27 | 0.88 | total[1] |
| | | 0.40 | 1.72 | 2.73 | 1.04 | 1.50 | 1.93 | 1.93 | 1.82 | total[3] |
| $\Delta G_1+\Delta G_2+\Delta G_3$ | A36-N39 | 0.20 | 0.50 | 0.46 | 0.19 | 0.63 | 1.00 | 0.77 | 0.53 | total[1] |
| | | 0.17 | 0.73 | 1.16 | 0.44 | 0.90 | 1.16 | 1.16 | 1.10 | total[3] |
| $K_{eq1}*K_{eq2}*K_{eq3}$ | A36-N39 | 0.94 | 0.41 | 0.46 | 0.98 | 0.27 | 0.00 | 0.16 | 0.37 | total[1] |
| | | 1.04 | 0.19 | -0.09 | 0.48 | 0.06 | -0.09 | -0.09 | -0.05 | total[3] |

| Parameter: | Segment: | ApU | ApC | ApA | ApG | GpU | GpC | GpA | GpG | Source: |
|---|---|---|---|---|---|---|---|---|---|---|
| $\Delta G_1$ | N38/G39:C31 | -0.06 | -0.25 | -0.33 | -0.08 | -0.06 | -0.25 | -0.33 | -0.08 | Table S4 |
| $\Delta G_3$ | N37/N38 | 0.52 | 0.29 | 0.29 | 0.64 | 0.58 | 0.12 | 0.50 | na | Table S6[1] |
| $\Delta G_3$ | N37/N38 | 0.05 | 0.12 | -0.41 | -0.05 | 0.26 | -0.19 | -0.17 | -0.12 | Table S6[2] |
| $\Delta G_2$ | C1:G36/N37 | -1.29 | -1.29 | -1.29 | -1.29 | -1.55 | -1.55 | -1.55 | -1.55 | Table S5 |
| $\Delta G_2$ | A1:U36/N37 | -0.93 | -0.93 | -0.93 | -0.93 | -0.95 | -0.95 | -0.95 | -0.95 | Table S5 |
| $\Delta G_1+\Delta G_2+\Delta G_3$ | G36-N39 | -0.83 | -1.26 | -1.33 | -0.73 | -1.03 | -1.68 | -1.38 | -1.75 | total[1] |
| | | -1.07 | -1.34 | -1.68 | -1.07 | -1.19 | -1.84 | -1.72 | -1.75 | total[3] |
| $K_{eq1}*K_{eq2}*K_{eq3}$ | G36-N39 | 4.10 | 8.51 | 9.74 | 3.46 | 5.79 | 17.72 | 10.62 | 19.88 | total[1] |
| | | 6.17 | 9.84 | 17.73 | 6.25 | 7.62 | 23.02 | 18.68 | 19.95 | total[3] |
| $\Delta G_1+\Delta G_2+\Delta G_3$ | A36-N39 | 2.21 | 4.59 | 5.25 | 1.87 | 2.07 | 6.34 | 3.80 | 7.11 | total[1] |
| | | 3.33 | 5.30 | 9.56 | 3.37 | 2.73 | 8.24 | 6.69 | 7.14 | total[3] |
| $K_{eq1}*K_{eq2}*K_{eq3}$ | A36-N39 | -0.47 | -0.89 | -0.97 | -0.37 | -0.43 | -1.08 | -0.78 | -1.03 | total[1] |
| | | -0.70 | -0.98 | -1.32 | -0.71 | -0.59 | -1.24 | -1.11 | -1.15 | total[3] |

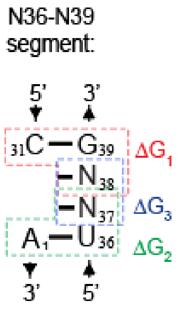


**Total free energy ($\Delta G_1+\Delta G_2+\Delta G_3$) of stacking and $K_{eq}$ for the N36-N39 segment.** Superscripts 1, 2 and 3 indicate $\Delta G_3$ values obtained from Davies 1978 (*114*), Frechet et al. 1979 (*115*), and their average, respectively. 'Total' values with superscript 1 were used as initial independent variables for modeling discard parameters (Fig. 4, B to E, Fig. S13), while those with superscript 3 were used for plotting decoding efficiencies vs. N36-N39 stability (Fig. 3A, inset).

**Table S4**

| | | 5'-pNp: | | | | |
|---|---|---|---|---|---|---|
| **Base pair:** | **T(K):** | **5'-pAp** | **5'-pCp** | **5'-pGp** | **5'-pUp** | **References:** |
| **(5')U-A** | 310 | -0,3 | -0,2 | -0,2 | -0,2 | a, c |
| **(5')G-C** | 310 | -0.2(-0.3) | -0.3(-0.2) | 0 | 0(-0.2) | a, b, d, e |
| | 293 | -0,329 | -0,25 | -0,08 | -0,06 | b, e |
| **(5')C-G** | 310 | -0,5 | -0,2 | -0,2 | -0,1 | a, b |
| **(5')A-U** | 310 | -0,3 | -0,3 | -0.4(-0.3) | -0,2 | a, c, d |

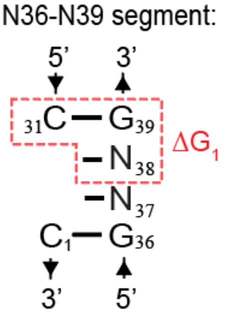


**Stacking free energy (ΔG₁) for 5'-dangling nucleotides.** Stability values at 310K for (5')G:C/pNp were derived as follows: for pAp, from Refs. a, *b*, and *e*; for pCp and pGp, from Refs. *a* and *e*; and for pUp, from Refs. *a* and *b*. Alternative values in parentheses for pAp, pCp, and pUp were derived from Refs. *b*, *d*, and *e*, respectively. ΔΔG values at 293K for (5')G:C/pNp, along with the 'temperature-independent' ΔΔH and ΔΔS, were derived from Ref. *b* for pAp and pCp and from Ref. *e* for pGp and pUp. The calculation was performed using the formula: ΔΔG = ΔΔH - 293·ΔΔS. Values used to calculate the total free energy change of stacking within N36-N39, as shown in Table S3, are highlighted. *a*(*111*), *b*(*116*), *c*(*117*), *d*(*118*), *e*(*119*)

**Table S5**

| | | 3'-pNp: | | | | |
|---|---|---|---|---|---|---|
| **base-pair:** | **T(K)** | **3'-pAp** | **3'-pCp** | **3'-pGp** | **3'-pUp** | **References:** |
| **U-A(3')** | 310 | -0.7(-0.8) | -0,5 | -0.7(-0.8) | -0,6 | a, c |
| **G-C(3')** | 310 | -1.8(-1.5) | -0.8(-1.5) | -1.75(-1.5) | -1.25(-1.0) | a, b, d, e, f |
| **C-G(3')** | 310 | -1,1 | -0,4 | -1.3(-1.2) | -0.6(-0.7) | a, b, e |
| | 293 | -1,29 | -0,49 | -1,55 | -0,73 | e |
| | 293 | -1,5 | | | -0,76 | g |
| **A-U(3')** | 310 | -0.6(-0.7) | -0,1 | -0.6(0.7) | -0,1 | a, c, d |
| | 293 | -0,93 | -0,19 | -0,95 | -0,23 | c |

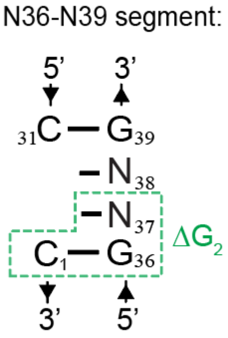


**Stacking free energy (ΔG₂) for 3'-dangling nucleotides.** Stability values at 310K for C:G(3')/pNp were obtained from Refs. *b* and *a*. Alternative values in parentheses were calculated based on parameters from Ref. *e*. Δ*ΔG* values at 293K were converted from those at 310K using van't Hoff's equation: $\ln K^{293}/K^{310} = -\frac{\Delta H}{R}\left(\frac{1}{T_2}-\frac{1}{T_1}\right)$, where $K^{293} = K^{310} \times e^{-\frac{\Delta H}{R}\left(\frac{1}{T_2}-\frac{1}{T_1}\right)}$. Δ*ΔH* values at 310K were derived from Ref. *e*. In parallel, stability values at 293K for pAp and pUp were estimated based on 'temperature-independent' parameters Δ*ΔH* and Δ*ΔS* from Ref. *g*. Calculations were performed as described in the legend for Table S1. Δ*ΔG* values at 310K were derived for pAp 3'-adjacent to A:U(3') from Refs. *c* and *d*; for pCp and pUp, from Ref. *a*; and for pGp, from Ref. *c*. Alternative values for both pAp and pGp were obtained from Ref. *a*. To estimate the ΔΔG values at 293K for A:U(3')/pNp, 'temperature-independent' parameters Δ*ΔH* and Δ*ΔS* were derived from Ref. *c*. Values used to calculate the total free energy change of stacking within N36-N39, as shown in table S3, are highlighted. *a*(*111*), *b*(*116*), *c*(*117*), *d*(*118*), *e*(*119*), *f*(*120*), *g*(*121*)

**Table S6**

| Parameter: | Method: | T(K) | UpU | UpC | UpA | UpG | CpU | CpC | CpA | CpG | Reference: |
|---|---|---|---|---|---|---|---|---|---|---|---|
| % stacked | | 293 | 0.11 | 0.18 | 0.15 | 0.10 | 0.29 | 0.32 | 0.24 | 0.25 | |
| ΔG (kcal/mol) | NMR | 293 | 1.23 | 0.89 | 1.02 | 1.29 | 0.52 | 0.44 | 0.68 | 0.64 | D |
| Keq | | 293 | 0.12 | 0.22 | 0.18 | 0.11 | 0.41 | 0.47 | 0.32 | 0.33 | |
| % stacked | | 293 | 0.08 | 0.32 | 0.53 | 0.38 | 0.46 | 0.39 | 0.42 | 0.59 | |
| ΔG (kcal/mol) | Hypo-chromicity | 293 | 1.43 | 0.44 | -0.07 | 0.29 | 0.09 | 0.26 | 0.19 | -0.21 | F |
| Keq | | 293 | 0.09 | 0.47 | 1.13 | 0.61 | 0.85 | 0.64 | 0.72 | 1.44 | |
| ΔG (kcal/mol) | MDS | 298 | -0.06 | -0.38 | -0.51 | -0.53 | -0.50 | -0.37 | -0.60 | -0.83 | B |
| ΔG (kcal/mol) | MDS | 310 | 0.10 | -0.40 | 0.10 | -0.50 | 0.60 | 0.60 | 0.10 | -0.10 | J |

| Parameter: | Method: | T(K) | ApU | ApC | ApA | ApG | GpU | GpC | GpA | GpG | Reference: |
|---|---|---|---|---|---|---|---|---|---|---|---|
| % stacked | | 293 | 0.29 | 0.38 | 0.38 | 0.25 | 0.27 | 0.45 | 0.30 | na | |
| ΔG (kcal/mol) | NMR | 293 | 0.52 | 0.29 | 0.29 | 0.64 | 0.58 | 0.12 | 0.50 | na | D |
| Keq | | 293 | 0.41 | 0.61 | 0.61 | 0.33 | 0.37 | 0.82 | 0.43 | na | |
| % stacked | | 293 | 0.48 | 0.45 | 0.67 | 0.52 | 0.39 | 0.58 | 0.57 | 0.31* | |
| ΔG (kcal/mol) | Hypo-chromicity | 293 | 0.05 | 0.12 | -0.41 | -0.05 | 0.26 | -0.19 | -0.17 | -0.12 | F |
| Keq | | 293 | 0.92 | 0.82 | 2.03 | 1.08 | 0.64 | 1.38 | 1.33 | 1.23 | |
| ΔG (kcal/mol) | MDS | 298 | -1.36 | -1.25 | -1.64 | -1.72 | -1.52 | -1.78 | -2.13 | -1.87 | B |
| ΔG (kcal/mol) | MDS | 310 | -0.70 | -0.70 | -1.10 | -1.20 | -0.60 | -0.40 | -1.40 | -1.40 | J |

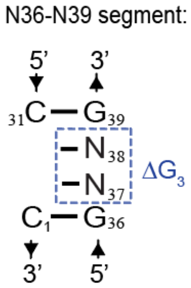


**Stacking free energy ($\Delta G_3$) for diribonucleoside monophosphates NpN.** D – Davies 1978 (*114*); F – Frechet et al. 1979 (*115*); B – Brown et al. 2015 (*122*); J – Jafilan et al. 2012 (*52*). The average of the highlighted free energy terms was used to calculate the stability of the N36-N39 segment (Table S3), which is plotted against decoding efficiencies in the inset of Fig. 3A.
